# Enhanced hexamerization of insulin via assembly pathway rerouting revealed by single particle studies

**DOI:** 10.1101/2022.04.06.487286

**Authors:** Freja Bohr, Søren S.-R. Bohr, Narendra Kumar Mishra, Nicolás Sebastian González Foutel, Henrik Dahl Pinholt, Shunliang Wu, Emilie Milan Nielsen, Min Zhang, Magnus Kjaergaard, Knud J. Jensen, Nikos S. Hatzakis

## Abstract

Insulin formulations are the hallmark of interventions for treatment of diabetes. Understanding the mechanism that governs insulin self assembly or disassembly —and the role of stabilizing additives—are essential for improving insulin formulations. We report here the real-time direct observation of single insulin self-assembly and disassembly events using single molecule fluorescence microscopy. Our direct observations revealed previously unaccounted monomeric additions to occur to all types of assemblies and allowed us to quantify the existence, abundance and kinetic characterization of diverse assembly pathways involving monomeric dimers or tetrameric insulin species. We proposed and experimentally validated a model where the insulin self-assembly pathway is rerouted favoring monomeric or oligomeric assembly events by solution concentration, additives and formulations. Our rate simulation predicted the abundance of each oligomeric species across a concentration range of 6 orders of magnitude. Besides providing fundamental new insights, the results and toolbox here can be universally applied contributing to the development of optimal insulin formulations and the deciphering of oligomerization mechanisms for other proteins.

## Introduction

Insulin is a small protein produced by the β-cells of the pancreas which is crucial for regulating the blood glucose level in all animals^1^. Since 1922, insulin has been used for treatment of patients with diabetes and research has focused on development of insulin analogs and formulations that act in either a rapid or protracted fashion^2–6^. Understanding the mechanisms underlying the self-assembly properties of insulin and its analogs are essential for interpreting how native insulin is secreted from the pancreas and tailoring the properties of therapeutic insulin for fast or protracted action^4,7^.

The commonly accepted model for insulin self-assembly is a monomer-dimer-hexamer equilibrium^8^, while monomer-dimer-tetramer-hexamer equilibrium has also been reported^9,10^. The model relies on a wealth of diverse experimental data, which all provide insights on the average oligomeric state^8^. The oligomerization data are fit with models, assuming coupling of dimer and tetramers, to yield the insulin hexamer. These models are supported by data showing insulin monomers containing two separate surfaces that are involved in the formation of dimers and hexamers^11,12^ respectively. The assembly process is shown to be further stabilized by ions and excipients. The tetramer is stabilized by Zn^2+^ and Ca^2+^ via chelating to His^B10^ and Glu^B13^, respectively, while the hexamer is stabilized by coordination of two Zn^2+^ ions to His^B10^ of three insulin dimers forming a toroidal hexamer^11,13-16^. The hexameric assembly is proposed to be further stabilized by phenolic ligands by inducing a conformational change from the T-state (tense) to the R-state (relaxed) forming the markedly more stable T_3_R_3_^17–23^.

Historically, insulin oligomerization studies have been impeded by experimental difficulties to directly observe all the individual oligomers of the assembly process and challenges in reliable recordings at biologically relevant concentrations. Current understanding of insulin self-assembly relies primarily on ensemble methodologies, correlating changes in a macroscopic property with the average oligomerization state (e.g. sedimentation equilibrium, stopped-flow, temperature jump kinetics, circular dichroism and dynamic light scattering)^8,10^. Because bulk kinetics cannot directly measure the existence of all intermediates, the community has relied on fitting the experimental observations at micromolar concentrations with models intuitively assuming, albeit not directly observing, addition of dimeric and tetrameric species. Assuming a three or four state equilibrium, reduce the number of rates from 30, if all possible transitions occur, to 4 or 6 depending on the model used. This assumption thus simplifies the analysis to allow extraction of rate constants from bulk data. The considerable challenges this simplification imposes is highlighted by the fact that, depending on experimental condition and model, the calculated hexamerisation constants (K_MD_ and K_DH_ (K_MD_: from monomer to dimer, K_DH_: dimer to hexamer) vary by ~ 2 orders of magnitude, ranging from K_MD_ = 10^3^ M^-1^ to K_MD_ = 10^5^ M^-1^ and K_DH_ = 10^8^ M^-2^ to K_DH_ = 10^9^ M^-2 8-10^. The existence of additional oligomeric forms ^24^ or oligomerization pathways involving monomeric additions, would have a profound impact on the extracted average oligomeric form, association rates and equilibrium constants and their dependence on additive and insulin formulation; all of which are crucial for the development of optimal insulin formulations.

Here, using single molecule studies, we directly observed the existence, abundance, and pathway organization of all intermediates of the self-assembly and dis-assembly process of insulin hexamers in equilibrium. We quantified the rate constants of association and dissociation that to the best of our knowledge has not been done before. While ensemble techniques require μM-mM insulin concentration, similar to those found in formulations for pharmaceutical use, single molecule techniques can directly observe phenomena in a concentration range that is similar to the pM physiological insulin concentration^25^. In this concentration regime our direct observations revealed previously unaccounted monomeric additions to occur to all types of assemblies, thus prompting for revision of existing models that consider it to be negligible. The model-free analysis offered a comprehensive kinetic characterization of all possible pairs of monomeric, dimeric and tetrameric assembly and disassembly and their dependence on excipients (Zn^2+^ and phenol). Combined with rate simulations these data elucidated the relative abundance of each of the types of oligomer species across 6 orders of magnitude and their dependence on excipients. We found hexamerization enhancement by additives to operate via re-routing the self assembly pathway favoring dimeric or tetrameric addition.

## Results

### Direct observation of individual steps of the insulin self-assembly process

We used Total Internal Reflection Fluorescence (TIRF) microscopy to directly observe the dynamic assembly and disassembly events of individual fluorescently labeled insulin monomers *en route* to the hexamer formation. We chemically attached the fluorophore ATTO655 to LysB^29^ on human insulin (in the following abbreviated HI^655^) since it is well established not to interfere with insulin self-assembly (Fig. 1AB)^26,27^ (see Supplementary Fig. 10). In a typical experiment, 10 nM of HI^655^ was allowed to equilibrate on a passivated microscopy surface (see Methods), resulting in its immobilization on the surface followed by stochastic binding and unbinding of species in solution. We acquired time-series of TIRF images with hundreds of immobilized insulin molecules present in each field-of-view. Imaging with low penetration depth allowed recordings of the dynamic assembly events on the microscopy surface with high signal to noise ratio, while particles in solution are not detected (see Fig. 1A). Using quantitative image analysis, we determined the x-y coordinates of each insulin particle with sub-pixel resolution^28–33^ (Fig. 1CD). We recorded time dependent intensity fluctuations for each particle by integrating the intensity of each diffraction limited spot. The intensity changed in a stepwise manner (Fig. 1EF) between several discrete levels, with a stochastically varying dwell time (residence time of each individual oligomeric species) and transition order. We, and others, have shown that such fluctuations directly correlate with the assembly and disassembly events for other fluorescently tagged proteins^31,34-37^.

**Figure 1:**
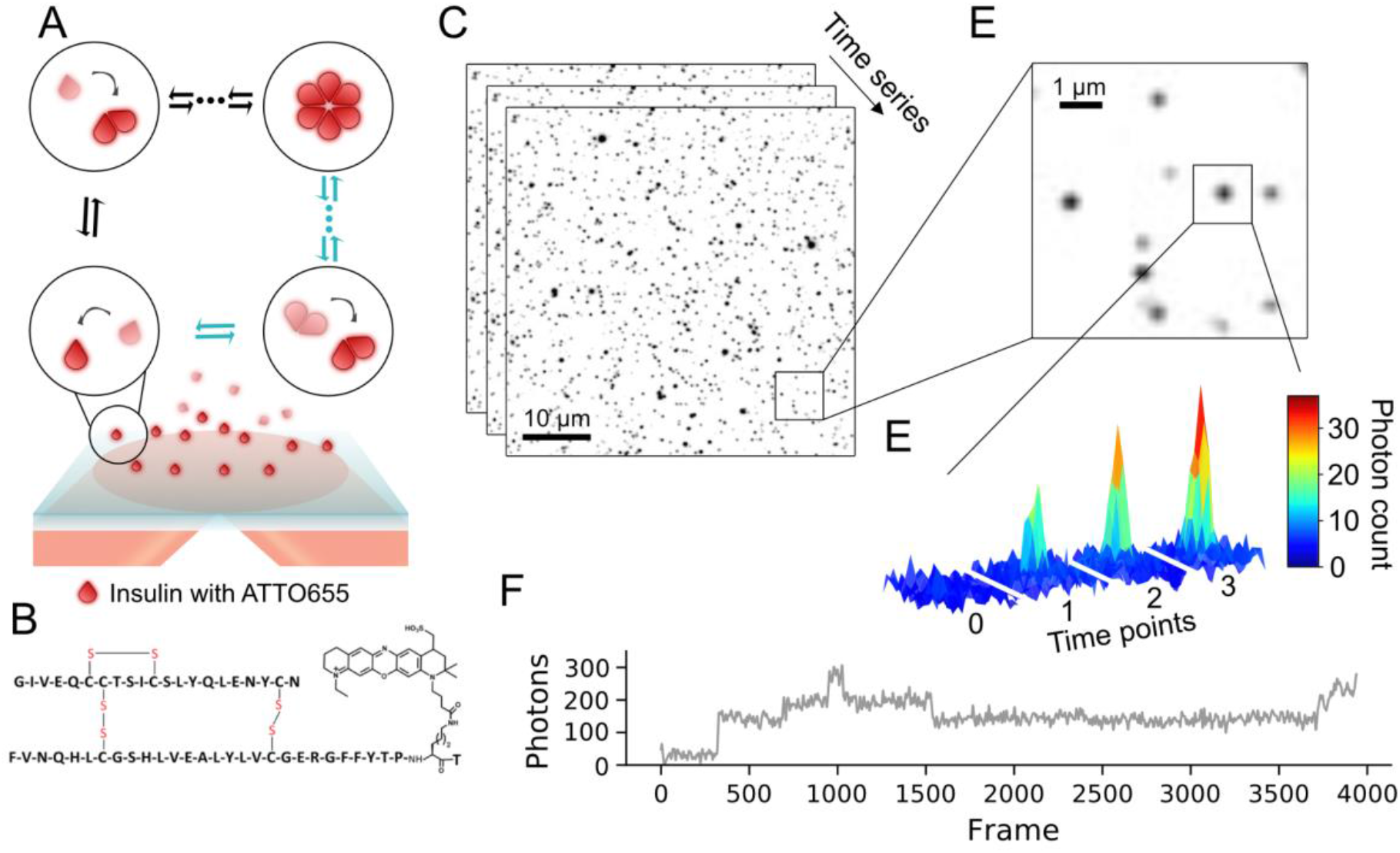
Experimental set up to observe oligomerization events of insulin using TIRF microscopy. **(A)** Representation (not to scale) of the experimental setup. ATTO655 labeled Human Insulin (HI^655^, see structure in **(B)**), are detected upon binding to the passivated microscopy surface for hexamerization. The evanescence field decays rapidly with distance from the surface and ensures that particles in solution are not detected (shaded red). The method allows to observe directly all types of oligomeric species addition. **(C)** Typical time series of micrographs recording assembly of hundreds of surface-bound HI^655^ in parallel (black spots) (scale bar 10 μm). **(D)** Close up, showing multiple insulin particles with varying intensity indicative of different oligomeric states (scale bar 1 μm). **(E)** The intensity of spots resemble the point spread function (PSF) of a diffraction limited spot suggesting they correspond to time dependent intensity variations corresponding to single oligomerization events. **(F)** Typical single molecule trajectory derived from (E) displaying discrete steps, the hallmark of single oligomerization events.

We performed several control experiments to establish that the distinct intensity shifts correspond to binding and unbinding of insulin species and the validity of our readouts. High labeling efficiency and purity (Supplementary Fig. 1-9 and Supplementary Table 1-6) minimized potential bias of kinetics from unlabeled species. Fluorophore addition did not affect kinetics or equilibrium^26^ as shown by recording of self-assembly for 1:1 mixture of HI^655^ and unmodified HI (Supplementary Fig. 14) as well as DLS measurements of unmodified HI and HI^655^ (Supplementary Fig 10). Similarly, fluorophore bleaching and blinking was quantified with immobilized monomeric biotin-labeled HI (HI^655^-Biotin) (See Supplementary Fig. 12-13 and Methods), which confirmed that photophysics do not bias our readouts. We corrected for the effects of fluorescent particles in solution, transient surface-docking, and fluctuations and unevenness of the laser excitation (Supplementary Fig. 11ABC) as described in Methods, which allowed proper calibration of the system, enabling us to confidently compare single oligomerization trajectories across images and conditions^28,29,31,38^. We directly converted diffraction limited fluorescent readouts to photons (see Methods and Supplementary Fig. 11DEF), which allowed interoperability across conditions offering future comparison to similar experiments with different experimental setups ^39^ In total, this ensured that the extracted intensity shifts corresponded to binding and unbinding of insulin monomers and dimers from solution to the immobilized insulin (see Fig. 1F and Supplementary Fig. 16). This methodology allows recordings of hundreds of individual insulin oligomerization processes and ~15,000 assembly or disassembly processes in a single experiment, providing the first real time observation of individual insulin self-assembly events at the single molecule level.

### Quantification of the abundance of oligomeric states and the kinetics of transition between them

The complexity of the self-assembly events is highlighted by the trace in Fig 2A. At 10 nM, we found each trace to stochastically sample ~20 discrete transitions (Supplementary Fig. 16+20). These events would be averaged out by conventional readouts, but are directly observed by single molecule recordings here. To classify the nature of the oligomeric species and extract the abundance and kinetics of all assembly and disassembly events we used Hidden Markov Model (HMM) analysis. We used a seven-state model representing the background and monomer through hexamer, which was fit to accurately describe transitions between all oligomeric states (See Methods for detailed description and Supplementary Fig. 15+17-19 and Supplementary Table 7). The horizontal red lines in Fig. 2A, represents a specific oligomeric state with a dwell time *τ* and vertical lines depicting transitions between oligomeric states. The residuals between trajectory (gray) and HMM prediction (red) follow a Gaussian distribution around zero and thus indicate no systematic errors, confirming the accuracy of our seven-state model (Fig. 2A, bottom and Supplementary Fig. 15C+17B+18B+19B+29B).

**Figure 2:**
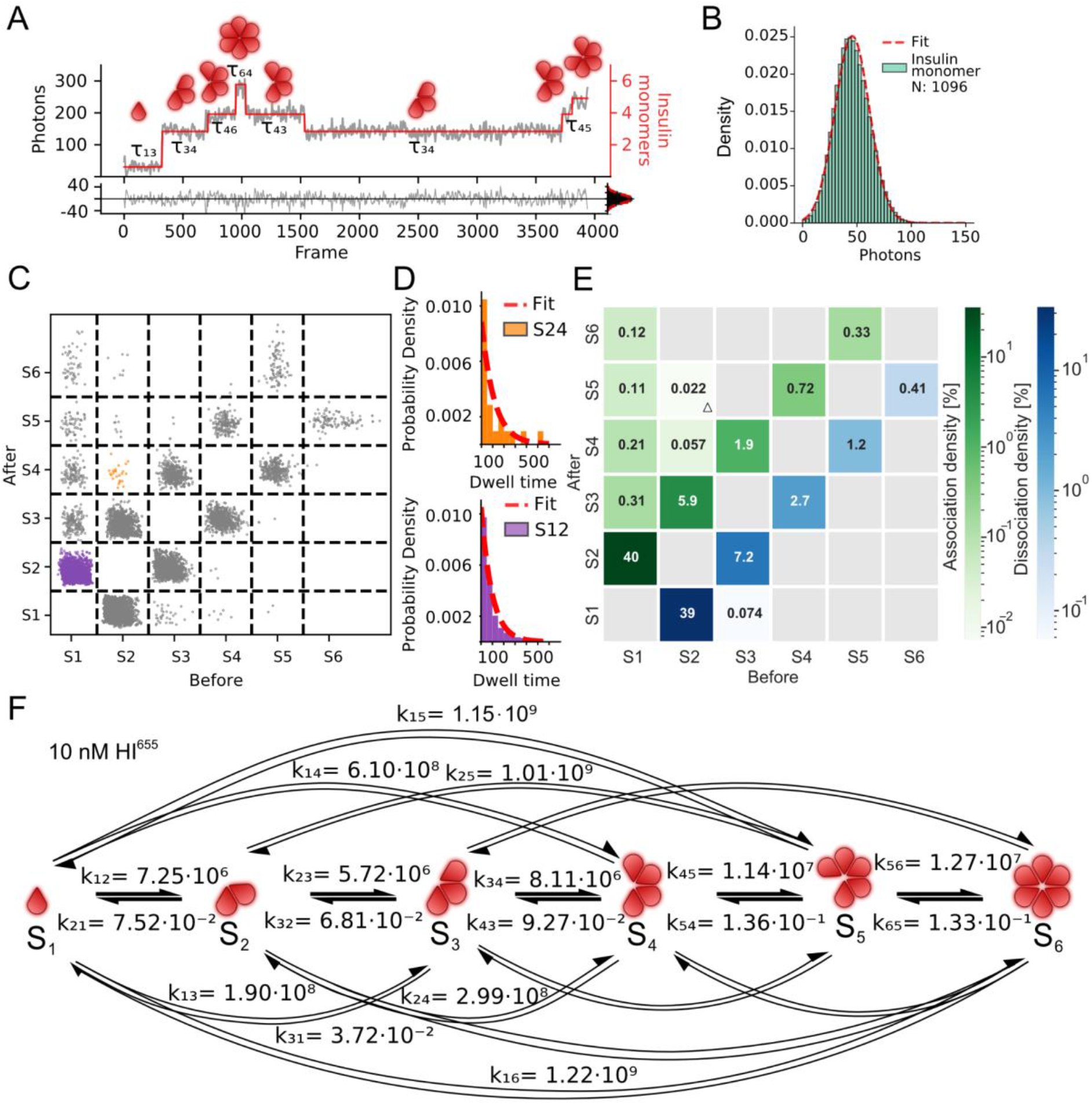
Single molecule recordings of individual insulin hexamerization events allow for extraction of kinetic parameters. **(A)** Representative trace showing discrete step-wise behavior (gray) corresponding to stochastic binding and unbinding of insulin during the hexamerization process. Hidden Markov Model (HMM) analysis using a seven-state model provide the Idealized trajectory (red) and the extraction of the corresponding dwell times *τ* for each oligomeric state. Bottom: Residuals between trajectory and idealized trajectory follow a normal distribution (μ = 0.0, σ = 14.0) confirming unbiased fitting. **(B)** Monomeric HI^655^ was imaged to calibrate the number of photons per monomer. A Gaussian fit (red line) of intensity distribution of surface passivated HI^655^ monomers revealed a mean photon value of 46 photons for a single insulin monomer (σ = 16.0) (n_videos_ = 12, N_particles_ = 1096). **(C)** Transition plot of idealized photon count found from HMM before and after a transition for 10 nM HI^655^ (n_videos_ = 4, N_particles_ = 2321, N_transitions_ = 48506). Each cluster represents a specific transition separated by a grid (-- black lines). The grid allows separation of transitions from one specific state to another. Purple denotes a transition from monomer to dimer (S1 to S2), while orange is a transition from a dimer to a tetramer (S2 to S4) **(D)** Dwell time distribution extracted from each cluster in (C) is fitted with a single exponential distribution (red dotted line) to obtain overall dwell time (τ), rate decay (that is converted to rate constant (k)) and density for that specific transition. Data shown for S12 corresponding to a transition from monomer to dimer, (N = 14501), and S24 corresponding to a transition from dimer to tetramer, (N = 21). **(E)** CHESS (Complete HEatmap of State transitionS) plot, displaying the densities of association and dissociation occupancies or each pair of possible transitions. Densities indicate that oligomerization mainly happens via monomeric addition of insulin. The numbers within the squares correspond to transition densities. Gray squares are transitions with no data points. A triangle indicates rate constants calculated using less than 10 transitions and thus a large error. **(F)** Model-free extraction of rate constants for association [M^-1^ s^-1^] and dissociation [s^-1^] with HMM analysis for 10 nM HI^655^ plotted for all observed oligomerization pathways. Notice the trend of the higher the association rate constant and the higher the dissociation constant for transitions involving higher order oligomers.

The step height of transitions showed peaks at multiples of around 50 photons (Supplementary Fig. 15B). To test whether this corresponded to addition of a single monomer, we calibrated intensity to fluorophore ratio by immobilizing biotin-labeled HI (HI^655^-Biotin) directly and indeed found the intensity to be 46 photons per labeled monomeric insulin (Fig. 2B) (see Methods). The width of the distribution is similar to the width of the residual further supporting the validity of the assay (σ = 14.0 compared to σ = 16.0, Fig. 2AB). Assembly and disassembly of insulin was found to primarily operate via association and dissociation of monomers, and to a lesser extent higher order oligomers in this concentration regime (Fig. 2B, see Methods for details and Supplementary Fig. 12+13).

For each transition between states, we extracted the start and end oligomeric state as well as the dwell time before the transition and thus the rate constant for each observed transition. The density of transitions identified by the HMM is displayed in a 2-dimensional transition plot (Fig. 2C and Supplementary Fig. 21). Each cluster of unique transitions contained up to ~15,000 transitions (see Fig. 2C and Methods), which allowed us to extract the rates for each specific transition from the distribution of dwell times. This is shown for transitions from dimer to tetramer (Fig. 2C orange and Fig. 2D top) and monomer to dimer (Fig. 2C purple and Fig. 2D, bottom, see Supplementary Fig. 22-25 and Supplementary Table 8-11 for all transitions). For the dissociation processes the unimolecular rate constant is directly extracted by the dwell time distribution. For association processes, the corresponding bimolecular rate constant can be calculated by dividing by the solution concentration of the added species (See Methods). To estimate the solution concentration of each oligomeric species in these experimental conditions we fitted the histogram of all recorded assembly transitions with 5 Gaussian distributions (see Methods and Supplementary Fig. 15D, 17C, 18C, 19C, 29C). The association and dissociation rate constants and density for each unique transition from one state (x-axis) to another (y-axis) are summarized in the CHESS (Complete HEatmap of State transitionS) (Fig. 2E, see Methods for detailed description). To the best of our knowledge this is the first detailed extraction of rates contracts for insulin hexamerization.

### Monomeric assembly and disassembly pathway for HI at nanomolar concentration in absence of additives

The direct observation of self-assembly events combined with detailed, model free analysis, allowed the extraction of the kinetic rate constants for 17 of the possible transitions involved in hexamer assembly and disassembly (Fig. 2F+3A). Consistent with earlier studies, association rate constant were found to increase for transitions involving higher order oligomeric states (e.g. k_34_ = 8.1 ± 0.3 · 10^6^ M^-1^ s^-1^ is 1.4-fold larger than k_23_ = 5.7 ± 0.1 · 10^6^ M^-1^ s^-1^) while the largest monomer addition rate constant observed for HI corresponds to a transition from pentamer to hexamer (k_56_ = 1.3 ± 0.1 · 10^7^ M^-1^ s^-1^, see Supplementary Table 12 for all data). Previously published monomer to dimer transition rates vary up to 20-fold depending on the experimental techniques used. The rate constants extracted here are a good match, though faster, to literature values^8,9^. The single particle sensitivity of the method allowed recording of monomeric additions (Fig. 2E and Supplementary Fig. 21) that would be averaged out by conventional assays focusing on dimeric additions (either monomer-> dimer -> hexamer or monomer -> dimer -> tetramer -> hexamer)^8,9,40^. Monomer assembly and disassembly represent the bulk of transitions observed (more than 99 %, Fig. 2E), in contrast to previous studies that emphasized the importance of dimer assemblies and subassemblies (Supplementary Table 12-15 and Supplementary Fig. 27B).

**Figure 3:**
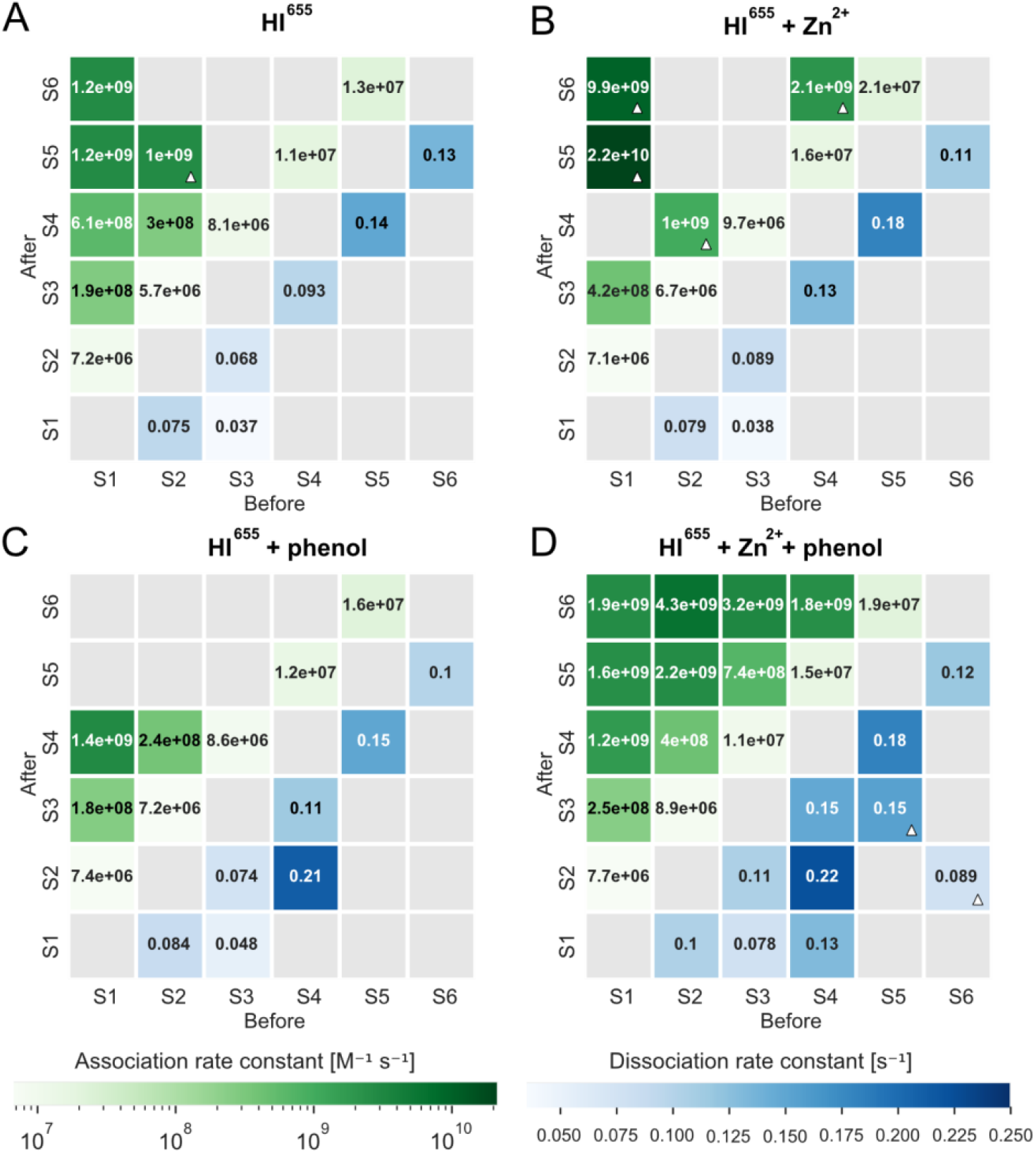
Additives stabilize addition of dimeric and tetrameric species in insulin self assembly. CHESS plots displaying the extracted association [M^-1^ s^-1^] and dissociation rate constants [s^-1^] for each specific transition for the tested conditions: **(A)** 10 nM HI^655^ (n_videos_ = 4, N_particles_ = 2321), **(B)** 10 nM HI^655^ + 100 μM Zn^2+^ (nvideos = 4, ^N^particles = 2073), **(C)** 10 nM HI^655^ + 25 μM phenol (n_videos_ = 7, N_particles_ = 3212) and **(D)** 10 nM HI^655^ + 100 μM Zn^2+^ + 25 μM phenol (n_videos_ = 5, N_particles_ = 2473). Rate constants marked with white triangle are extracted from data with less than 10 transitions. See Supplementary Table 8-11 for the number of transitions in each square.

The high sensitivity of the TIRF assay allowed the recording of the often elusive dimer to tetramer transition and furthermore so, extracting the k_on_ rate (k_24_ = 3.0 ± 0.6 · 10^8^ M^-1^ s^-1^). k_24_ is 50-fold larger than k_23_ (k_23_ = 5.7 ± 0.1 · 10^6^ M^-1^ s^-1^), which shows that transition to tetramer is kinetically favored over transition to trimers. This is consistent with the assumption underlying assembly models that emphasize dimer addition. The transition from dimer to hexamer had too low abundance (6 events out of 48,506 total transitions and 2321 trajectories, Fig. 4D, Supplementary Fig. 21A and Supplementary Table 8) in this concentration, supporting HI oligomerization to mainly operate via monomeric addition, although the monomer - dimer - tetramer - hexamer equilibrium appears to be kinetically favored.

**Figure 4:**
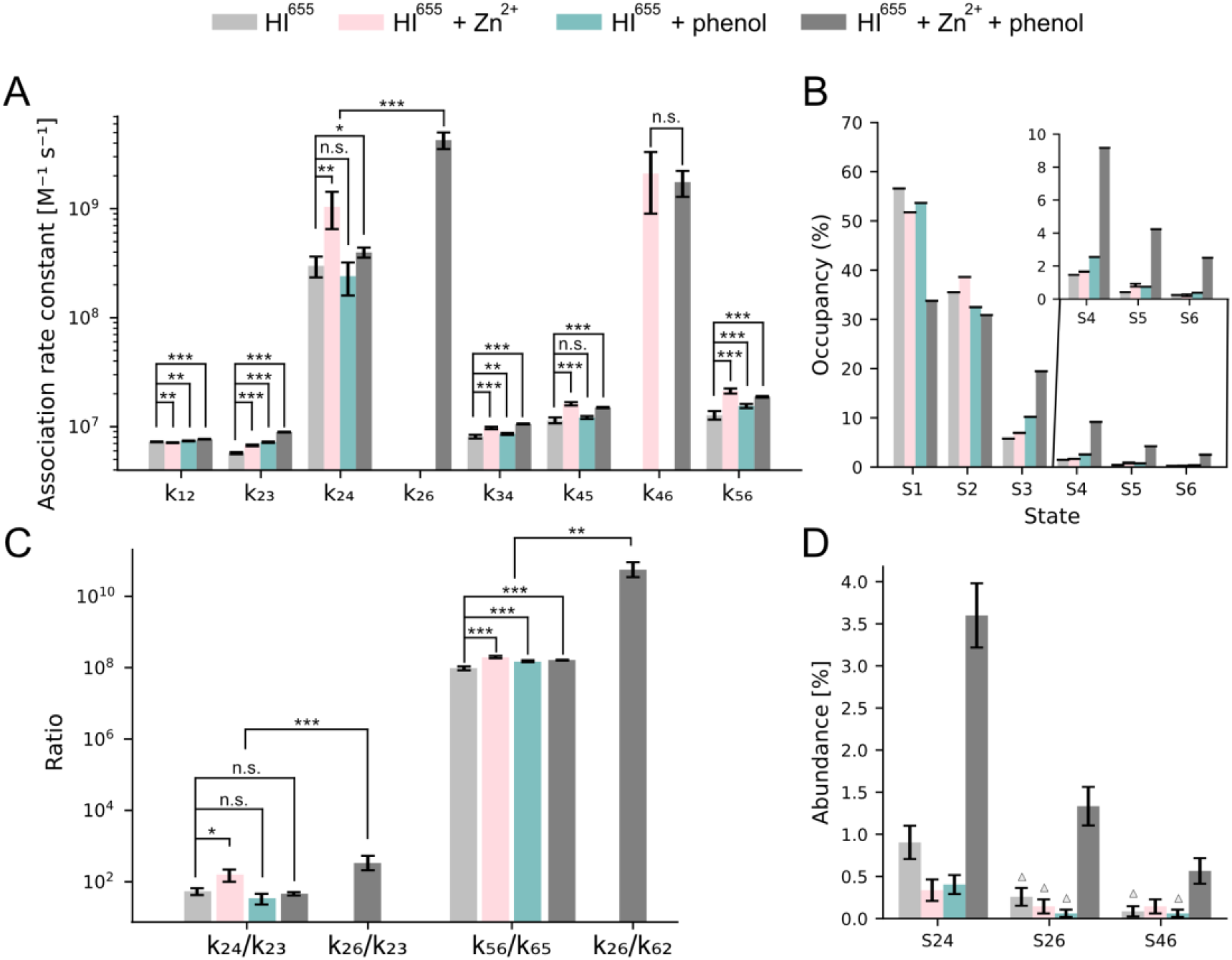
Insulin oligomerization pathway and its dependence on Zn^2+^ and phenol. **(A)** Rate constants for transitions involved in monomeric, dimeric and tetrameric addition for all experimental conditions. The rate constant of the dimer to tetramer transition is two orders of magnitude faster than monomeric addition. **(B)** State occupancies for all conditions, showing that Zn^2+^ and phenol increase the abundance of hexamers (see Methods) (**C**) Rate constant ratio between significant transitions. Rate constant ratio dimer/tetramer (k_24_) or dimer/hexamer (k_26_) and dimer/trimer transition (k_23_). The increase in ratio suggests that addition of Zn^2+^ favors addition of dimers via the tetramer over addition of monomers, while phenol has no effect. No effect on k_24_/k_23_ is observed with both phenol and Zn^2+^ suggesting that the hexamerization operates via a different route that involves addition of tetrameric insulin species from solution, namely via k_26_. The equilibrium constant (K_56_ = K_56_/k_65_) between pentamer and hexamer is increased by both phenol, Zn^2+^ and the combination of the two. The effect of Zn^2+^ and phenol is even bigger for k_26_ (k_26_/k_62_) (not observed for other conditions). **(D)** Abundance of transitions related to the dimeric additions (S24, S26 and S46) for all conditions. Triangle denotes if the transition were observed with too rarely to extract accurate rate constants. Errors are estimated as the square root of the number. N_S24_ = (21, 7, 13, 89), N_S26_ = (6, 3, 2, 33), N_S46_ = (2, 3, 2, 14), N_particles_ = (2321, 2078, 3212, 2473), n_videos_ = (4, 4, 7, 5). Error bars for rate constants are fit errors (of 4-7 measurements). Level of significance is determined by a Welch’s t-test (*p-value < 0.05; **p-value < 0.01; ***p-value < 0.001; see SI/Methods for details).

A crucial prediction from these rate constants is that the pathway of insulin assembly changes with concentration. We propose here a model where the self-assembly pathway of insulin is reliant on concentration: At nM concentrations similar to the secreted insulin concentrations used in the single molecule assays, monomer addition dominates as higher rate constant of dimer addition cannot compensate for the low population of dimers. At higher concentrations resembling pharmaceutical preparations and bulk experiments, the increasing dimer-to-monomer ratio shifts the assembly pathway towards dimer addition due to the intrinsically higher rate constant.

To test for this model we performed studies under otherwise identical conditions on the fast-acting NovoRapid insulin (Supplementary Fig. 29-30), which has been altered to exhibit reduced dimerization^3,41,42^. Consistent with current understanding the association rates involving dimer addition or higher order oligomers were reduced (e.g. k_24_ = 2.0 ± 0.3 · 10^8^ M^-1^ s^-1^ compared to k_24_ = 3.0 ± 0.6 · 10^8^ M^-1^ s^-1^ for HI^655^). Interestingly, we found NovoRapid to display increased dissociation kinetics as compared to HI^655^ (e.g. k_31_ = 0.037 ± 0.007 s^-1^ for HI^655^, and k_31_ = 0.062 ± 0.009 s^-1^ for NovoRapid) suggesting NovoRapid to have a different assembly and disassembly pathway than HI^655^.

### Zn^2+^ stabilizes the insulin hexamer and increases dimer additions

To further evaluate our model we used additives that are known to stabilize insulin hexamers. *In vivo*, such as Zn^2+^ ions, and in pharmaceutical formulations Zn^2+^ ions and phenol. Zn^2+^ is expected to stabilize the hexamer by coordinating His^B10 43–46^ in the insulin interface and expected to enhance the dimer to hexamer transition^13,14^. We quantified the effect of Zn^2+^ by addition of 100 μM Zn^2+^, an excess amount of Zn^2+^ in solution compared to HI. As expected, addition of Zn^2+^ results in an overall increase in association rate constants (See Fig. 3AB+4A and Supplementary Fig. 17+23+27 and Supplementary Table 13). The rate constant for dimer to tetramer transition increased 2.5-fold as compared to the Zn^2+^ free condition (p-value = 0.009, k_24_ = 3.0 ± 0.6 · 10^8^ M^-1^ s^-1^ to k_24_ = 1.0 ± 0.4 · 10^9^ M^-1^ s^-1^), resulting in increased likelihood of forming a tetramer from dimer, rather than a trimer (See Fig. 3AB and Fig. 4AC). Combined with the 1.2-fold increase in the rate constant of monomer addition to trimers k_34_ (k_34_ = 8.1 ± 0.3 · 10^6^ M^-1^ s^-1^ to k_34_ = 9.7 ± 0.2 · 10^6^ M^-1^ s^-1^) after addition of Zn^2+^, this results in an overall increase in tetramer density in the presence of Zn^2+^ (See Fig. 4B, Supplementary Fig. 17A+26AB and Supplementary Table 7). The overall equilibrium shift to hexamer is further compounded by the high transition rate from tetramer to hexamer (k_46_ = 2.1 ± 1.2 · 10^9^ M^-1^ s^-1^), a transition that is practically not observed in the absence of Zn^2+^ (See Fig. 3AB, and Supplementary Table 8-11). To substantiate this further we calculated the relevant equilibrium constants. K_56_ increased from 0.96 ± 0.01 · 10^8^ M^-1^ to 1.95 ± 0.15 · 10^8^ M^-1^ (p-value = 4.6 · 10^-5^, see Fig. 4C). The overall reported earlier^8,18^ 700-fold enhanced hexamerization of insulin by Zn^2+^ appears to operate via accessing the rapid tetramer to hexamer transition. Similarly the transition from dimer to hexamer is not observed (see Fig. 3B+4D and Supplementary Table 10) suggesting that in the presence of Zn^2+^ the favored route to forming a stable hexamer is via monomer - dimer - tetramer - hexamer equilibrium.

### Phenol stabilizes the insulin hexamer and has no effect on dimer additions

We then tested the effect of phenol that is also known to stabilize the hexamer, via a conformational change from T_6_ to the more stable R_6_ and is expected to do so without enhancing dimer association^44^. Indeed the addition of 25 μM phenol displayed no effect on dimer addition as compared to HI^655^ (p-value = 0.24, k_24_ = 3.0 ± 0.6 · 10^8^ M^-1^ s^-1^ to k_24_ = 2.4 ± 0.8 · 10^8^ M^-1^ s^-1^) (Fig. 3AC+4C, Supplementary Fig. 24 and Supplementary Table 14). Similarly the transition from tetramer to hexamer had a very low abundance (2 out of 60,176 transitions, Fig. 4D and Supplementary Table 9). However, phenol shifted the equilibrium constant for pentamer to hexamer transition by 55 % to K_56_ = 1.49 ± 0.09 · 10^8^ (p-value = 1.4 · 10^-5^, Fig. 3C+4C). This appears to primarily originate from reducing the hexamer dissociation to pentamer (k_65_ = 0.10 ± 0.004 s^-1^ with phenol compared to k_65_ = 0.13 ± 0.01 s^-1^ without, p-value = 0.0001) supporting the stable formation of hexamer. Besides being consistent with the proposed role of phenol stabilizing the hexamer by a conformational change from T_6_ to R_6_, these data further confirm the pathway rerouting we propose.

### Rerouting of the oligomerization pathway by the combination of Zn^2+^ and phenol additives

The combined effect of Zn^2+^ and phenol offers a remarkable increase in addition of dimeric and tetrameric species (Fig. 3D + 4C) as well as a significant equilibrium shift towards hexamer (Fig. 4B, Supplementary Fig. 26D and Supplementary Table 15). It results in 45-fold more favorable transition from dimer to tetramer as compared to transition to a trimer (p-value = 3.2 · 10^-8^, k_23_ = 8.9 ± 0.07 · 10^6^ M^-1^ s^-1^ to k_24_ = 4.0 ± 0.4 · 10^8^ M^-1^ s^-1^, Fig. 4C). Similarly, the rate constant of hexamer formation directly from a tetramer is 120 times faster (p-value = 3.3 · 10^-5^, k_45_ = 1.5 ± 0.01 · 10^7^ M^-1^ s^-1^ to k_46_ = 1.8 ± 0.5 · 10^9^ M^-1^ s^-1^). The high fidelity of the method allowed a direct observation of the transition from a dimer to a hexamer, a transition only sampled with enough statistics, when both additives are present (Fig. 4D and Supplementary Fig. 25). The community often accepts dimer to hexamer transition to operate as a tri-molecular reaction between three dimers^8,10^. Our measurements here also record directly a bimolecular reaction of addition of tetrameric species to the insulin dimer on the surface, under the assumption that concerted addition of two dimers does not occur rapidly compared to the 50 ms temporal resolution. The rate constant of transiting from dimer to hexamer is 480 times higher than transiting to a trimer (p-value = 1.3 · 10^-6^, k_26_ = 4.3 ± 0.7 · 10^9^ M^-1^ s^-1^ to k_23_ = 8.9 ± 0.07 · 10^6^ M^-1^ s^-1^, Fig. 3D + 4C). The calculated rate constant for transition from dimer to hexamer (k_26_ = 4.3 ± 0.7 · 10^9^ M^-1^ s^-1^) and tetramer to hexamer (k_46_ = 1.8 ± 0.5 · 10^9^ M^-1^ s^-1^) are similar (Fig. 3D and Supplementary Table 15) as one would expect to occur in solution supporting that the surface immobilization is not biasing the readout. The equilibrium constant of tetrameric addition is K_26_ = 4.8 ± 2.1 · 10^10^ M^-1^ is more than 100 times larger than the equilibrium constant K_56_ (Fig. 4C, p-value = 0.002). The existing monomeric and dimeric pathways will have a small contribution to the hexamerization and the overall increased hexameric form of insulin in the presence of both Zn^2+^ and phenol appears to operate via selection of an alternative, faster oligomerization pathway, that of monomer - dimer - hexamer equilibrium.

### Bridging classical ensemble studies and single molecule approach

The comprehensive extraction of multiple rate constants allowed us to simulate the time course of insulin assembly under a spectrum of 6 orders of magnitude different initial concentrations, from 1 nM to the mM range, that are not always experimentally accessible (Fig. 5 and Supplementary Fig. 32-35). We simulated the time evolution of each oligomeric species (S1 to S6) using a reaction scheme with association and dissociation of monomers, dimers and tetramers (Supplementary Fig. 31), since these transitions are dominating the experimental observations. The influence of Zn^2+^ and phenol are not explicitly modeled but are implicitly considered in the rate constants. We simulated the four different initial conditions recorded in this study (HI^655^, HI^655^ + Zn^2+^, HI^655^ + phenol and the combination of Zn^2+^ and phenol) using experimental rate constants as input (Supplementary Table 16-19). In all simulations, kinetic equilibrium was reached within ~30 seconds, where the different oligomeric species stabilize at different levels depending on the presence of additives and initial concentrations of HI (Supplementary Fig. 32-35). As our experiments do not consider the time evolution, we focus analysis on the fraction of oligomers at the end point. Extrapolation of the fraction of oligomeric species for different concentrations displays the overall sigmoidal curve reported from modeling a monomer-dimer-(tetramer)-hexamer equilibrium^8^, shifted to lower concentrations. In the absence of oligomer stabilizers, the predominant oligomeric species of insulin is the hexamer for initial concentrations above 1 μM. However, as unstabilized hexamers rapidly shed monomers and dimers, there are also considerable populations of tetramers and pentamers but also lower oligomers that are in general not accounted for by conventional analysis assuming equilibrium between three or four oligomeric species (Fig. 5A).

**Figure 5:**
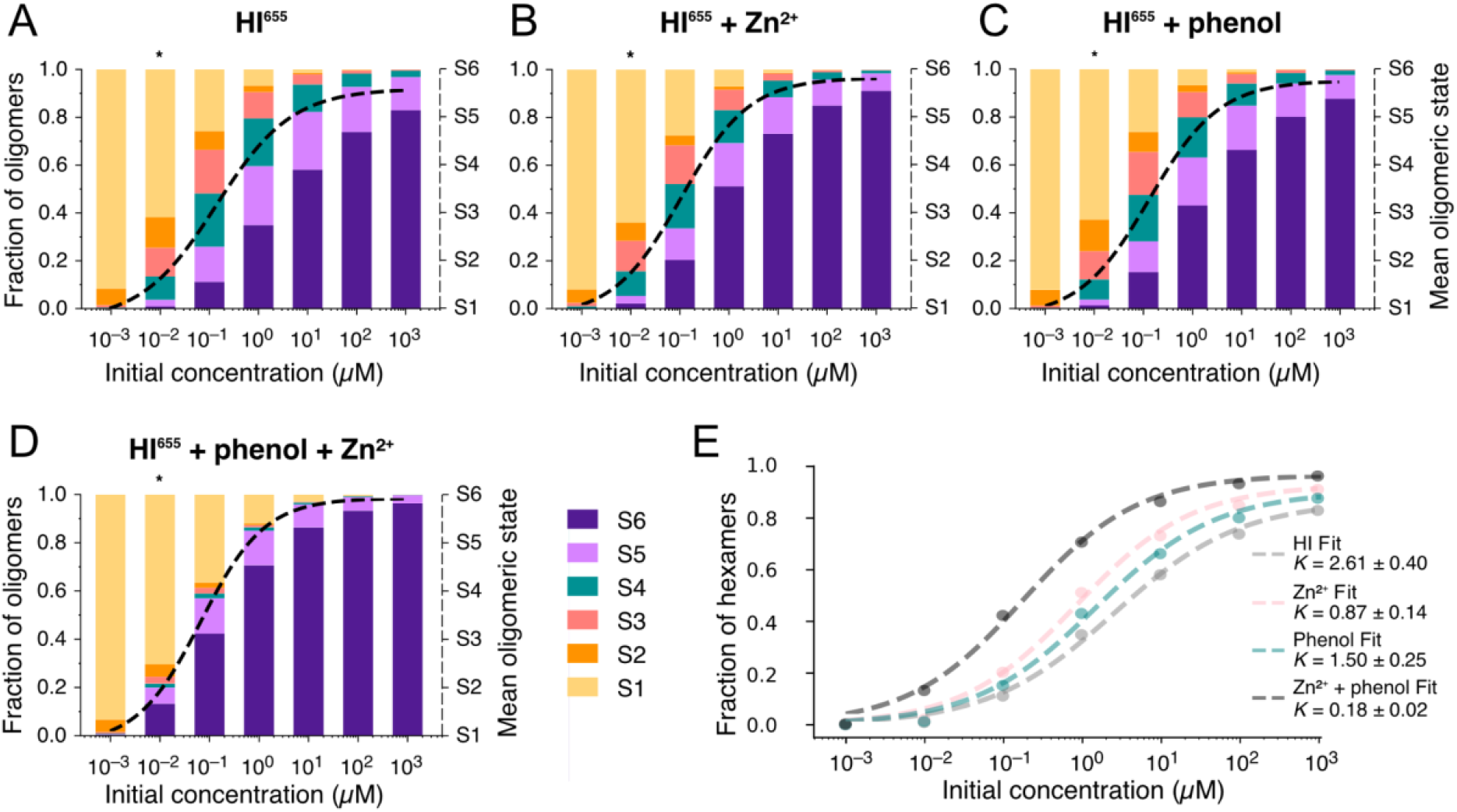
Extrapolation of the fraction of oligomerization species at varying initial insulin concentrations for each type of stabilizing additive. (A, B, C, D) Stacked plots of the fraction of oligomeric species reached at numerically simulated equilibrium for different initial HI concentrations in solution (ranging from 1 nM to 1 mM) under addition of different additives. Initial concentration represents the initial input monomer concentration before simulation has been started. (*) corresponds to the experimental conditions here of 10 nM HI^655^. Color code represents different oligomeric species. For each initial concentration, the mean oligomeric state (see Methods) is displayed as a dotted line, directly representing the read-out of an ensemble averaging readout. The method allows deconvolution of all individual species. **(E)** Fit of hexamer fraction (dark purple bar in ABCD) for each experimental condition. See Methods and Supplementary Table 20 for fitting details.

To compare with bulk experiments, we fitted the hexameric fraction increase with the Hill equation (Fig. 5E) for each experimental condition. This provides apparent affinity akin to what one would measure by bulk readouts and modeling with a three or four state equilibrium, (e.g. monomer-dimer-hexamer), but now derived from the high fidelity single molecule recordings. The extracted Hill coefficient is consistent with reports on binding on multiple sites with different affinity (apparent Hill coefficient n_h_ is < 1)^47^(see Methods and Supplementary Table 20). In the presence of stabilizers, the transition from monomer to oligomers occurs at slightly lower HI concentrations reflected by changes in the apparent hexamer affinity (see Methods). The extracted apparent affinity is lower than the one calculated by bulk readouts. Consistent with earlier studies, addition of Zn^2+^ shifts the apparent affinity by ~3-fold (from 2.6 μM to 0.87 μM, while phenol has a more moderate effect resulting in 50 % of the initial value. The combination of Zn^2+^ and phenol has a remarkable reduction of 14-fold in the apparent affinity. Our results are in general consistent with previous studies that preferentially observed hexamers in the μM to mM range^44^ and further hexamer stabilization by Zn^2+^ and phenol but provide a mechanistic insight for this stabilization and self-assembly pathway rerouting.

## Discussion and Conclusions

The widely accepted model for insulin hexamerization is monomer-dimer-hexamer, while an additional tetrameric intermediate is sometimes also considered. In both cases, the hexamer formation is promoted by complexation with two Zn^2+^ ions. These mechanisms are supported by a wide range of bulk studies with diverse experimental settings and insulin concentration in high μM range. Ensemble averaged studies provide equilibrium concentration of higher order assemblies, and extract kinetic and thermodynamic parameters based on a model assuming addition of dimers. Crucially, ensemble-based studies cannot observe these assembly events directly.

We directly observed insulin self-assembly events to be more complex than accounted for in any of the current models. We detect all oligomeric species between monomer and hexamers and observe stochastic transitions between them via monomeric and dimeric assembly and disassembly. This complexity imposes considerable challenges in current understanding, as it will be masked by conventional bulk readouts that average the behavior of a large ensemble of molecules. Indeed to the best of our knowledge to date there is no experimental data, theory or simulation, supporting the existence of sequential assembly or disassembly of insulin along the entire pathway. These results constitute the first direct validation that, contrary to current models, monomeric additions could occur to all types of oligomeric states, thus prompting for revision of existing models that intuitively assume insulin self assembly to operate via addition of dimers or tetramers.

The single molecule recordings here transformed the stochastic nature of insulin assembly from an inaccessible problem in bulk assays to an experimental asset and offered the parallelized recording of the existence, abundance and dwell time of thousands of individual assembly and disassembly events of all types of insulin oligomeric species. This resulted in the model-free extraction of 17 to 25 kinetic rate constants for these transitions, which to the best of our knowledge has not been achieved before. The directly extracted rate constants here are faster than the rates calculated by previous studies based on fitting a three or four states equilibrium^8-10,48,49^ and one cannot exclude that single molecule readout on surface immobilized molecules can have an effect. The wealth of control experiments compounded with the fact that they capture the proposed general trends, is consistent with no artifacts of the method. Notably the rate constants of some transitions involving higher order oligomers are close to what is considered the diffusion limit, where most collisions lead to binding. This is especially important for k_26_ in the presence of Zn^2+^ and phenol, further supporting that at enough concentrations a transition from dimer to hexamer is the dominant mechanism.

We proposed here a model where the self-assembly pathway of insulin is rerouted by concentration, additives and formulations (See Fig. 6) At nM concentrations relevant for insulin secretion, monomer addition dominates. At higher concentrations resembling pharmaceutical preparations, the increasing dimer-to-monomer ratio shifts the assembly pathway towards dimer addition due to the intrinsically higher rate constant. Additives such as Zn^2+^ and phenol may promote addition of dimers while the combination of Zn^2+^ and phenol reroutes the hexamerization pathway enhancing the direct transition from dimer to hexamer, a pathway rarely sampled by insulin alone. The fact that the presented method not only confirms current theory of insulin oligomerization, but simultaneously captures each individual formation step with single molecule resolution directly highlights the robustness of the analysis.

**Figure 6:**
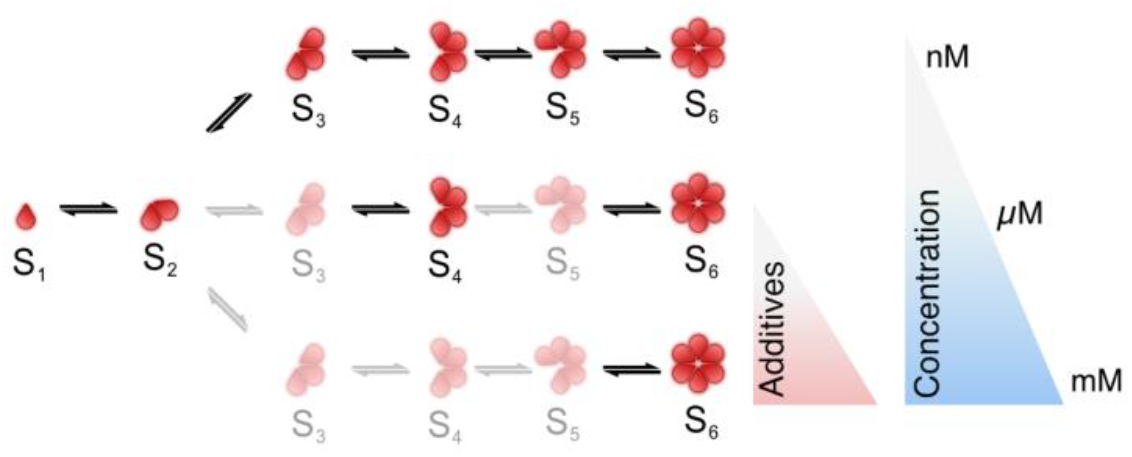
Model of pathway rerouting for insulin formulations by additives and its dependence on concentrations. Understanding mechanistics of pathway rerouting by additives and concentration enables emulation of desired conditions or behavior that may be crucial for specific pharmaceutical targets.

Insulin is believed to be secreted in hexameric form *in vivo* and to rapidly dissociate to monomers. Quantification of the dissociation of hexamers is therefore a key design parameter for pharmaceutical insulin formulations. The concentration range directly recorded here is relatively close to the insulin concentration in physiological conditions^25^. Our extraction of dissociation rates reveals dimeric dissociation to operate twice as fast as monomeric, a layer of information that is crucial for understanding secreted insulin levels in blood and how the extent of oligomerization is regulated by Zn^2+^ and phenol. The capacity of extrapolating behavior on the other hand at higher concentration for each type of formulation is crucially important for understanding and tailoring formulation for subcutaneous injection.

The experimental observations support and augment existing knowledge offering mechanistic insights on self assembly pathways rerouting as a decisive element in enhanced hexamer formation by additives. Quantitative understanding of the processes and pathways that drive association and dissociation of insulin and how they are remodeled by formulations and environmental conditions can aid both the optimized use of existing insulin formulation, the development of new novel formulations for optimized treatment of diabetes and help guide the development of glucose-responsive insulins^50^. Besides deciphering the mechanism of insulin self assembly regulation by additives, the work presented here establishes a universal methodological foundation for advancing our understanding on the regulation of the self assembly process of additional biomolecular entities.

## Materials and Methods

### Materials

All chemicals are of analytical grade and purchased from Sigma-Aldrich Denmark, unless otherwise stated. ATTO655-NHS ester was purchased from ATTO-TEC. Recombinant human insulin was purchased from Thermo Fisher USA. MilliQ water was used for aqueous preparations. The buffer used for all experiments was made from 10 mM Na_2_HPO_4_, 10 mM NaH_2_PO_4_, 5 mM NaCl, pH = 7.32.

### Insulin synthesis and labeling with chromophores (ATTO655)

#### General

High-resolution mass spectrometry was obtained on an UHPLC-MS with a QTOF Impact HD (Bruker) and Dionex UltiMate 3000 (Thermo) system equipped with a Kinetex^®^ 2.6 μm EVO C18 100 Å column (50 × 2.1 mm, Phenomenex). Purification of conjugates was done on a Biotage-Isolera HPFC 300 system with a C18 column (SNAP Ultra, C18, 30g). CH_3_CN - H_2_O (0.1 % HCOOH) was used as an eluent with a flow of 25 mL/min.

#### Synthesis of ATTO655-Human Insulin

Human insulin (21 mg, 0.0036 mmol, 3 equivalents) was suspended in 0.1 M tris buffer (0.2 mL), the pH was adjusted to 10.5 to dissolve it completely, ATTO655-NHS ester (1.0 mg, 0.00122 mmol, 1.0 equivalent) was dissolved in DMF (0.3 mL), added dropwise over 5 minutes to the stirring solution of Human insulin, and allowed the reaction mixture to stir for 15.0 min. The reaction was monitored by LCMS. Then the reaction mixture was diluted with 2.0 mL of H_2_O and pH was adjusted to pH 7.8. The product was isolated using RP-HPLC, on a Biotage SNAP ultra-column (C18, 30 g, 25 μm). CH_3_CN/H_2_O mixed with 0.1 % formic acid was used as eluents at a linear gradient of 5-50 % CH_3_CN over 20 minutes, and a flow rate of 25 mL/min. Each fraction was analyzed through LCMS. Monosubstituted products were collected separately, CH_3_CN was removed at reduced pressure using rotary evaporator, followed by lyophilized to give product as a dark green (or blackish green) powder (HI^655^-Yield: 5.6 mg, 79 %).

#### Synthesis of Di-Fmoc-Human Insulin

To the stirred solution of human insulin (300 mg, 0.0516 mmol) in tris buffer (100 mM, 4.0 mL) at pH 10.5, Fmoc-OSu (25 mg) dissolved in DMF (4.0 mL) was added, and the reaction mixture was stirred for 20 minutes at RT. LCMS analysis confirmed the completion of the reaction. The reaction mixture was diluted with water and pH of the reaction was adjusted from pH 10.5 to pH 7.8. The reaction mixture was purified on RP-HPFC (Isolera) using Biotage SNAP ultra-column C18, 60 g. The different fractions were analyzed by LCMS. The pure fractions were collected, concentrated and then lyophilized to obtain the product Gly^A1^*Fmoc*-Lys^B29^*Fmoc*-HI as a solid fluffy white powder (Gly^A1^*Fmoc*-Lys^B29^*Fmoc*-HI, Yield 139 mg: 43 %).

#### Synthesis of Di-Fmoc-Phe^B1^-Biotin-PEG_3_-HI

Gly^A1^*Fmoc*-Lys^B29^*Fmoc*-HI (135.0 mg, 0.022 mmol) was dissolved in tris buffer at pH 10.5 (100 mM, 2.5 mL) and the pH of the solution reduced to 7.2. *Biotin-PEG_3_-NHS-ester* (14.0 mg, 0.024 mmol) dissolved in DMF (2.5 mL) was added portion wise over 5 minutes, the pH was further reduced to 7.0 and the reaction mixture stirred for 40 minutes at RT. The reaction was monitored by LCMS at defined time intervals, which confirmed completion of reaction after 40 minutes. Then, the reaction mixture was diluted with water (5.0 mL) and the pH was adjusted to 7.8. The reaction mixture was purified by RP-HPFC using Biotage SNAP ultra-column (C18, 60g, 25 um), using CH_3_CN/H_2_O mixed with 0.1% formic acid with linear gradient of acetonitrile of 5-50%. Different fractions were separately analyzed by LCMS. Pure fractions were collected and freeze dried to provide Gly^A1^ *Fmoc-Lys^B29^Fmoc-Phe^B1^Biotin-PEG_3_-HI* as a solid fluffy white powder (Yield 92 mg, 62.0 %).

#### Synthesis of Phe^B1^Biotin-PEG_3_-HI

Gly^A1^*Fmoc*-Lys^B29^*Fmoc*-Phe^B1^ *Biotin-PEG_3_-HI* (90.0 mg, 0.013 mmol) was dissolved in DMSO (2.0 mL) over the stirring of 5 minute at RT. 5% Piperidine in DMF (0.3 mL) was added to the stirred solution of Gly^A1^ *Fmoc*-Lys^B29^*Fmoc*-Phe^B1^ *Biotin-PEG_4_-HI* and allowed the reaction mixture to stir for 10 minutes. Formation of product is confirmed by LCMS. Reaction mixture was further purified with RP-HPFC Isolera using Biotage SNAP ultra-column (C18, 60g, 25 um). CH_3_CN/H_2_O mixed with 0.1% formic acid were used as eluents at a linear gradient of 5-60 % CH_3_CN over 20 minutes, and a flow rate of 50 mL/min. Different fractions were separately analyzed through LCMS. Pure fractions were collected and freeze dried to get product *Phe^B1^Biotin-PEG_3_-HI* as a solid fluffy white powder (Yield 60.0 mg, 72 %).

#### Synthesis of Phe^B1^Biotin-PEG_3_-Lys^B29^Atto655-HI

To the stirred solution of Phe^B1^ *-Biotin-PEG_3_-HI* (25 mg, 0.0037 mmol) in tris buffer (100 mM, 1.5 mL) at pH 10.5, *Atto-655-NHS-ester* dissolved (1.2 mg, 0.0019 mmol) in DMF (1.5 mL) was added, and the reaction mixture was stirred for 15 minutes at RT. After 15 minutes, LCMS analysis confirmed that the reaction was completed. The reaction mixture was then diluted with water and the pH was adjusted from 10.5 to 7.8. The reaction mixture was passed through a Biotage Snap BioC4 (300 Å, 10g, C4, 20 μm) column with a flow rate of 12 mL/minute with a gradient of 5-40 % acetonitrile. Different fractions were separately analyzed by LCMS with a gradient of acetonitrile/water with 0.1% formic acid. Pure fractions were collected and freeze dried to provide *Phe^B1^-Biotin-PEG3-Lys^B29^Atto655-HI* as a dark green fluffy solid as a product (4.0 mg, Yield 31 %).

#### V8 Enzymatic analysis of Phe^B1^Biotin-PEG_3_-Lys^B29^Atto655-HI

To verify the substitution pattern, Phe^B1^*-Biotin-PEG_3_*-Lys^B29^*Atto655*-HI was subjected to enzymatic digestion by treatment with endoproteinase Glu-C from Staphylococcus. Analysis of the fragments confirmed that Atto655 was covalently attached at Lys^B29^ and Biotin-PEG_3_ substituted at Phe^B1^ position, respectively:

- Gly^A1^-Glu^A4^ (C_18_H_32_N_4_O_7_) Calculated: 416.22, Observed: 417.213
- Asn^A18^-Asn^A21^ + Ala^B14^-Glu^B21^ (C_59_H_88_N_14_O_20_S_2_), Calculated [M+2H]^2+^: 689.29, Observed: 689.27
- Arg^B22^-Lys^B29^*Atto-655*-Thr^B30^ (C_82_H_111_N_16_O_15_), Calculated [M+2H]^2+^: 813.39, Observed [M+2H]^2+^: 813.37.
- Gln^A5^-Glu^A17^ + Phe^B1^*-Biotin-PEG_3_*-Glu^B13^(C_147_H_229_N_37_O_48_S_5_) Calculated [M+2H]^2+^: 1722.271, Observed: 1722.25.

#### Synthesis of Lys^B29^Atto655-insulin aspart

Freshly purified insulin aspart (NovoRapid) (21 mg, 0.0036 mmol, 3 equivalents) was suspended in 0.1 M tris buffer (0.2 mL), and the pH was adjusted to 10.5 to dissolve it completely. ATTO-655-NHS ester (1.0 mg, 0.00122 mmol, 1.0 equivalent) was dissolved in DMF (0.3 mL) and added dropwise over 5 minutes to the stirred solution of insulin aspart, whereafter and allowed the reaction mixture to stir for 15.0 min. The reaction was monitored by LCMS^26,51,52^. Then the reaction mixture was diluted with H_2_O (2.0 mL) and the pH was adjusted to pH 7.8. The product was isolated using HPFC using a SNAP ultra-column (C18, 30 g, 25 um). CH_3_CN/H_2_O mixed with 0.1% formic acid was used as eluents with a linear gradient of 5-50 % CH_3_CN over 20 minutes, and a flow rate of 25 mL/min. Each fraction was analyzed by LCMS. Monosubstituted products were collected separately, CH_3_CN was removed at reduced pressure on a rotatory evaporator, followed by lyophilization to provide the desired product as a dark green (or blackish green) powder (Lys^B29^Atto-655-NovoRapid-HI-Yield: 5.6 mg, 79 %).

#### V8 Enzymatic analysis of Lys^B29^Atto655-insulin aspart

To verify the substitution pattern, *LysB^29^Atto655-insulin aspart* was subjected to enzymatic digestion by treatment with endoproteinase Glu-C from Staphylococcus. Analysis of the fragments confirmed that Atto655 was covalently attached to Lys^B29^:

- Gly^A1^-Glu^A4^ (C_18_H_32_N_4_O_7_) Calculated: 416.22, Observed: 417.213
- Asn^A18^-Asn^A21^ + Ala^B14^-Glu^B21^ (C_59_H_88_N_14_O_20_S_2_), Calculated [M+2H]^2+^: 689.29, Observed: 689.27
- Arg^B22^-Lys^B29^*Atto-655*-Thr^B30^ (C_80_H_107_N_16_O_20_S), Calculated [M+2H]^2+^: 822.38, Observed: [M+2H]^2+^: 822.36.
- Gln^A5^-Glu^A17^ + Phe^B1^-Glu^B13^(C_126_H_197_N_34_O_41_S_4_), Calculated [M+3H]^3+^: 990.442, Observed: 990.422, Calculated [M+4H]^4+^: 743.084, Observed: 743.084.

#### Sample preparation

Degassed MilliQ H_2_O (1 mL) was added to HI^655^, HI^655^-Biotin and NovoRapid^655^ (1.0 mg of dry powder) and shaken gently for 2-3 minute, at which time it appeared as a suspension. The pH of solution was increased to pH 10.0 by addition of 0.5 M NaOH (added 1-2 μL for 2-3 times to reach the pH of 10.0 to dissolve it completely as a transparent solution. Further pH was lowered with 0.5 M HCl and shaken gently so that cloudy insulin dissolved, and the solution appeared transparent. Afterwards, the pH of the solution was adjusted 7.4-7.5 using 0.2 M HCl. The solution was further filtered into another Eppendorf tube through 0.2 μm syringe filters in order to remove any precipitate/aggregates. The concentration of insulin was further determined on a Nanodrop instrument. The molar absorption coefficient value for Atto655 is 1.25 · 10^5^ M^-1^cm^-1^. After determination of concentration, stock solutions were utilized for further experiments.

### Dynamic Light Scattering

DLS measurement was carried out on a Malvern Zetasizer (Malvern, United Kingdom) μV instrument at 25 °C using a 2 microlitre Quartz cuvette with 1.25 mm light path length. Hydrodynamic radius was calculated using a standard equation with dynamic viscosity of water at 25 °C which is embedded in Malvern program. Sample was measured at varying concentrations of 50 and 40 μM insulin in 10 mM Na_2_HPO_4_, 10 mM Na_2_HPO_4_, 10 mM NaH_2_PO4, and 5 mM NaCl at pH 7.5. The mean hydrodynamic radius for each condition was found with a log-normal fit to the data.

### Microscopy surface preparation and surface passivation

Microscope coverslips were cleaned thoroughly by sonication in 3 x 2 % MilliQ Hellmanex solution, 3 x MilliQ and 1 x methanol for 15 min each. In between each round of sonication, the coverslips were rinsed with MilliQ. The clean coverslips were stored in methanol solution until usage. Clean coverslips were dried under a nitrogen flow and plasma cleaned for at least 4 minutes before attaching a sticky slide to it. Surfaces were passivated with 80 μL 100:1 mixture of 1 g/L PLL-PEG / PLL-PEG-biotin that incubated for at least 2 hours. After incubation, the wells were thoroughly rinsed with MilliQ and functionalized with 80 μL 0.1 g/L neutravidin as described previously^28,37^. After functionalization the wells were again thoroughly rinsed and the surfaces were stored in MilliQ until insulin addition.

### Total Internal Reflection Fluorescence microscopy (TIRFm) imaging

Before imaging a video of an empty surface with only buffer was acquired for subsequent background correction. 350 μL insulin solution (10 nM HI^655^) in buffer was flushed into the chamber, and imaging was started immediately. All experiments were carried out in buffer at room temperature with Zn^2+^ and phenol in excess amounts (100 and 25 μM). Data from at least 4 measurements were combined in all figures.

For HI^655^-Biotin control experiment, 350 μL 1 pM insulin was flushed into the chamber and immediately flushed extensively with buffer to remove any non-bound insulin. A total of 12 videos at different positions on the surface were recorded in a single chamber.

All experiments were conducted at a Total Internal Reflection Fluorescence microscope (TIRFm, IX 83, Olympus) equipped with one EMCCD camera (Hamamatsu) and an oil immersion 100x objective (UAPON 100XOTIRF, Olympus). ATTO655 fluorophores were excited using a 640 nm solid state laser line. Imaging was performed with 10 % laser power (200 μW), 50 ms exposure time (followed 100 ms waiting time between frames resulting in a frame rate of 6.7 s^-1^), 100 nm penetration depth and 300 EM gain. Imaging was done for a total of 4000 frames per video, resulting in a total imaging time of approximately 10 minutes.

### Image analysis

Quantitative image analysis was performed using an in-house software based on previous publications^28,29,53,54^ and outlined in the following paragraphs. Traces were sorted based on different criteria, e.g., signal/background ratio, noisiness and whether the traces displayed a clear stepwise behavior.

#### EMCCD calibration

Prior to the addition of particles, a control movie—with identical acquisition parameters to the subsequent measurement—was recorded on an empty sample for camera calibration (Supplementary Fig. 11D, Supplementary Fig. 11E shows a movie with particles for comparison). The calibration allowed conversion of intensity values into photons counts for subsequent measurements.

400 random pixels were selected from the control movie, forming 400 histograms of intensity values for each. Assuming a constant mean pixel value for such arrays, fluctuations are expected to arise only through EMCCD measurement noise. The noise is well described by the convolution of a Poisson distribution and an Erlang distribution. With the Poisson distribution modeling effects of shot noise and the Erlang distribution modeling the birth-death processes in the EM-gain. The resulting three-parameter distribution for pixel intensity is

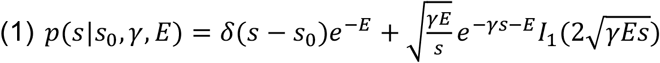

where *s* is the pixel intensity and *I*_1_ is the modified Bessel function of the first kind. The three parameters are *s*_0_, a factory-set offset to the observed camera intensity, *γ*, the inverse gain of the camera, and *E*, the expected photon count from the pixel array ^39,55,56^. Each of the 400 pixel arrays were fit with equation 1 using a chi-2 fit (Supplementary Fig. 11F). The average of all parameters with a p-value greater than 1% were then used to estimate the mean photon counts in pixels of subsequently acquired movies. This was done by simply offsetting and scaling the observed intensities

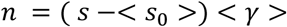

#### Illumination profile correction

To confidently compare photon counts across the images with fluorescent particles, a correction from the Gaussian illumination profile had to be performed (Supplementary Fig. 11ABC). The time-average of each movie was convolved with a Gaussian (*σ* = 30 pixels) to estimate the background illumination profile (Supplementary Fig. 11B). Each movie was then divided with its estimated background illumination profile to get the corrected set of movies (Supplementary Fig. 11C). Finally, to improve image contrast for subsequent particle identification, the movies were convolved with a Gaussian along the time axis (*σ* = 3 frames).

#### Particle localization and signal extraction

Localization of each particle was performed on an average representation of the time series since the particles are not moving. The x-y position of each fluorescent particle was determined by locating Gaussian shaped peaks of fluorescence with a certain size as described previously^28,29,53,54^. The specific parameters used for localizing particles in Trackpy were set as diameter = 11 pixels, sep = 6 pixels while minmass is calculated as *minmass = meanavgvideo · 0.4*, where meanavgvideo is the mean average pixel value of the Gaussian and illumination corrected movie.

Subsequently, the exact position is refined using a 2D Gaussian fit to the PSF. The baseline (called *b*) for the 2D Gaussian fit was used for background correction. After localization, the signal was integrated with a roi (region of interest) of 9 pixels in diameter in all 4000 frames and lastly corrected for local background variations by subtracting the background value found from Gaussian fitting to obtain a single background corrected trajectory as:

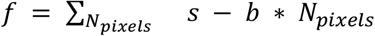

where *s* is the pixel intensity, *b* is the baseline from 2D Gaussian fit and *N_Pixels_* represents the number of pixels within a roi (region of interest).

#### Intensity calibration to number of insulin monomers

Visual inspection of the trajectory displayed in Fig. 2A suggests that insulin assembly and disassembly events primarily result in a signal change in integers of ~50 photons. To validate this, we directly converted the diffraction limited fluorescent readouts to photons from 12 experiments with a surface-passivated biotinylated monomeric insulin (HI^655^-Biotin, N = 1096 particles) (Fig. 2B). Trajectories arising from aggregates were discarded. The methodology allows interoperability across experimental conditions and setups^39^. Analysis revealed an excellent agreement between both visual inspection, model prediction and Gaussian fit (μ = 46 photons, σ = 16 photons) to monomeric insulin (Supplementary Fig. 12AB). We excluded artifactual monomeric addition events by ensuring minimal bias from the surrounding via a *roundness* parameter allowing us to exclude incidents where particles were localized in the same pixel. Similar analysis was performed under addition of phenol, to ensure minimum bias from phenol on fluorescent readout (N = 781). As expected, no effect was found (μ = 41 photons, σ = 14 photons) (Supplementary Fig. 13A).

### Bleaching and blinking control experiment

Experiments with HI^655^-Biotin were used to quantify bleaching and blinking. Trajectories arising from aggregates were discarded. Bleaching analysis was made in increments of 400 frames. If the mean photon count for a specific particle was equal to or less than 30 photons in an increment, it was denoted as bleached. Blinking analysis was performed on trajectories from frame 100 to frame 1000 to remove any bias from bleaching. If the residual (photon count in a specific frame) was bigger than or equal to 3 standard deviations of the trajectory it was denoted as a dark state, i.e. the fluorophore is blinking (Supplementary Fig. 12CDE). Similar analysis was performed under addition of phenol, to ensure minimum bias from phenol on fluorescent readout. As expected, no effect was found (Supplementary Fig. 13C).

### Insulin oligomerization experiment

Experiments with HI^655^ nonspecifically binding to the surface were used to observe and quantify single oligomerization events. The traces for the same conditions (insulin-, Zn^2+^ and phenol concentration) from different surfaces were merged for later use. The summed photon histogram for all trajectories followed a Gaussian mixture model of seven Gaussian consistent with a seven-state model accounting for the six steps of hexamerization plus background (essentially no particles observed) (Supplementary Fig. 15A). Thorough investigation into the fit revealed that the best seven distributions were found to be as listed in table 1 and were applied to the HMM analysis. A hard threshold for photon count was set to be 500 photons, and traces containing higher photon count than this were discarded before the HMM analysis as they were higher order aggregates.

**Table 1:** Hidden Markov Model input as initial guesses

| Distribution: | S0 | S1 | S2 | S3 | S4 | S5 | S6 |
| --- | --- | --- | --- | --- | --- | --- | --- |
| Input [photons]: | 20±25 | 50±25 | 100±35 | 150±35 | 200±35 | 250±35 | 300±35 |

For 10 nM NovoRapid^655^ the signal to noise was lower than for experiments with HI^655^. For this reason, we used Gaussian smoothing with a width of σ = 5, on the raw trajectories for 10 nM NovoRapid^655^ to obtain comparable signal to noise (Supplementary Fig. 15C and Supplementary Fig 29B).

### Hidden Markov Model analysis

The HMM data analysis was implemented using the Pomegranate package for Python similar to published methodologies^28,31,57^. Due to the low occupancy of the higher order oligomers of the insulin (tetramer, pentamer and hexamer, Supplementary Table 7), the state values had to be frozen in order for the HMM to fit the entire dataset. For each trace, a model with 7 states (as defined in Table 1) was fitted. The idealized photon histogram was fitted with a mixture of 7 Gaussians to extract state occupancies (Supplementary Fig. 15B, 17A, 18A, 19A, 29A). To determine the error, we employed a parametric bootstrapping approach, where the fit was reinitiated 10 times to generate a bootstrap distribution of fit parameters estimates. From each parameter distribution, the mean and error was determined.

Although fluorescent intensity values are not technically Gaussian distributed, it has in practice been shown to be a robust method with little discrepancy when the values are distributed far away from 0 due to central limit theorem. Also, the HMM fit was evaluated by plotting the residuals that showed no systematic error of HMM fit for all conditions (Supplementary Fig. 15C, 17B, 18B, 19B, 29B).

### Determination of transition rate constants

The idealized traces were further investigated by plotting all transitions in a Transition Density Plot as this is the current established method. Extraction of specific transitions is classically performed using fitting (e.g. k means clustering ^28,31^) for simpler systems with fewer transitions, to reliably identify up to 64 clusters in total for all conditions. However, this method was not optimal for this data due to the many transitions possible and low sampling of higher order transitions/states. This is despite very thorough examination of the data and extensive optimization of initial guesses for x-y positions of each cluster.

Here we applied a series of thresholds that can be subjected to extensive analysis to extract kinetic and thermodynamic insights. The grid thresholds were made in agreement with the HMM input, so that a specific state or transition found by the HMM, would be correctly separated by the grid. This allowed for a total theoretical number of 42 separable clusters (excluding those involved with S0). The TDP plots showcase insulin association and dissociation at 10 nM concentration regime to operate primarily from monomeric additions and to lesser extent via dimer or higher order oligomer addition (Supplementary Fig. 21 and Supplementary Table 8-11).

Transition rate constants were found by fitting the dwell times contained in every cluster with a single exponential decay using a maximum likelihood fitting scheme (Supplementary Fig. 22-25 and Supplementary Fig. 30). Dwell times above 75 s were not fitted to avoid long-lived outliers that do not follow a single-exponential decay (less than 5% of observable dwell times) ^58^. The occupancy for each transition represents how many times the transition was observed.

Because dissociation of oligomers is unimolecular and association a bi-molecular reaction and thus oligomer concentration dependent, the decay rate for association was converted to rate constant by dividing with oligomer concentration in solution. Solution concentration can ideally be measured by the density of each oligomer landing directly on the surface. In order to get statistical significance, we estimated the oligomer concentration in solution by fitting the histogram of assembly transitions with 5 Gaussian distributions (representing monomer, dimer, trimer, tetramer and pentamer addition). In short, the histogram revealed 5 distributions corresponding to monomer, dimer, trimer, tetramer and pentamer addition, respectively, and the weight from each distribution denotes the percentage of free soluble oligomers (See Supplementary Fig. 15D, 17C, 18C, 19C, 29C).

The found rate constant for association and dissociation can be found in Supplementary Table 12-15, Supplementary Fig. 22-25 and Supplementary Fig. 30).

### CHESS plot

We constructed CHESS (Complete HEatmap of State transitionS) for a transparent and convenient way of inspecting the multidimensional kinetic and thermodynamic information (Fig. 2E, Fig. 3, Supplementary Fig. 26). The x-axis shows the state before a transition, while the y-axis shows the state after a transition. Each square denotes one transition, the kinetic (rate constant) and/or thermodynamic (transition density) parameter written as the number in the square. The color code of each square represents the value written inside. Shown in Fig. 2E with transition densities for association and dissociation for all quantified transitions for 10 nM HI^655^.

### Calculation of free energies

Free energies were calculated based on transition state theory, where the monomer state was considered a ground state with relative energy 0 kJ/mol ^58^. The relative free energy difference between states are

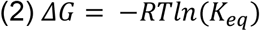

where R is the gas constant and T = 298 K. Equilibrium constant *K_eq_* is given by

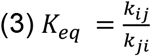

where *k_ij_* and *k_ji_* are transition rates between two states *S_i_* and *S_j_*. The Gibbs energy of activation (energy barrier) for each transition is given as:‡

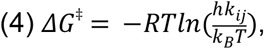

### Statistical analysis

All values reported are the average at minimum of four independent replicates from separate experiments. Error bars represent SD as indicated in the corresponding figure legends, unless otherwise stated.

Level of significance is determined by a Welch’s t-test (*p-value < 0.05; **p-value < 0.01; ***p-value < 0.001) if not stated otherwise in the text.

### Simulations

Simulations of the time course evolution of oligomeric species concentration were performed using KinTek Global Kinetic Explorer software23^59^ based on a kinetic model where the oligomeric species (S2 to S6) resulted from monomer and dimer association and dissociation steps as described in Supplementary Fig. 31A. As input for the simulations we used experimentally obtained association rate constants (k_12_, k_23_, k_34_, k_45_, K_56_, k_24_, k_26_ in μM^-1^ s^-1^) and dissociation rate constants (k_21_, k_32_, k_42_, k_43_, k_54_, k_62_, k_65_ in s^-1^) for each of the four experimental setups (Supplementary Table 16-19). The only constant missing from all experimental data sets is dimer dissociation rate, k_64_. This value was too rare to be quantified, which suggests that the value is low in all cases. We can thus assign an upper bound based on the number of events that would be observable. For k_64_, we used a value of 0.01 s^-1^ which is an order of magnitude slower than the monomer dissociation from the hexamer. The rate k_42_ could only be reliably extracted for the experiments containing phenol and Zn^2+^ + phenol and the experimental value from phenol was extended to the other conditions (HI and HI + Zn^2+^). The missing dissociation rate constants were supplied as follows: k_64_ was arbitrarily set to an upper limit value of 0.01 s^-1^ for the four conditions. k_42_, obtained from the experiment with phenol, was used to perform simulations in presence of Zn^2+^ and in absence of oligomer stabilizers. k_26_ and k_62_ were only observed for the combination of Zn^2+^ and phenol, and were set as zero in other conditions. The reactions were simulated using a default sigma value of 0.001 in 200 steps for a reaction time of 300 s with HI^655^ (S1) initial concentration varying from 10 nM to 100 mM. The oligomer fractions were then calculated considering 1 the sum of the end point concentrations of each oligomeric species (Supplementary Fig. 32-35).

The mean oligomeric state was calculated as

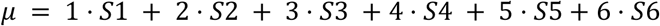

where S1 to S6 represents the fractions of monomers to hexamers displayed in bars in Fig. 5ABCD.

Evolution of hexamers in Fig. 5ABCD have been fitted with the Hill equation to find apparent affinity K:

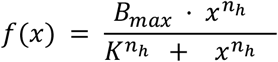

where K is the concentration needed for half-maximum hexamer formation. n_h_ is the hill coefficient, and B_max_ is the maximum hexamer fraction^47^,^60^.

## Acknowledgements

N.S.H. and K.J.J acknowledges funding from Villum foundation center BIONEC (grant grant 18333) N.S.H. acknowledges funding from Villum foundation project (grant 40801) Carlsberg Foundation Distinguished Associate Professor Program (CF16-0797) and the NovoNordisk Center for Biopharmaceuticals and Biobarriers in Drug Delivery (NNF16OC0021948). The Novo Nordisk Foundation Center for Protein Research (CPR) is funded by a generous donation from the Novo Nordisk Foundation (grant no. NNF14CC0001). N.S.H. is a member of the Integrative Structural Biology Cluster (ISBUC) at the University of Copenhagen.

## Author contributions

N.S.H. designed the study with K.J.J. and serious inputs from F.B., S.S.-R.B.. N.K.M. performed insulin synthesis and labelling, purification, characterization, enzymatic assay and dynamic light scattering experiments. F.B. performed all TIRF microscopy experiments and designed the controls, together with E.M.N. and M.Z.. S.S.-R.B. and F.B. developed the data analysis, and H.D.P. developed the method for conversion from intensity to photons. F.B. carried out all data analysis and interpretation. N.S.G. and M.K. performed simulations. N.S.H. with tight interaction with K.J.J. had the overall supervision of the project. F.B. and N.S.H wrote the manuscript with input from all coauthors.

## Data availability

The authors declare that the data supporting the findings of this study are available within the paper and its Supplementary information files. Additional and relevant data are available from the corresponding authors on reasonable request.

## Code availability

Codes used for localization, extraction of single molecule trajectories and HMM fitting will be fully available on Hatzakislab github after peer review acceptance.

## Bibliography

1. Saltiel, A. R. & Kahn, C. R. Insulin signalling and the regulation of glucose and lipid metabolism. Nature 414, 799–806 (2001).

2. Home, P. The challenge of poorly controlled diabetes mellitus. Diabetes Metab. 29, 101–109 (2003).

3. Hermansen, K. et al. Insulin analogues (insulin detemir and insulin aspart) versus traditional human insulins (NPH insulin and regular human insulin) in basal-bolus therapy for patients with type 1 diabetes. Diabetologia 47, 622–629 (2004).

4. Zaykov, A. N., Mayer, J. P. & DiMarchi, R. D. Pursuit of a perfect insulin. Nat. Rev. Drug Discov. 15, 425–439 (2016).

5. Kramer, C. K., Retnakaran, R. & Zinman, B. Insulin and insulin analogs as antidiabetic therapy: A perspective from clinical trials. Cell Metab. 33, 740–747 (2021).

6. Mathieu, C., Gillard, P. & Benhalima, K. Insulin analogues in type 1 diabetes mellitus:getting better all the time. Nat. Rev. Endocrinol. 13, 385–399 (2017).

7. Bakh, N. A. et al. Glucose-responsive insulin by molecular and physical design. Nat.Chem. 9, 937–943 (2017).

8. Søeborg, T., Rasmussen, C. H., Mosekilde, E. & Colding-Jørgensen, M. Absorption kinetics of insulin after subcutaneous administration. Eur. J. Pharm. Sci. 36, 78–90 (2009).

9. Chitta, R. K., Rempel, D. L., Grayson, M. A., Remsen, E. E. & Gross, M. L. Application *of SIMSTEX to oligomerization of insulin analogs and mutants*. J. Am. Soc. Mass Spectrom. 17, 1526–1534 (2006).

10. Pocker, Y. & Biswas, S. B. Self-association of insulin and the role of hydrophobic bonding: a thermodynamic model of insulin dimerization. Biochemistry 20, 4354–4361 (1981).

11. Baker, E. N. et al. The structure of 2Zn pig insulin crystals at 1.5 A resolution. Philos. Trans. R. Soc. Lond. B Biol. Sci. 319, 369–456 (1988).

12. Blundell, T., Dodson, G., Hodgkin, D. & Mercola, D. Insulin: the structure in the crystal and its reflection in chemistry and biology by. in vol. 26 279–402 (Elsevier, 1972).

13. Lisi, G. P., Png, C. Y. M. & Wilcox, D. E. Thermodynamic contributions to the stability of the insulin hexamer. Biochemistry 53, 3576–3584 (2014).

14. Carpenter, M. C. & Wilcox, D. E. Thermodynamics of formation of the insulin hexamer:metal-stabilized proton-coupled assembly of quaternary structure. Biochemistry 53, 1296–1301 (2014).

15. Chothia, C., Lesk, A. M., Dodson, G. G. & Hodgkin, D. C. Transmission of conformational change in insulin. Nature 302, 500–505 (1983).

16. Brader, M. L. Zinc coordination, asymmetry, and allostery of the human insulin hexamer. J. Am. Chem. Soc. 119, 7603–7604 (1997).

17. Pekar, A. H. & Frank, B. H. Conformation of proinsulin. A comparison of insulin and proinsulin self-association at neutral pH. Biochemistry 11, 4013–4016 (1972).

18. Hvidt, S. Insulin association in neutral solutions studied by light scattering. Biophys. Chem. 39, 205–213 (1991).

19. Derewenda, U. et al. Phenol stabilizes more helix in a new symmetrical zinc insulin hexamer. Nature 338, 594–596 (1989).

20. Choi, W. E., Brader, M. L., Aguilar, V., Kaarsholm, N. C. & Dunn, M. F. The allosteric transition of the insulin hexamer is modulated by homotropic and heterotropic interactions. Biochemistry 32, 11638–11645 (1993).

21. Berchtold, H. & Hilgenfeld, R. Binding of phenol to R6 insulin hexamers. Biopolymers 51, 165–172 (1999).

22. Huus, K. et al. Ligand binding and thermostability of different allosteric states of the insulin zinc-hexamer. Biochemistry 45, 4014–4024 (2006).

23. Brzović, P. S., Choi, W. E., Borchardt, D., Kaarsholm, N. C. & Dunn, M. F. Structural asymmetry and half-site reactivity in the T to R allosteric transition of the insulin hexamer. Biochemistry 33, 13057–13069 (1994).

24. Boga Raja, U. K., Injeti, S., Culver, T., McCabe, J. W. & Angel, L. A. Probing the stability of insulin oligomers using electrospray ionization ion mobility mass spectrometry. Eur J Mass Spectrom (Chichester, Eng) 21, 759–774 (2015).

25. Daly, M. E. et al. Acute effects on insulin sensitivity and diurnal metabolic profiles of a high-sucrose compared with a high-starch diet. Am. J. Clin. Nutr. 67, 1186–1196 (1998).

26. Østergaard, M., Mishra, N. K. & Jensen, K. J. The ABC of insulin: the organic chemistry of a small protein. Chem. Eur. J 26, 8341–8357 (2020).

27. Hassiepen, U., Federwisch, M., Mülders, T. & Wollmer, A. The lifetime of insulin hexamers. Biophys. J. 77, 1638–1654 (1999).

28. Stella, S. et al. Conformational Activation Promotes CRISPR-Cas12a Catalysis and Resetting of the Endonuclease Activity. Cell 175, 1856–1871.e21 (2018).

29. Thomsen, R. P. et al. A large size-selective DNA nanopore with sensing applications. Nat. Commun. 10, 5655 (2019).

30. Thomsen, J. et al. DeepFRET, a software for rapid and automated single-molecule FRET data classification using deep learning. eLife 9, (2020).

31. Bohr, S. S.-R. et al. Direct observation of Thermomyces lanuginosus lipase diffusional states by Single Particle Tracking and their remodeling by mutations and inhibition. Sci. Rep. 9, 16169 (2019).

32. Moses, M. E. et al. Single-Molecule Study of Thermomyces lanuginosus Lipase in a Detergency Application System Reveals Diffusion Pattern Remodeling by Surfactants and Calcium. ACS Appl. Mater. Interfaces 13, 33704–33712 (2021).

33. Jensen, S. B. et al. Biased cytochrome P450-mediated metabolism via small-molecule ligands binding P450 oxidoreductase. Nat. Commun. 12, 2260 (2021).

34. Moses, M. E., Hedegård, P. & Hatzakis, N. S. Quantification of functional dynamics of membrane proteins reconstituted in nanodiscs membranes by single turnover functional readout. Meth. Enzymol. 581, 227–256 (2016).

35. Montoya, G., Hatzakis, N. & et. al. Conformational Activation of CRISPR-Cpf1 Catalysis and Endonuclease Function Recycling. Submitted (2018).

36. Bavishi, K. et al. Direct observation of multiple conformational states in Cytochrome P450 oxidoreductase and their modulation by membrane environment and ionic strength. Sci. Rep. 8, 6817 (2018).

37. Laursen, T. et al. Single molecule activity measurements of cytochrome P450 oxidoreductase reveal the existence of two discrete functional states. ACS Chem. Biol. 9, 630–634 (2014).

38. Streck, S. et al. Interactions of Cell-Penetrating Peptide-Modified Nanoparticles with Cells Evaluated Using Single Particle Tracking. ACS Appl. Bio Mater. (2021) doi:10.1021/acsabm.0c01563.

39. Mortensen, K. I., Churchman, L. S., Spudich, J. A. & Flyvbjerg, H. Optimized localization analysis for single-molecule tracking and super-resolution microscopy. Nat. Methods 7, 377–381 (2010).

40. Brems, D. N. et al. Altering the association properties of insulin by amino acid replacement. Protein Eng. 5, 527–533 (1992).

41. Hermansen, K., Bohl, M. & Schioldan, A. G. Insulin aspart in the management of diabetes mellitus: 15 years of clinical experience. Drugs 76, 41–74 (2016).

42. Gast, K. et al. Rapid-Acting and Human Insulins: Hexamer Dissociation Kinetics upon Dilution of the Pharmaceutical Formulation. Pharm. Res. 34, 2270–2286 (2017).

43. Rahuel-Clermont, S., French, C. A., Kaarsholm, N. C., Dunn, M. F. & Chou, C. I. Mechanisms of stabilization of the insulin hexamer through allosteric ligand interactions. Biochemistry 36, 5837–5845 (1997).

44. Dunn, M. F. Zinc-ligand interactions modulate assembly and stability of the insulin hexamer -- a review. Biometals 18, 295–303 (2005).

45. Cunningham, L. W., Fischer, R. L. & Vestling, C. S. A study of the binding of zinc and cobalt by insulin. J. Am. Chem. Soc. 77, 5703–5707 (1955).

46. Fisher, A. M. THE EFFECT OF ZINC SALTS ON THE ACTION OF INSULIN. Journal of Pharmacology and Experimental Therapeutics 55, 206–221 (1935).

47. Sevlever, F., Di Bella, J. P. & Ventura, A. C. Discriminating between negative cooperativity and ligand binding to independent sites using pre-equilibrium properties of binding curves. PLoS Comput. Biol. 16, e1007929 (2020).

48. Coffman, F. D. & Dunn, M. F. Insulin-metal ion interactions: the binding of divalent cations to insulin hexamers and tetramers and the assembly of insulin hexamers. Biochemistry 27, 6179–6187 (1988).

49. Kadima, W. et al. The influence of ionic strength and pH on the aggregation properties of zinc-free insulin studied by static and dynamic laser light scattering. Biopolymers 33, 1643–1657 (1993).

50. Mannerstedt, K. et al. An aldehyde responsive, cleavable linker for glucose responsive insulins. Chem. Eur. J 27, 3166–3176 (2021).

51. Munch, H. K. et al. Controlled self-assembly of re-engineered insulin by Fe(II). Chem. Eur. J 17, 7198–7204 (2011).

52. Munch, H. K. et al. Construction of Insulin 18-mer Nanoassemblies Driven by Coordination to Iron(II) and Zinc(II) Ions at Distinct Sites. Angew Chem Int Ed Engl 55, 2378–2381 (2016).

53. Bohr, S. S.-R., Thorlaksen, C., Kühnel, R. M., Günther-Pomorski, T. & Hatzakis, N. S. Label-Free Fluorescence Quantification of Hydrolytic Enzyme Activity on Native Substrates Reveals How Lipase Function Depends on Membrane Curvature. Langmuir 36, 6473–6481 (2020).

54. Singh, P. K., Bohr, S. S.-R. & Hatzakis, N. S. Direct observation of sophorolipid micelle docking in model membranes and cells by single particle studies reveals optimal fusion conditions. Biomolecules 10, (2020).

55. Chao, J., Ward, E. S. & Ober, R. J. Fisher information matrix for branching processes with application to electron-multiplying charge-coupled devices. Multidimens. Syst. Signal Process. 23, 349–379 (2012).

56. Ulbrich, M. H. & Isacoff, E. Y. Subunit counting in membrane-bound proteins. Nat. Methods 4, 319–321 (2007).

57. McKinney, S. A., Joo, C. & Ha, T. Analysis of single-molecule FRET trajectories using hidden Markov modeling. Biophys. J. 91, 1941–1951 (2006).

58. Atkins, P. & De Paula, J. Atkins’ Physical Chemistry. (W.H. Freeman and Company, 2010).

59. Johnson, K. A., Simpson, Z. B. & Blom, T. Global kinetic explorer: a new computer program for dynamic simulation and fitting of kinetic data. Anal. Biochem. 387, 20–29 (2009).

60. © 1995-2022 GraphPad Software, LLC. All rights reserved. Specific binding with Hill slope. https://www.graphpad.com/guides/prism/latest/curve-fitting/reg_specific_hill.htm.

