## Supplementary Information for "Enhanced hexamerization of insulin via assembly pathway rerouting revealed by single particle studies"

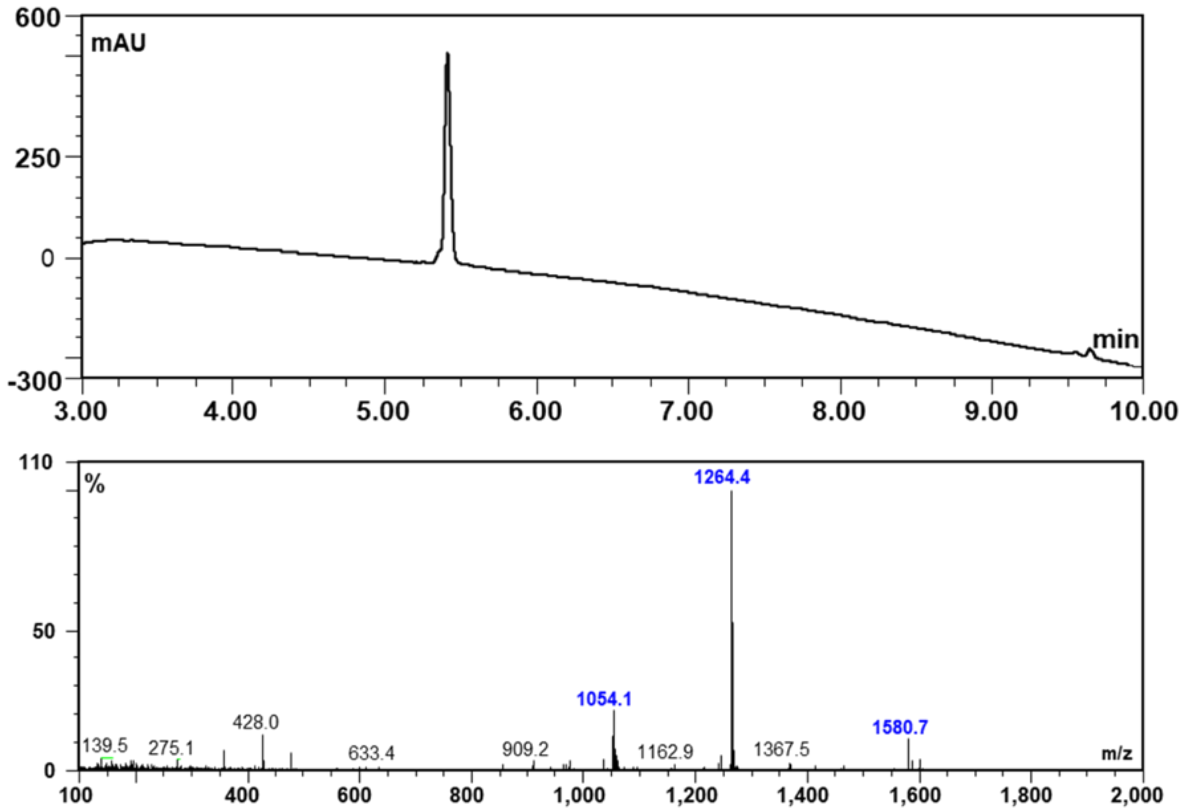


**Supplementary Figure 1:** LCMS chromatogram and mass spectrum for HI$^{655}$ after purification.

| Serial No. | Peak | Calculated | Observed |
|---|---|---|---|
| 1. | $[M]^{+}$ | 6318.26 | Not Observed |
| 2. | $[M+4H]^{4+}$ | 1579.72 | 1580.7 |
| 3. | $[M+5H]^{5+}$ | 1263.98 | 1264.4 |
| 4. | $[M+6H]^{6+}$ | 1053.65 | 1054.1 |

**Supplementary Table 1:** LCMS analysis of HI$^{655.}$

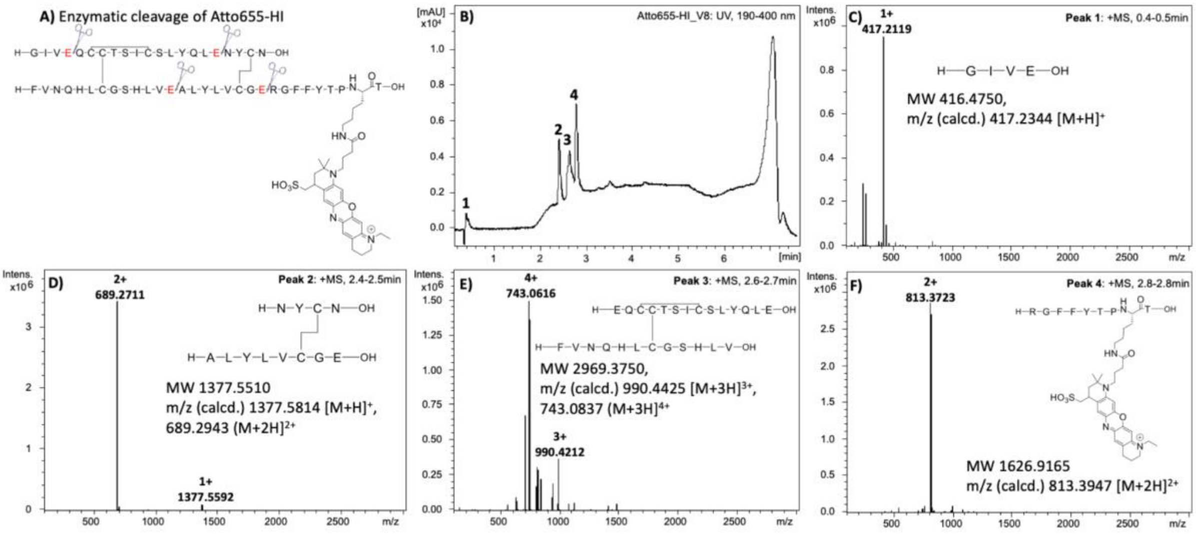


**Supplementary Figure 2:** Enzymatic cleavage study of pure Atto655-HI using Endoproteinase Glu-C (V8 protease) from *Staphyloccus auereus* V8. **A)** The C-terminal V8 cleavage site at glutamic acid (E) indicated in red. **B)** HPLC (UV) chromatogram of pure Atto655-HI treated with V8 protease. Four peaks were observed and showed in their respective mass spectrum **C)** to **F)**. **F)** The observed mass in peak 4 showed the acylation of Atto655 at the B29 Lysine of HI.

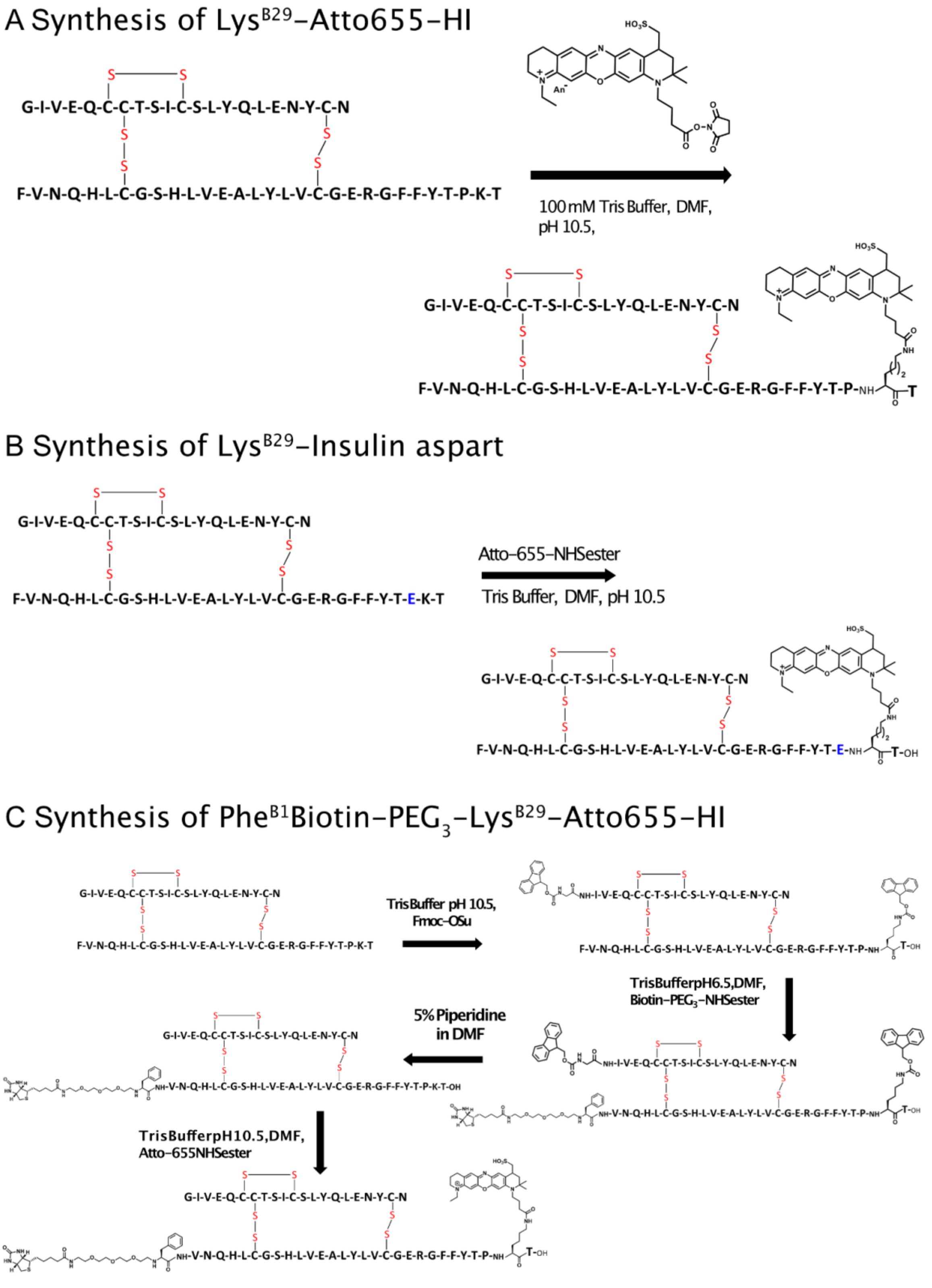


**Supplementary Scheme 1: (A)** For the synthesis of Lys$^{B29}$-Atto655-HI **(A)** For the synthesis of Lys$^{B29}$-Insulin aspart **(C)** For the synthesis of Phe$^{B1}$Biotin-Lys$^{B29}$-Atto655-HI

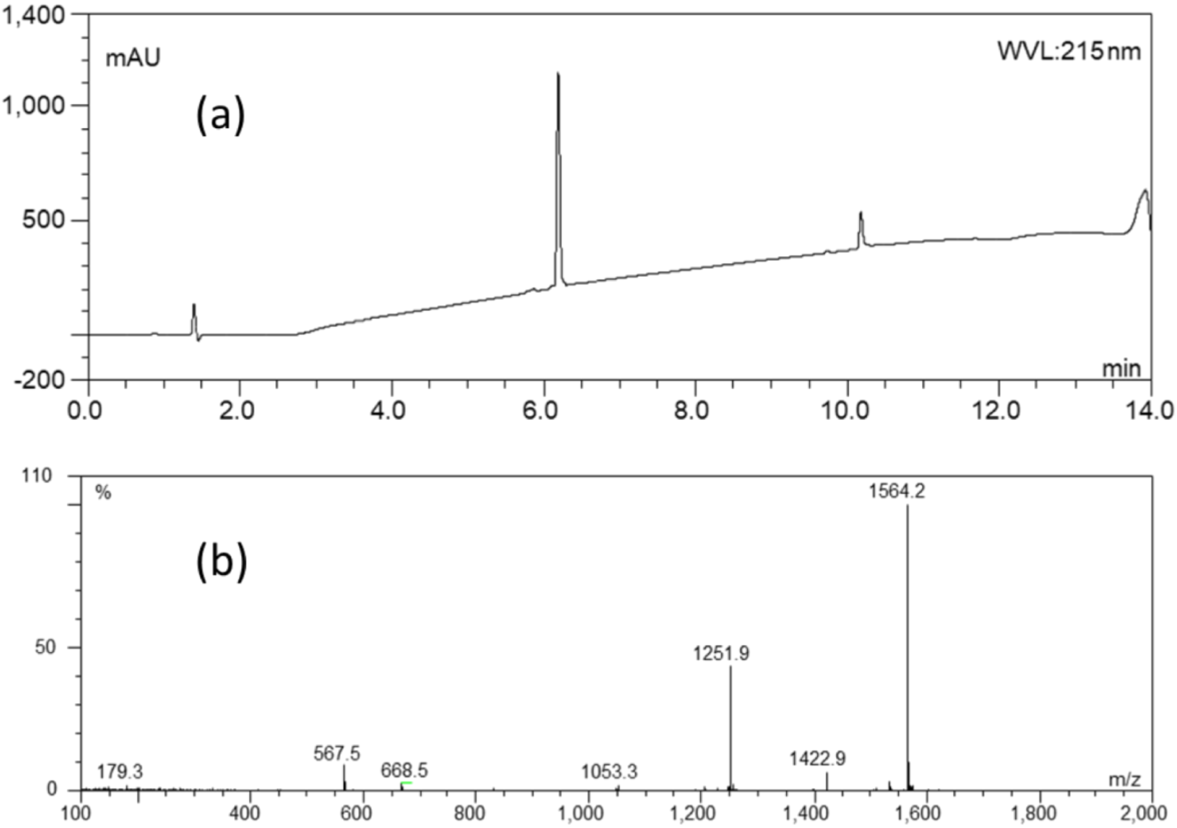


**Supplementary Figure 3:** After purification **(a)** LCMS chromatogram of Di-Fmoc-HI **(b)** Mass spectra of Di-Fmoc-HI

| Serial No. | Peak | Calculated | Observed |
|---|---|---|---|
| 1. | $[M]^{+}$ | 6250.78 | Not Observed |
| 2. | $[M+4H]^{4+}$ | 1563.45 | 1564.2 |
| 3. | $[M+5H]^{5+}$ | 1250.96 | 1251.9 |

**Supplementary Table 2:** LCMS Analysis of Di-Fmoc-HI

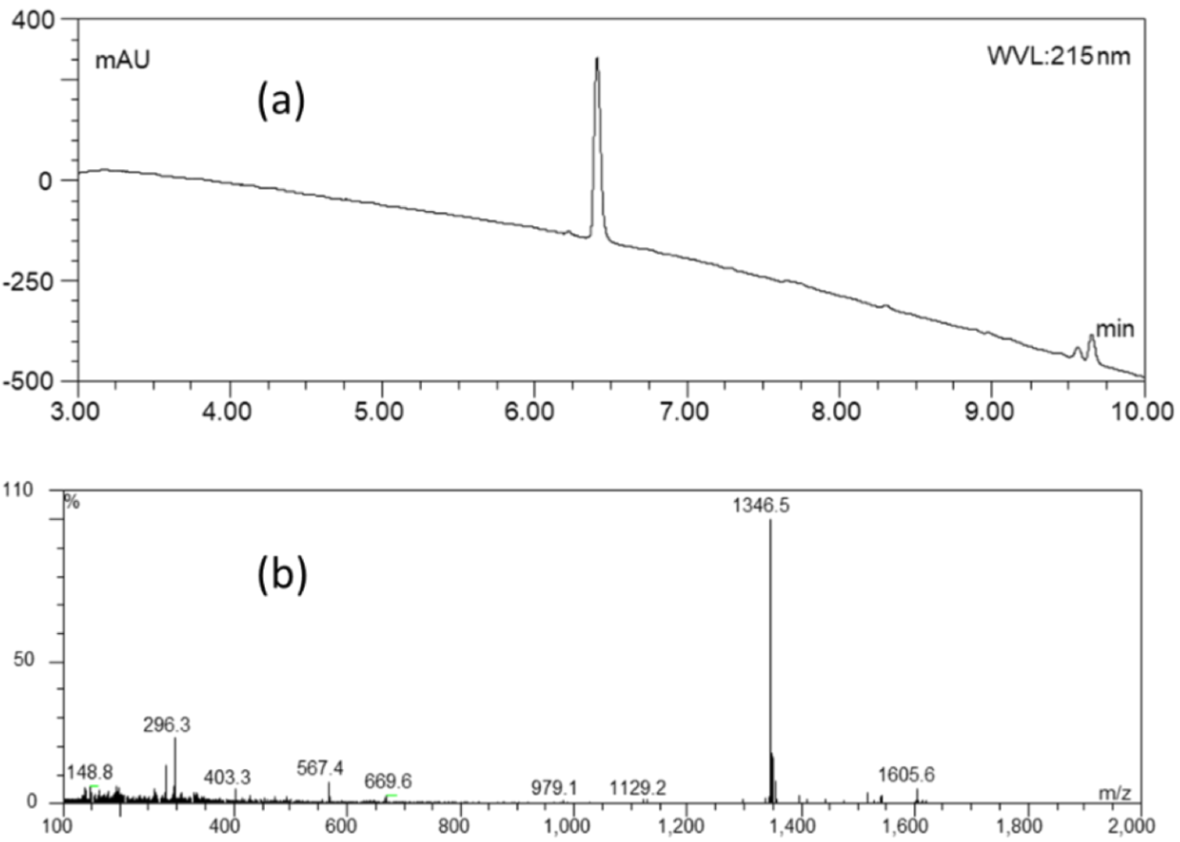


**Supplementary Figure 4:** (a) LCMS chromatogram of DiFmoc-*Phe*$^{B1}$*Biotin-PEG*$_3$-Human Insulin, (b) Mass spectra of DiFmoc-Phe$^{B1}$*Biotin-PEG*$_3$-Human Insulin.

| Serial No. | Peak | Calculated | Observed |
|---|---|---|---|
| 1. | $[M]^+$ | 6720.993 | Not Observed |
| 2. | $[M+4H]^{4+}$ | 1345.808 | 1346.5 |

**Supplementary Table 3:** LCMS Analysis of DiFmoc-Phe$^{B1}$*Biotin-PEG*$_3$-HI

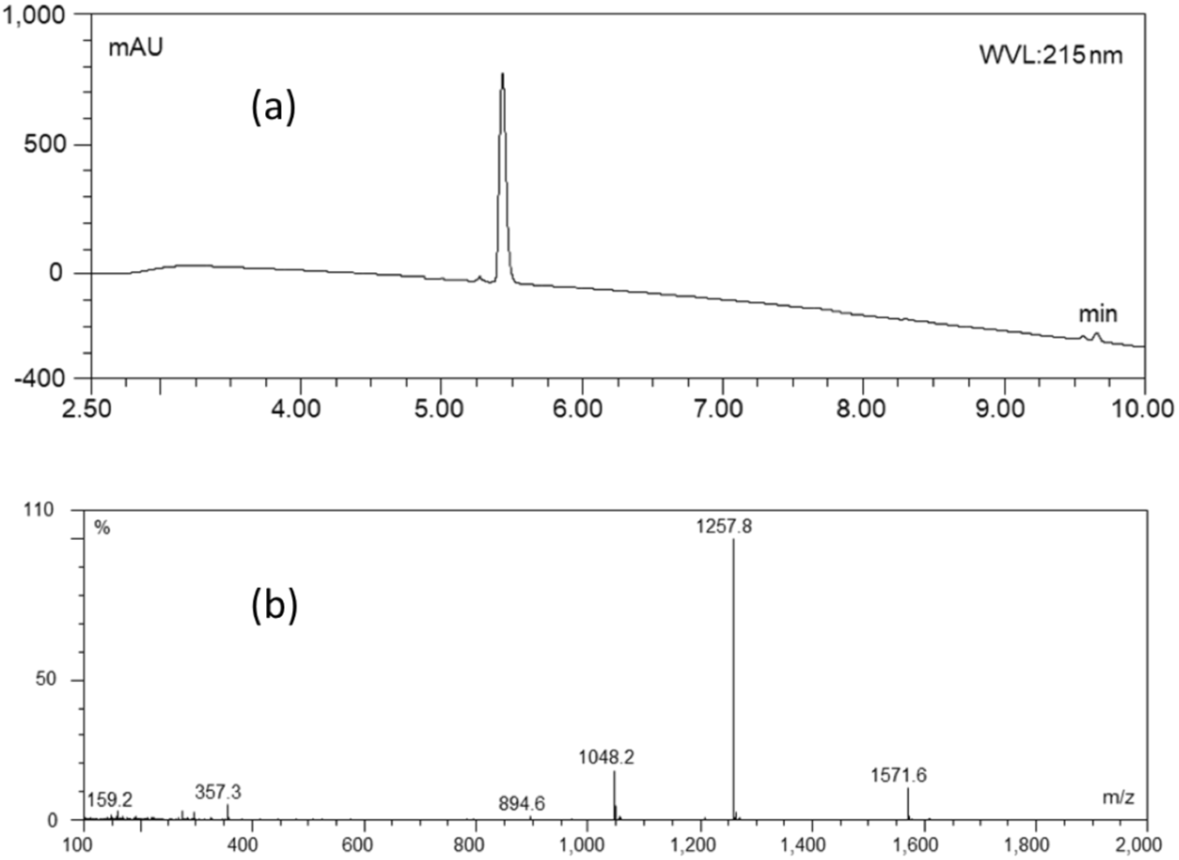


**Supplementary Figure 5:** After purification (a) LCMS chromatogram of *Phe*$^{B1}$*Biotin-PEG$_3$*-HI (b) Mass spectra of *Phe*$^{B1}$Biotin-PEG$_3$-HI

| Serial No. | Peak | Calculated | Observed |
|---|---|---|---|
| 1. | $[M]^+$ | 6250.78 | Not Observed |
| 2. | $[M+4H]^{4+}$ | 1563.45 | 1571.6 |
| 3. | $[M+5H]^{5+}$ | 1256.98 | 1257.8 |

**Supplementary Table 4:** LCMS Analysis of Phe$^{B1}$*Biotin-PEG$_3$*-HI

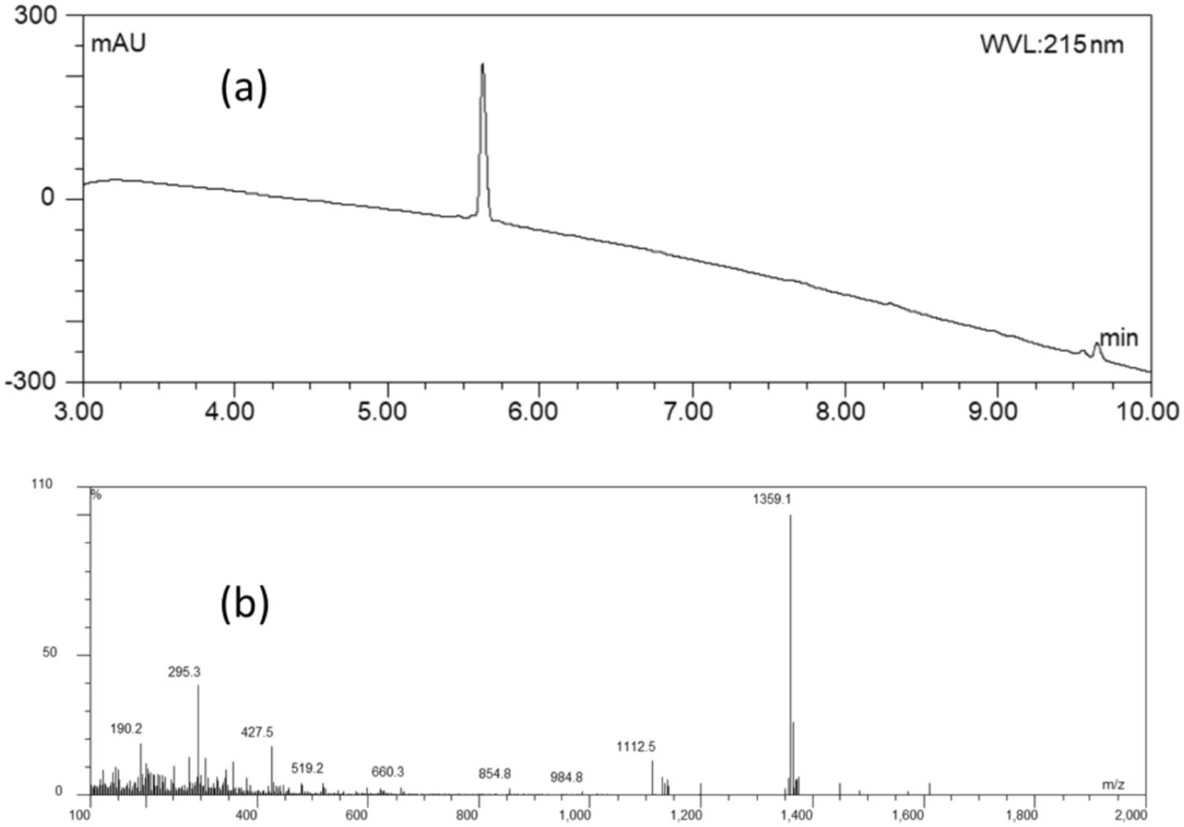


**Supplementary Figure 6:** After purification (a) LCMS chromatogram of Phe$^{B1}$*Biotin-PEG$_3$*-Lys$^{B29}$*Atto-655*-HI (b) Mass spectra of Phe$^{B1}$*Biotin-PEG$_3$*-Lys$^{B29}$*Atto-655*-HI

| Serial No. | Peak | Calculated | Observed |
|---|---|---|---|
| 1. | $[M]^{+}$ | 6787.063 | Not Observed |
| 2. | $[M+4H]^{4+}$ | 1358.820 | 1359.10 |

**Supplementary Table 5:** LCMS Analysis of Phe$^{B1}$-*Biotin-PEG$_3$*-Lys$^{B29}$*Atto-655*-HI

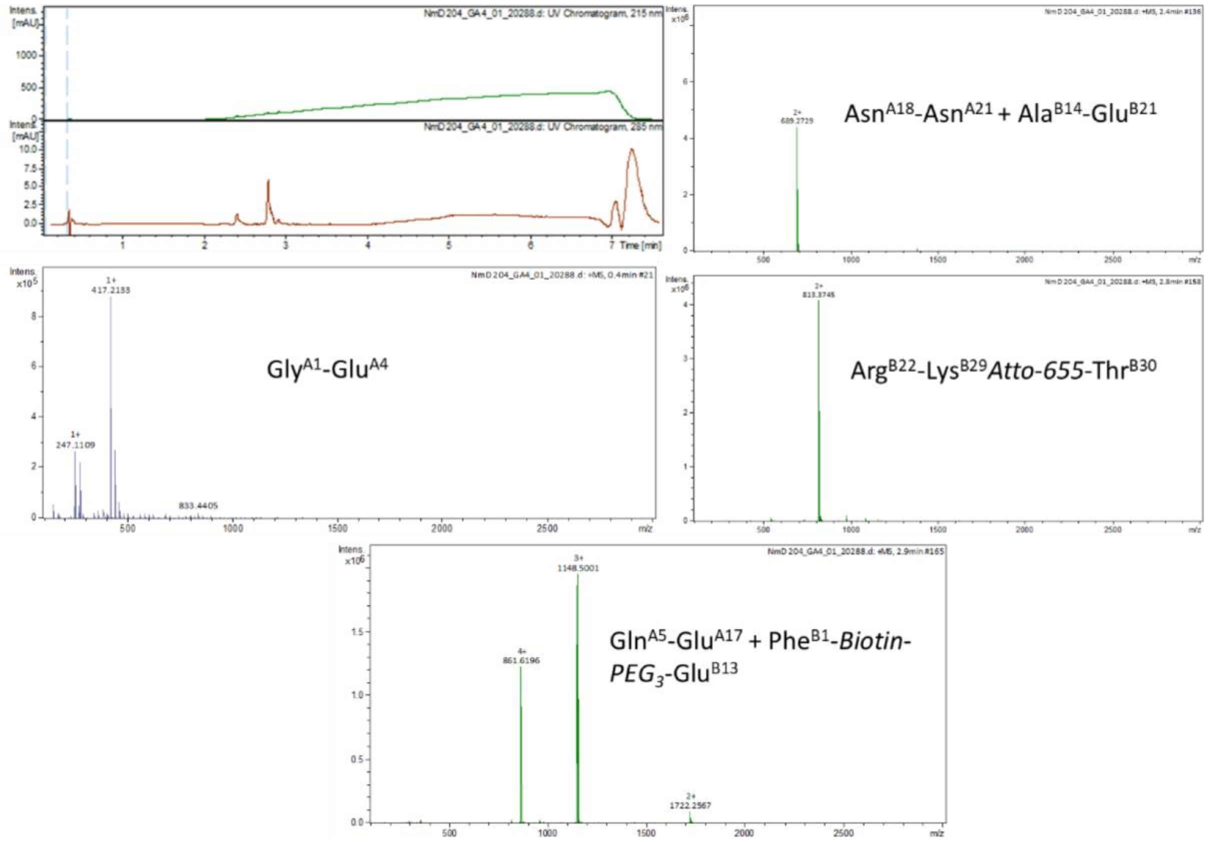


**Supplementary Figure 7:** LCMS chromatogram and mass spectra of Phe$^{B1}$*Biotin-PEG$_3$*-Lys$^{B29}$*Atto-655*-HI after V8 enzymatic cleavage.

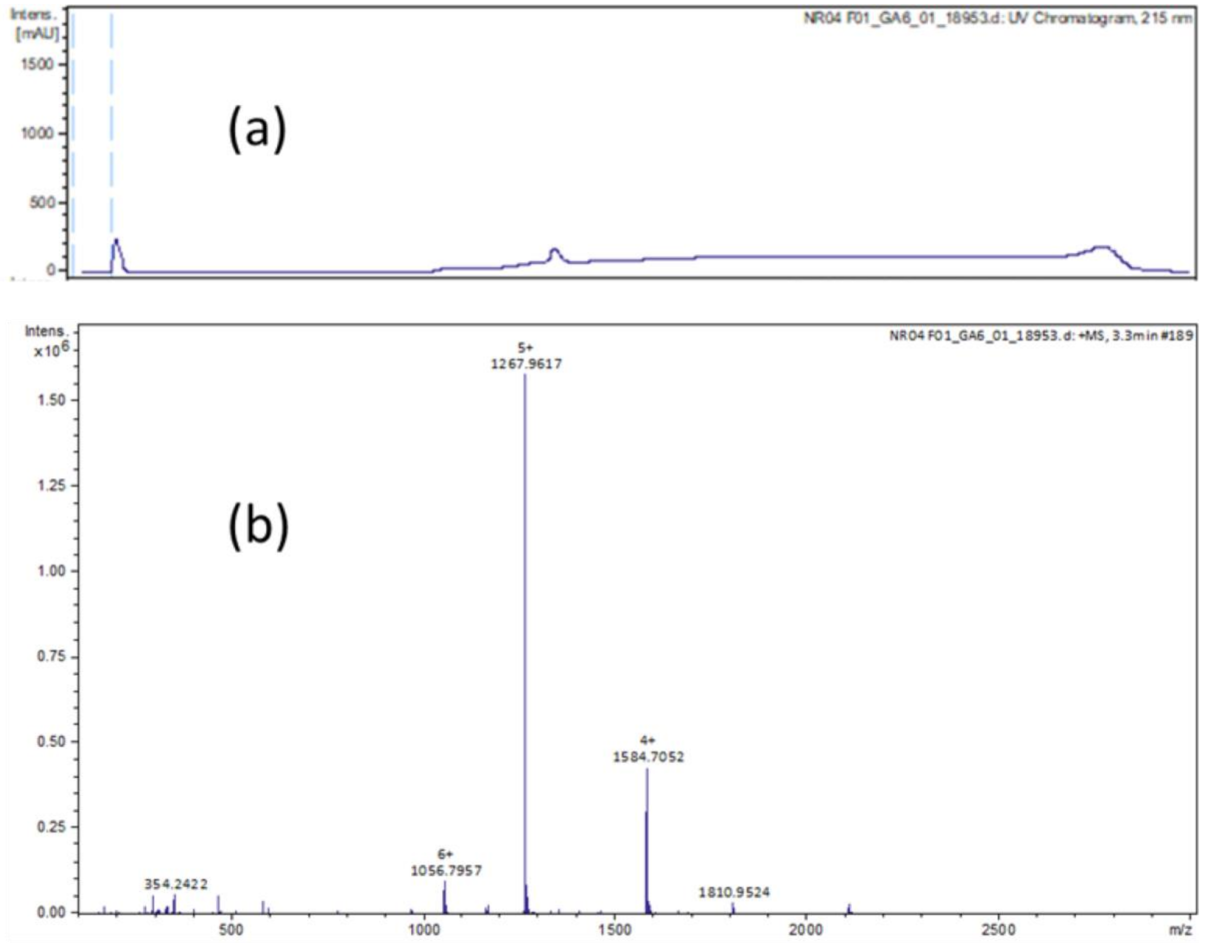


**Supplementary Figure 8:** After purification (a) LCMS chromatogram of NovoRapid-Atto655 (2) Mass spectra of LysB29-Atto655 NovoRapid HI.

| Serial No. | Peak | Calculated | Observed |
|---|---|---|---|
| 1. | $[M]^{+}$ | 6331.81 | Not Observed |
| 2. | $[M+4H]^{4+}$ | 1584.46 | 1584.70 |
| 3. | $[M+5H]^{5+}$ | 1267.77 | 1267.96 |
| 4. | $[M+6H]^{6+}$ | 1056.64 | 1056.79 |

**Supplementary Table 6:** LCMS Analysis of Phe$^{B1}$-*Biotin-PEG*$_{3}$-Lys$^{B29}$*Atto-655*-HI

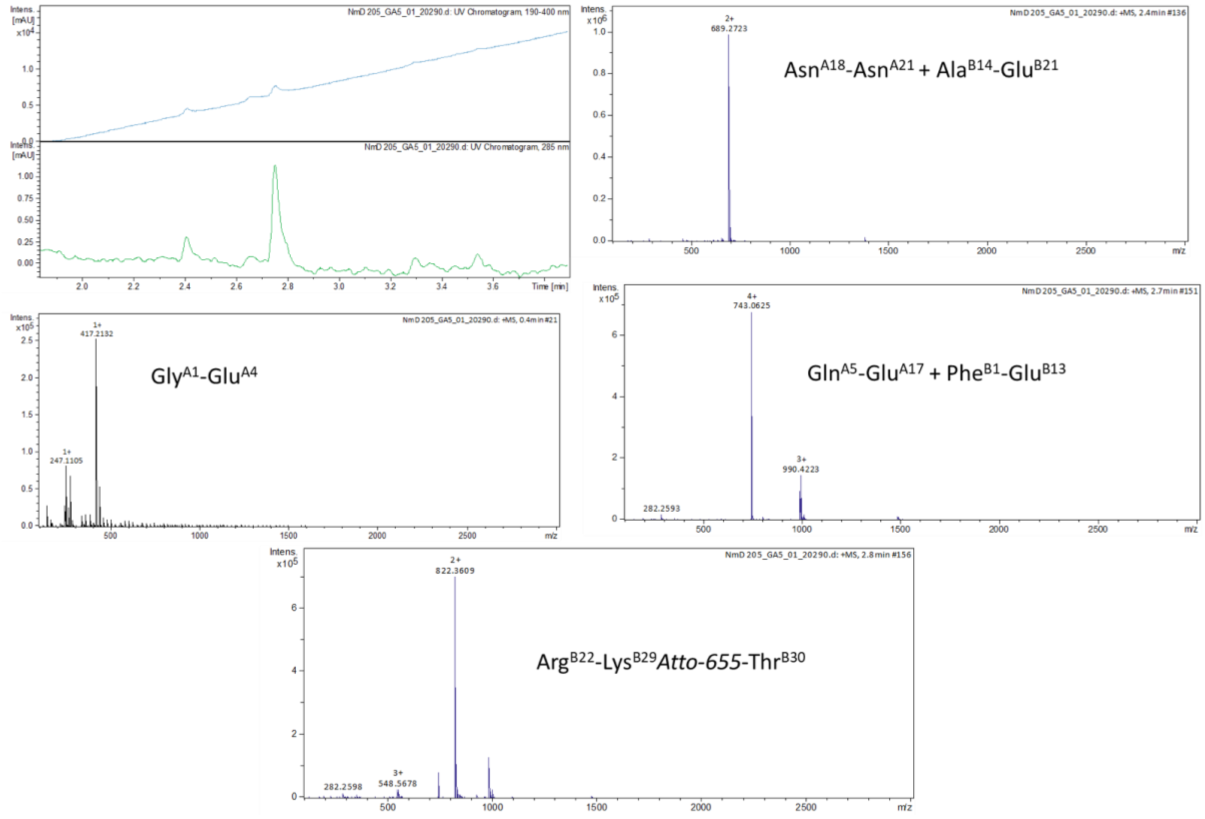


**Supplementary Figure 9:** LCMS chromatogram and mass spectra of Lys$^{B29}$-*Atto 655*-aspart insulin after V8 enzymatic cleavage

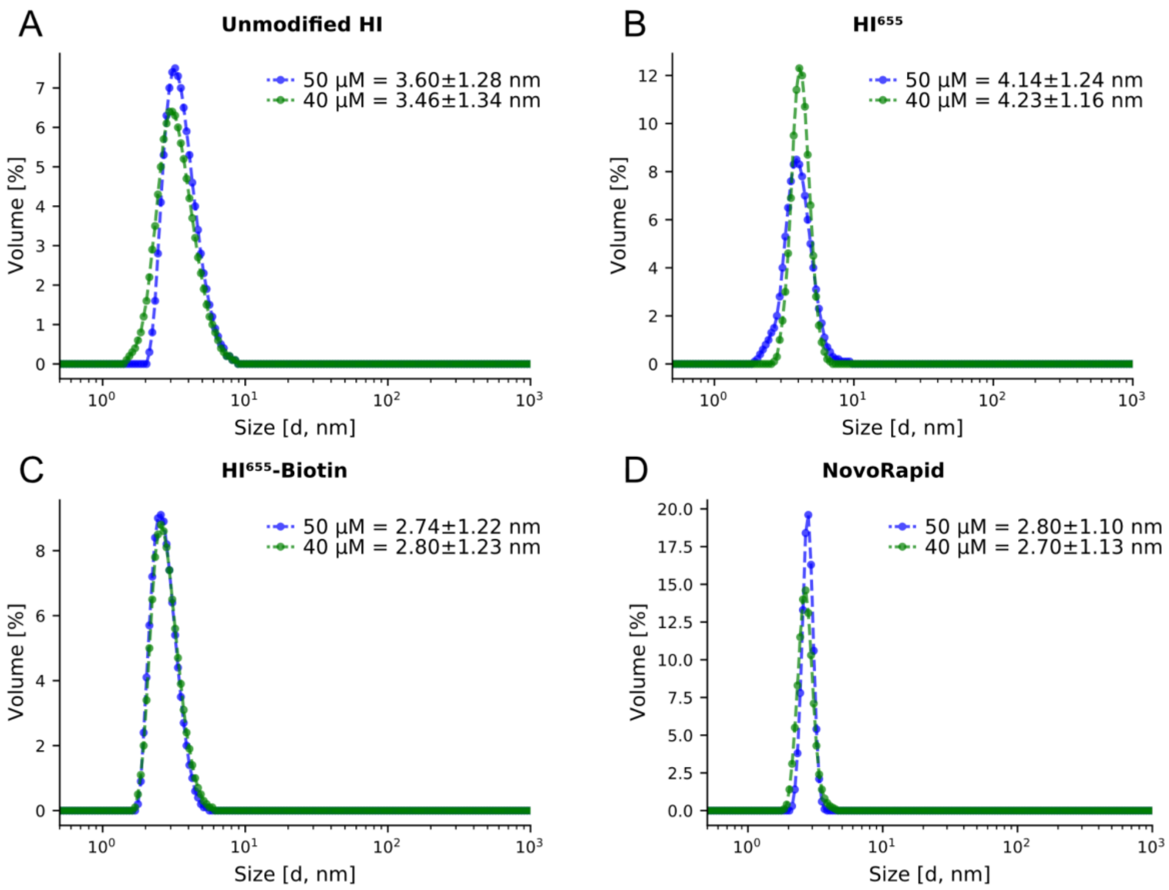


**Supplementary Figure 10:** Dynamic Light Scattering experiments at varying concentration. **(A)** DLS measurements of 50 and 40 µM unmodified HI in 10 mM $Na_2HPO_4$, 10 mM $Na_2HPO_4$, 10 mM $NaH_2PO4$, and 5 mM NaCl at pH 7.5. **(B)** DLS measurements of 50 and 40 µM $HI^{655}$ in 10 mM $Na_2HPO_4$, 10 mM $Na_2HPO_4$, 10 mM $NaH_2PO4$, and 5 mM NaCl at pH 7.5. **(C)** DLS measurements of 50 and 40 µM $HI^{655}$-Biotin in 10 mM $Na_2HPO_4$, 10 mM $Na_2HPO_4$, 10 mM $NaH_2PO4$, and 5 mM NaCl at pH 7.5. **(D)** DLS measurements of 50 and 40 µM NovoRapid in 10 mM $Na_2HPO_4$, 10 mM $Na_2HPO_4$, 10 mM $NaH_2PO4$, and 5 mM NaCl at pH 7.5.

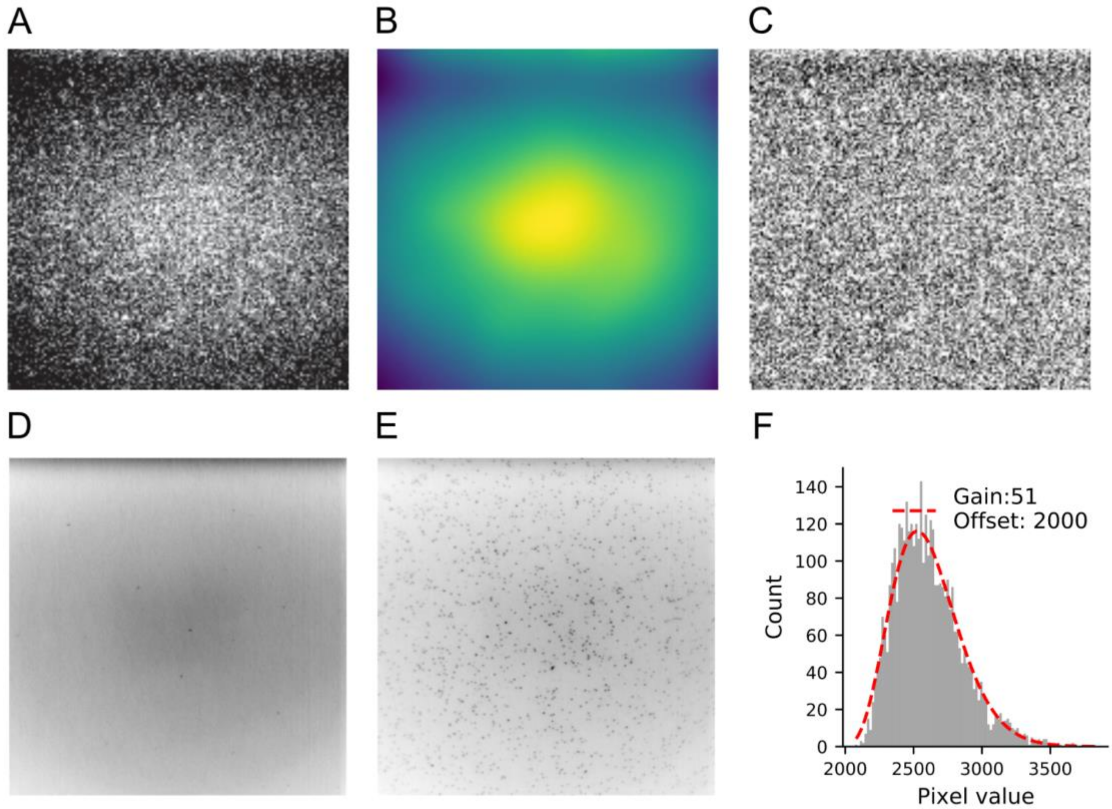


**Supplementary Figure 11:** A-C: Illumination profile correction. **(A)** 512 x 512 pixel microscope image with nonspecifically bound fluorophores before correcting for uneven illumination profile. **(B)** Illumination profile estimated. **(C)** The same microscope image after correcting for uneven illumination profile. D-F: Electron multiplying charge coupled device (EMCCD) calibration directly from a background movie. **(D)** Time averaged movie of a background movie before adding fluorescent particles. **(E)** Time averaged movie of a movie after adding fluorescent particles. **(F)** EMCCD fit of a random pixel in (D) resulting in Gain and Offset.

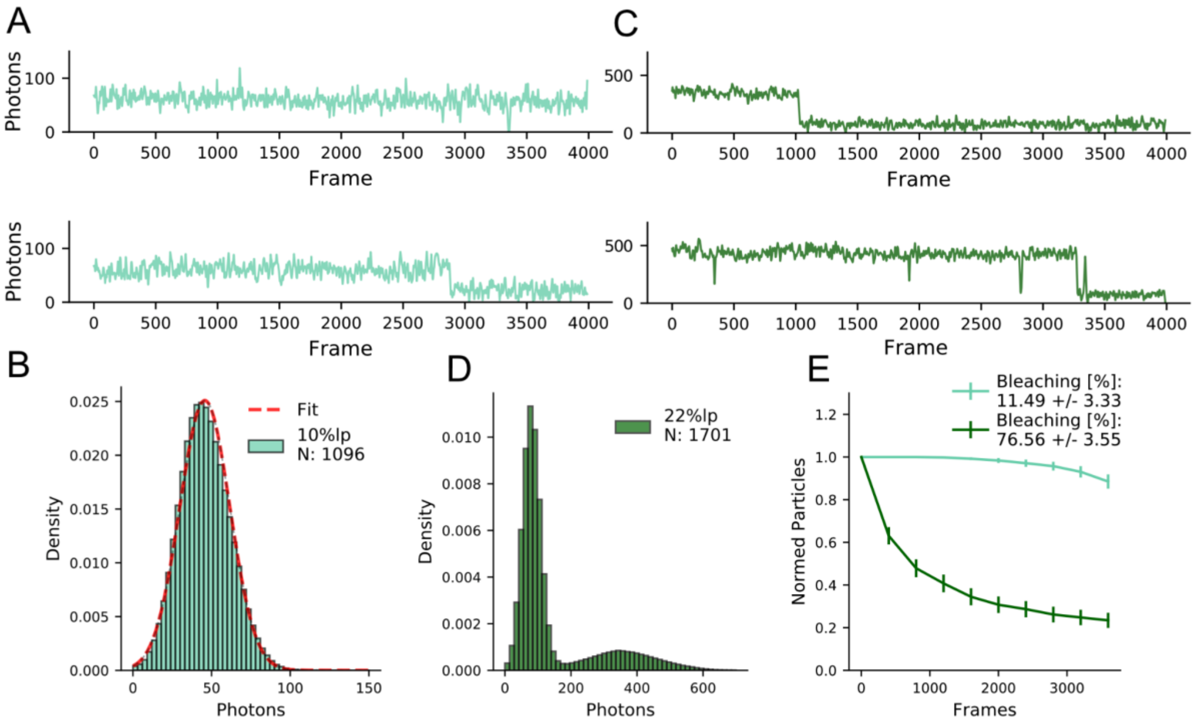


**Supplementary Figure 12:** $HI^{655}$-Biotin control experiment showed fluorophore blinking and bleaching as negligible since we only observe 0.6 % blinking and 12 % bleaching, ensuring minimum bias on observed fluorescent readout. **(A)** Representative traces of single ATTO655 fluorophores coupled to HI. Top shows no bleaching, while bottom shows a single bleaching step for representation. Laser power = 10 % **(B)** Intensity profile for ATTO655 at 10 % laser power. Fit with a Gaussian distribution (red dotted line) reveals a mean of 46 ± 16 photons ($n_{videos}$ = 12). **(C)** Representative traces of single ATTO655 fluorophores at 22 % laser power. Both display single bleaching events, indicative of fluorescent monolabelling. **(D)** Intensity profile for ATTO655 at 22 % laser power showing two distinct distributions. One represents the background, while the other represents ATTO655. **(E)** Quantification of bleaching for 10 % and 22 % laser power reveals 12 % and close to 80 % bleaching respectively, in the full experimental timeframe of 4000 frames. Error bars are the standard deviation from 12 movies respectively.

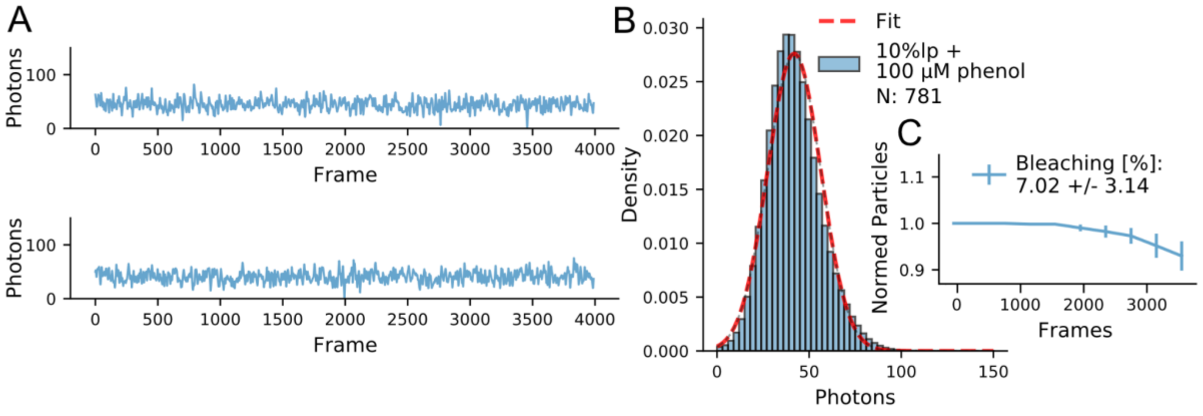


**Supplementary Figure 13:** HI$^{655}$-Biotin control experiment with 100 mikroM phenol showed fluorophore blinking and bleaching as negligible since we only observe 0.5 % blinking and 7 % bleaching, ensuring minimum bias on observed fluorescent readout. **(A)** Representative traces of single ATTO655 fluorophores coupled to HI. Laser power = 10 % **(B)** Intensity profile for ATTO655 at 10 % laser power. Fit with a Gaussian distribution (red dotted line) reveals a mean of 41 ± 14 photons ($N_{videos}$ = 12) **(C)** Quantification of bleaching for 10 % laser power reveals 7 % bleaching, in the full experimental timeframe of 4000 frames. Error bars are the standard deviation from 12 movies respectively.

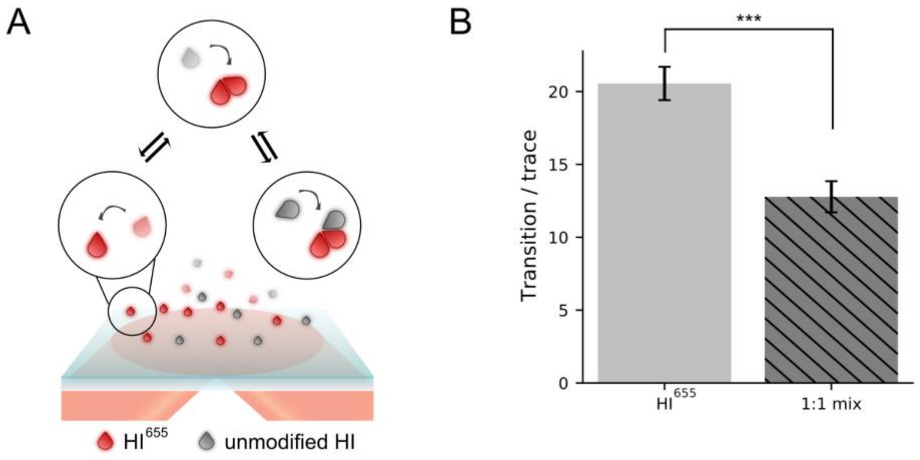


**Supplementary Figure 14:** 1:1 mixture of 10 nM HI$^{655}$ and unmodified HI showed close to 50 % reduction of transitions per trace confirming that the label has little effect on the kinetics. **(A)** Representation (not to scale) of the experimental setup. ATTO655 labeled Human Insulin (HI$^{655}$), are detected upon binding to PLL-PEG/biotin microscopy surface for hexamerization. Particles in solution are not detected due to decay of the evanescence field excitation (pink/shaded red). Unmodified insulin (grey) is not detected either. **(B)** Transition pr trace for the two conditions.

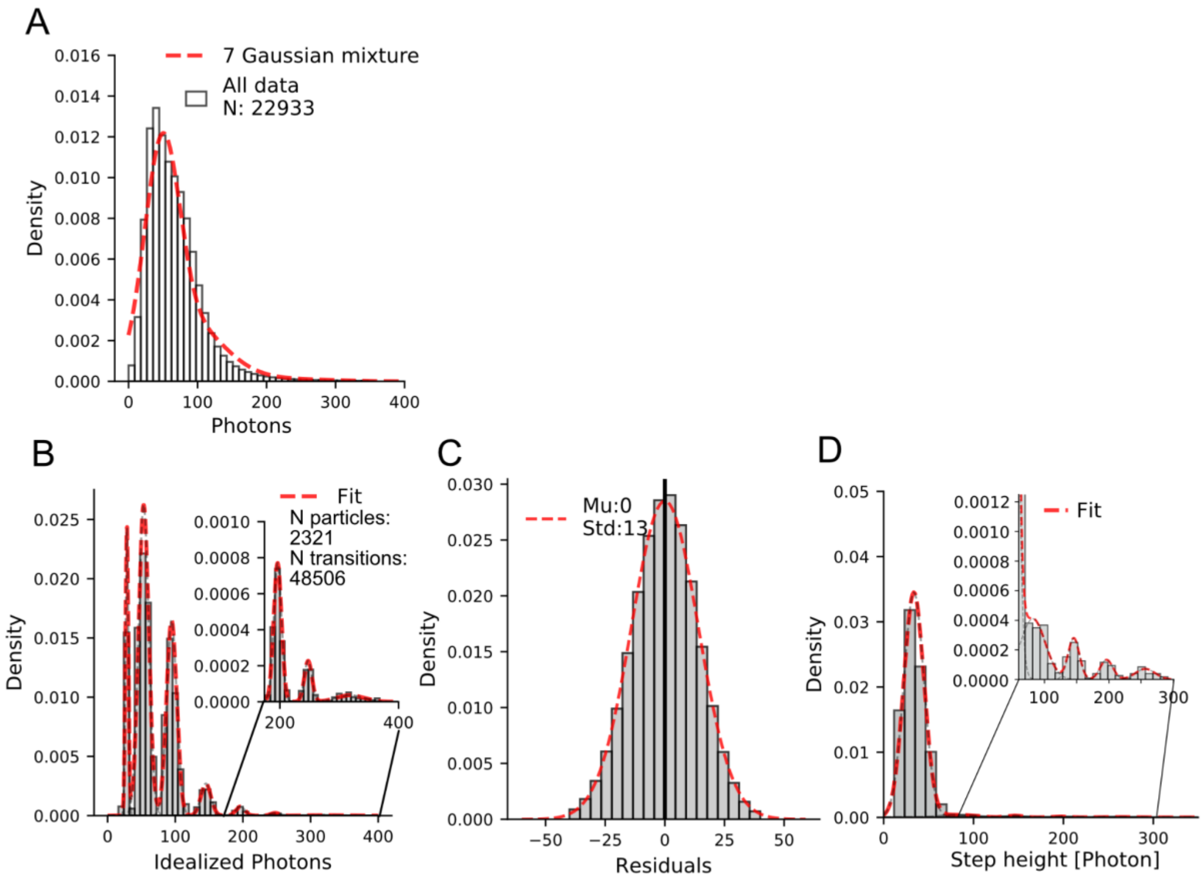


**Supplementary Figure 15:** Hidden Markov Modeling of $HI^{655}$ **(A)** Traces for $HI^{655}$ grouped in a histogram. Red line shows an overlay with a Gaussian mixture model of seven distinct Gaussian distributions indicating that a seven-state model indeed reflects the photon counts observed. N = 22933 is the total number of trajectories. **(B)** Idealized photon histogram for 10 nM $HI^{655}$ with a Gaussian mixture model fit to the seven distinct distributions. **(C)** Residuals for experiments with10 nM $HI^{655}$ grouped in histograms and fitted with Gaussian. μ and σ are displayed for each condition and show no systematic error of HMM fit. **(D)** Estimation of oligomer concentration in solution for 10 nM $HI^{655}$. Histograms show stepheight (essentially the difference between photon count after a transition and photon count before a transition). Distributions correspond to monomer addition, dimer addition, trimer addition, tetramer addition and pentamer addition. Histograms are fitted with five Gaussian distributions to get weight for each distribution as an estimation of oligomer concentration in solution.

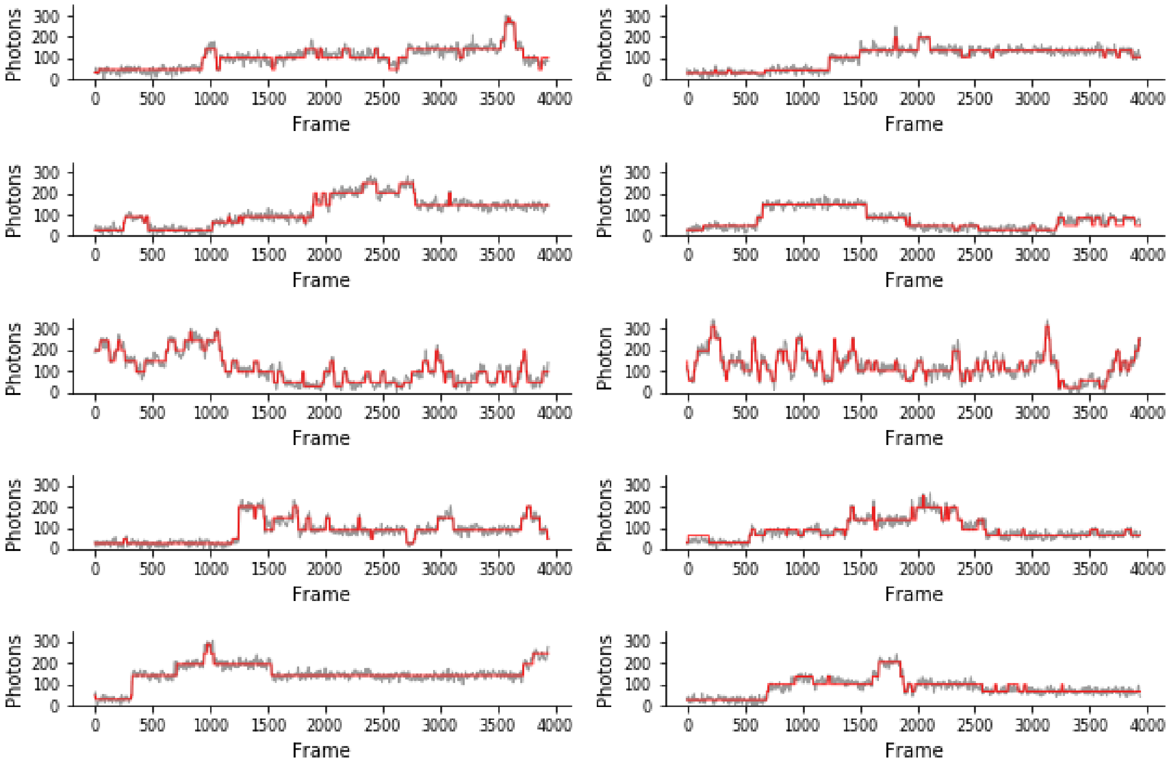


**Supplementary Figure 16:** Representative trajectories overlayed with idealized trace from HMM prediction.

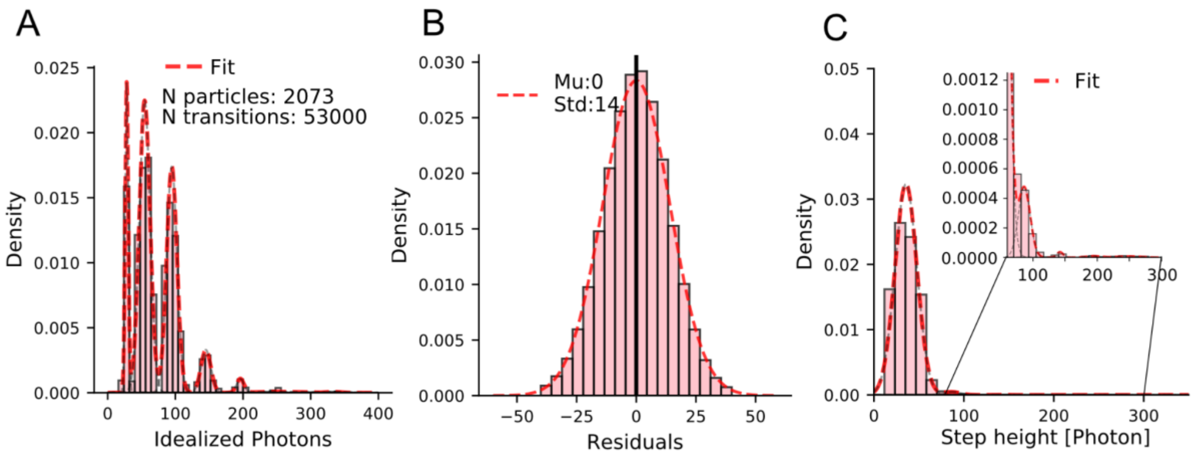


**Supplementary Figure 17:** Hidden Markov Modeling of 10 nM HI$^{655}$ under addition of $Zn^{2+}$ **(A)** Idealized photon histogram for 10 nM + 100 µM $Zn^{2+}$ with a Gaussian mixture model fit to the seven distinct distributions. **(B)** Residuals for experiments with 10 nM HI$^{655}$ + 100 µM $Zn^{2+}$ grouped in histogram and fitted with Gaussian. μ and σ are displayed and show no systematic error in HMM fit. **(C)** Estimation of oligomer concentration on solution. Histogram of stepheight (essentially difference between photon count after a transition and photon count before a transition). Distributions correspond to monomer addition, dimer addition, trimer addition, tetramer addition and pentamer addition. Histogram fitted with five Gaussian distributions to get weight for each distribution.

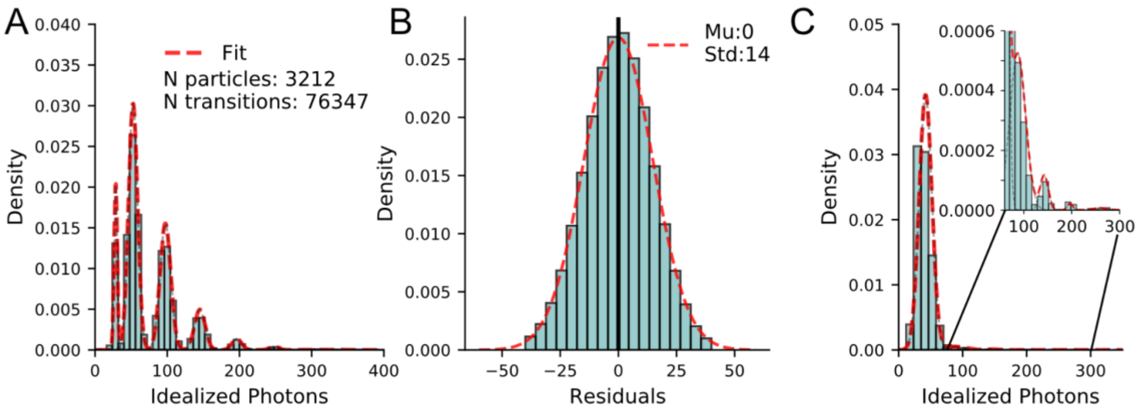


**Supplementary Figure 18:** Hidden Markov Modeling of 10 nM HI[655] under addition of phenol **(A)** Idealized photon histogram for 10 nM + 25 µM phenol with a Gaussian mixture model fit to the seven distinct distributions. **(B)** Residuals for experiments with 10 nM HI[655] + 25 µM phenol grouped in histogram and fitted with Gaussian. μ and σ are displayed and show no systematic error in HMM fit. **(C)** Estimation of oligomer concentration on solution. Histogram of stepheight (essentially difference between photon count after a transition and photon count before a transition). Distributions correspond to monomer addition, dimer addition, trimer addition, tetramer addition and pentamer addition. Histogram fitted with five Gaussian distributions to get weight for each distribution.

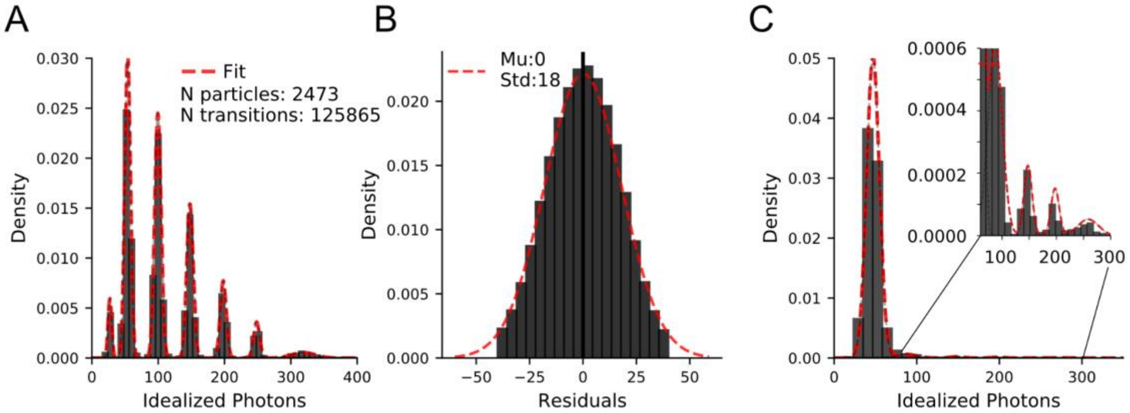


**Supplementary Figure 19:** Hidden Markov Modeling of 10 nM HI$^{655}$ under addition of $Zn^{2+}$ and phenol **(A)** Idealized photon histogram for 10 nM + 100 µM $Zn^{2+}$ + 25 µM phenol with a Gaussian mixture model fit to the seven distinct distributions. **(B)** Residuals for experiments with 10 nM HI$^{655}$ + 100 µM $Zn^{2+}$ + 25 µM phenol grouped in histogram and fitted with Gaussian. µ and σ are displayed and show no systematic error in HMM fit. **(C)** Estimation of oligomer concentration on solution. Histogram of stepheight (essentially difference between photon count after a transition and photon count before a transition). Distributions correspond to monomer addition, dimer addition, trimer addition, tetramer addition and pentamer addition. Histogram fitted with five Gaussian distributions to get weight for each distribution.

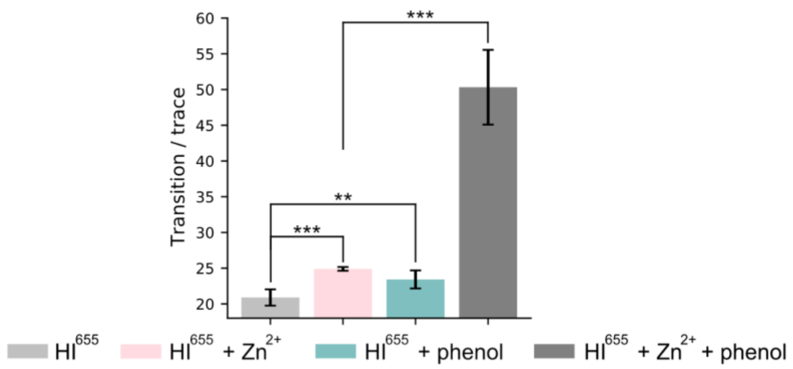


**Supplementary Figure 20**: Transition pr trace show increased dynamics for 10 nM $HI^{655}$ (20.90 ±1.14) upon addition of $Zn^{2+}$ (p-value = 0.0004 (24.90 ± 0.26)) and upon addition with phenol (p-value = 0.009 (24.42 ± 1.26)). Transitions pr trace increase even more when both $Zn^{2+}$ and phenol is added together (p-value = 1.2 * $10^{-5}$ (50.32 ± 5.23)). ( Level of significance is determined by a Welch's t-test (*p-value < 0.05; **p-value < 0.01; ***p-value < 0.001).

| | Monomer | Dimer | Trimer | Tetramer | Pentamer | Hexamer |
|---|---|---|---|---|---|---|
| 10 nM $HI^{655}$ | 56.6 ± 0.0 % | 35.5 ± 0.0 % | 5.8 ± 0.0 % | 1.5 ± 0.0 % | 0.4 ± 0.0 % | 0.2 ± 0.0 % |
| 10 nM $HI^{655}$ + 100 μM $Zn^{2+}$ | 51.7 ± 0.0 % | 38.6 ± 0.0 % | 6.9 ± 0.0 % | 1.7 ± 0.0 % | 0.8 ± 0.1 % | 0.2 ± 0.1 % |
| 10 nM $HI^{655}$I + 25 μM phenol | 53.6 ± 0.0 % | 32.5 ± 0.0 % | 10.2 ± 0.0 % | 2.5 ± 0.0 % | 0.7 ± 0.0 % | 0.4 ± 0.0 % |
| 10 nM $HI^{655}$I + $Zn^{2+}$ + phenol | 33.8 ± 0.0 % | 30.9 ± 0.0 % | 19.4 ± 0.0 % | 9.2 ± 0.0 % | 4.2 ± 0.0 % | 2.5 ± 0.0 % |

**Supplementary Table 7:** State occupancies from Gaussian fit for 10 nM $HI^{655}$ with addition of $Zn^{2+}$ and phenol. Error bars are SEM from 10 x bootstrap fit to the idealized histogram.

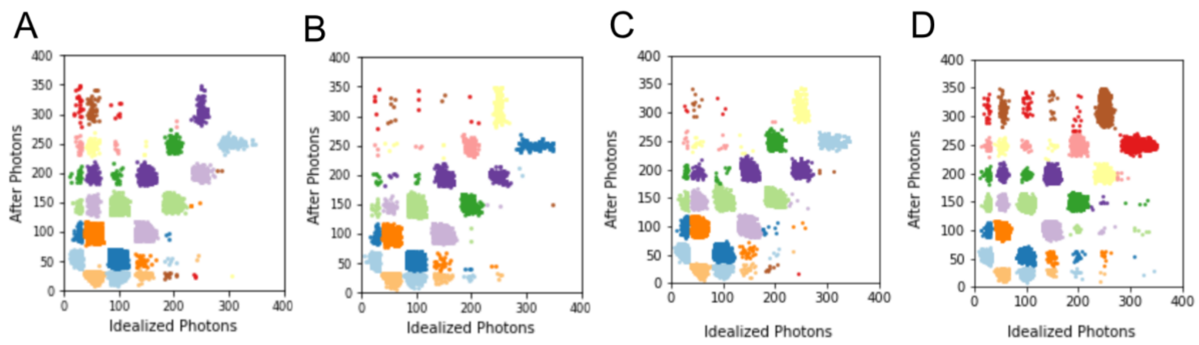


**Supplementary Figure 21:** Transition scatter plot of photon count before a transition and after a transition for experiments with **(A)** 10 nM HI$^{655}$, **(B)** 10 nM HI$^{655}$ + 100 µM $Zn^{2+}$, **(C)** 10 nM HI$^{655}$ + 25 µM phenol and **(D)** 10 nM HI$^{655}$ + 100 µM $Zn^{2+}$ + 25 µM phenol. The color of each cluster has no significance, other than visually easy separation.

| | | | | | | |
|---|---|---|---|---|---|---|
| S6 | 44 | 6 | 0 | 2 | 122 | |
| S5 | 40 | 8 | 5 | 264 | | 149 |
| S4 | 78 | 21 | 692 | | 435 | 2 |
| S3 | 112 | 2155 | | 1006 | 4 | 0 |
| S2 | 14501 | | 2637 | 3 | 0 | 0 |
| S1 | | 14332 | 27 | 6 | 2 | 0 |
| | S1 | S2 | S3 | S4 | S5 | S6 |

**Supplementary Table 8:** Raw count of transitions frequency for all observed transitions for 10 nM $HI^{655}$. Not all transitions have been observed. For some transitions with too little statistics, the rate could not be reliably extracted. Transitions involving S0 have been removed. N = 36653 transitions.

| | | | | | | |
|---|---|---|---|---|---|---|
| S6 | 9 | 2 | 0 | 2 | 542 | |
| S5 | 13 | 6 | 5 | 1208 | | 542 |
| S4 | 47 | 13 | 2689 | | 1255 | 3 |
| S3 | 233 | 6048 | | 2863 | 5 | 0 |
| S2 | 19320 | | 6495 | 10 | 3 | 0 |
| S1 | | 18830 | 27 | 5 | 1 | 0 |
| | S1 | S2 | S3 | S4 | S5 | S6 |

**Supplementary Table 9:** Raw count of transitions frequency for all observed transitions for 10 nM $HI^{655}$ + 25 µM phenol . Not all transitions have been observed. For some transitions with too little statistics, the rate could not be reliably extracted. Transitions involving S0 have been removed. N = 60176 transitions.

| | | | | | | |
|---|---|---|---|---|---|---|
| S6 | 7 | 3 | 2 | 3 | 338 | |
| S5 | 3 | 3 | 3 | 761 | | 340 |
| S4 | 5 | 7 | 1539 | | 784 | 2 |
| S3 | 45 | 3348 | | 1597 | 2 | 1 |
| S2 | 14946 | | 3517 | 1 | 0 | 0 |
| S1 | | 14157 | 24 | 3 | 2 | 0 |
| | S1 | S2 | S3 | S4 | S5 | S6 |

**Supplementary Table 10:** Raw count of transitions frequency for all observed transitions for 10 nM $HI^{655}$ + 100 µM $Zn^{2+}$. Not all transitions have been observed. For some transitions with too little statistics, the rate could not be reliably extracted. Transitions involving S0 have been removed. N = 41443 transitions.

| | | | | | | |
|---|---|---|---|---|---|---|
| S6 | 80 | 33 | 15 | 14 | 3424 | |
| S5 | 98 | 31 | 14 | 6755 | | 3444 |
| S4 | 199 | 89 | 10807 | | 7087 | 7 |
| S3 | 271 | 14464 | | 11473 | 8 | 3 |
| S2 | 21549 | | 15438 | 15 | 8 | 6 |
| S1 | | 21418 | 56 | 22 | 11 | 1 |
| | S1 | S2 | S3 | S4 | S5 | S6 |

**Supplementary Table 11:** Raw count of transitions frequency for all observed transitions for 10 nM $HI^{655}$ + 100 µM $Zn^{2+}$ + 25 µM phenol. Not all transitions have been observed. For some transitions with too little statistics, the rate could not be reliably extracted. Transitions involving S0 have been removed. N = 116840 transitions.

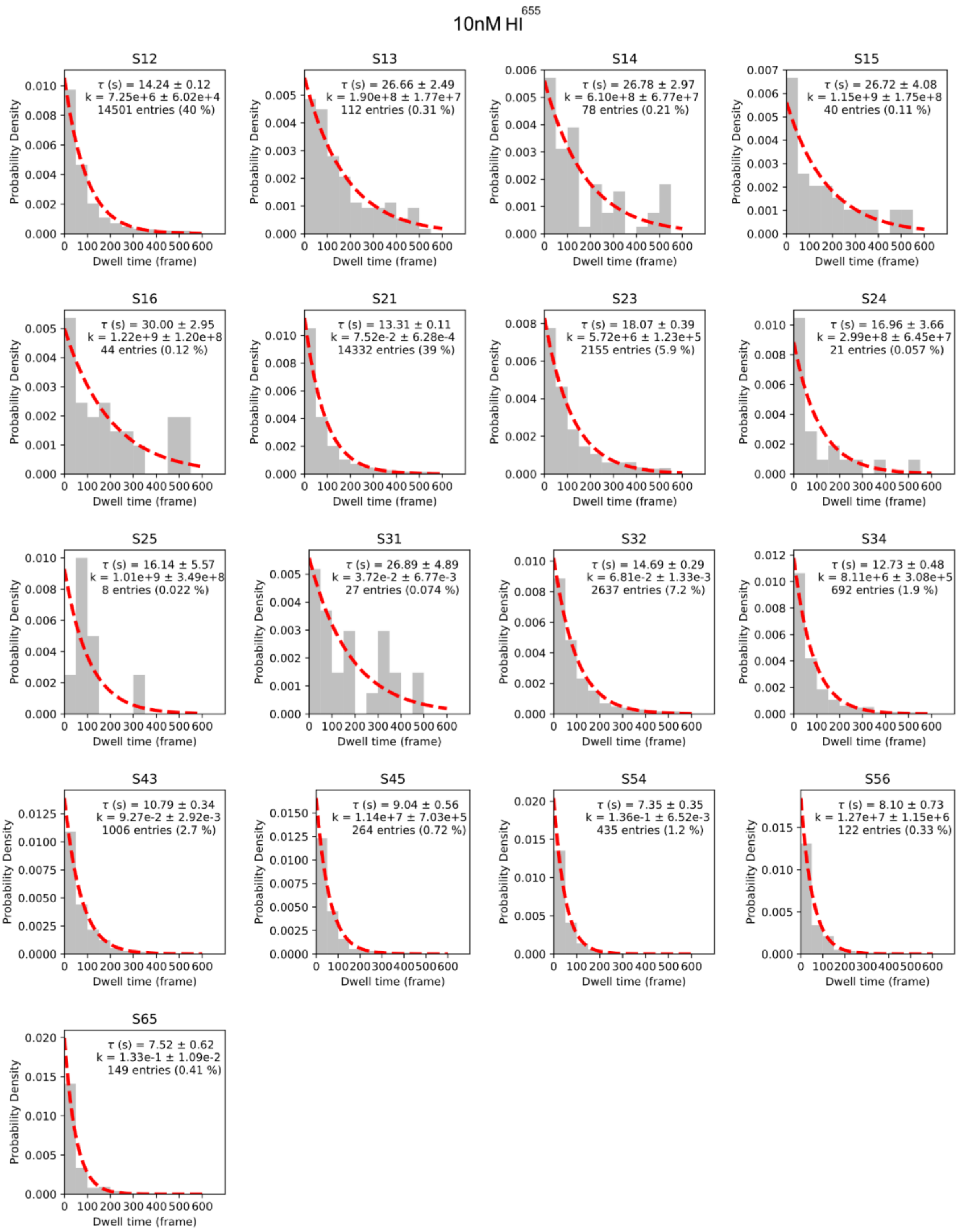


**Supplementary Figure 22:** Single exponential fit (red dotted line) to dwell times (gray) for 17 separable clusters of unique transitions for 10 nM $HI^{655}$. Within each fit is the extracted overall dwell time, rate constant and density (how many times the transition is observed) shown.

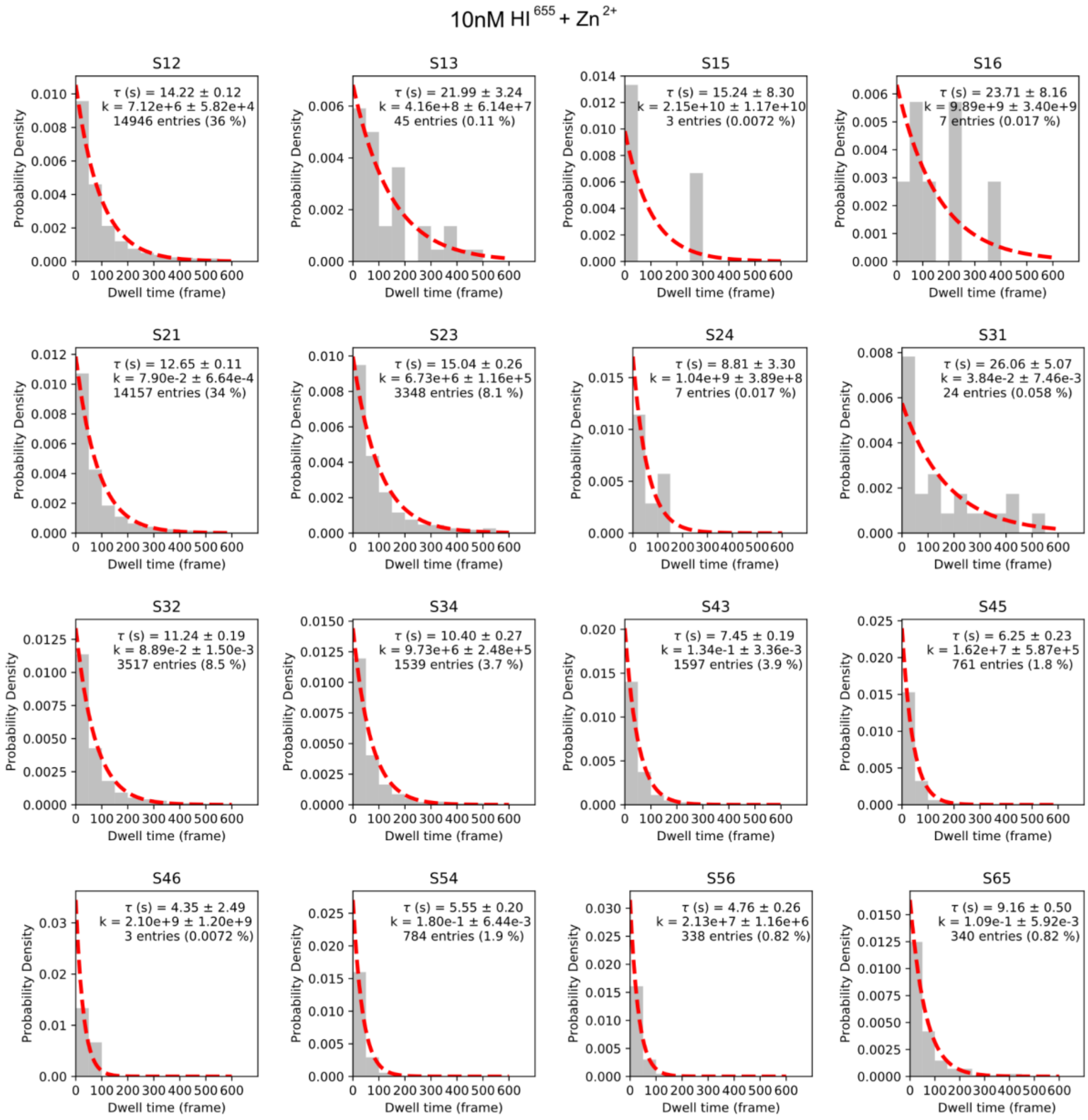


**Supplementary Figure 23:** Single exponential fit (red dotted line) to dwell times (gray) for 17 separable clusters of unique transitions for 10 nM $HI^{655}$ + 100 μM $Zn^{2+}$. Within each fit is the extracted overall dwell time, rate constant and density (how many times the transition is observed) shown.

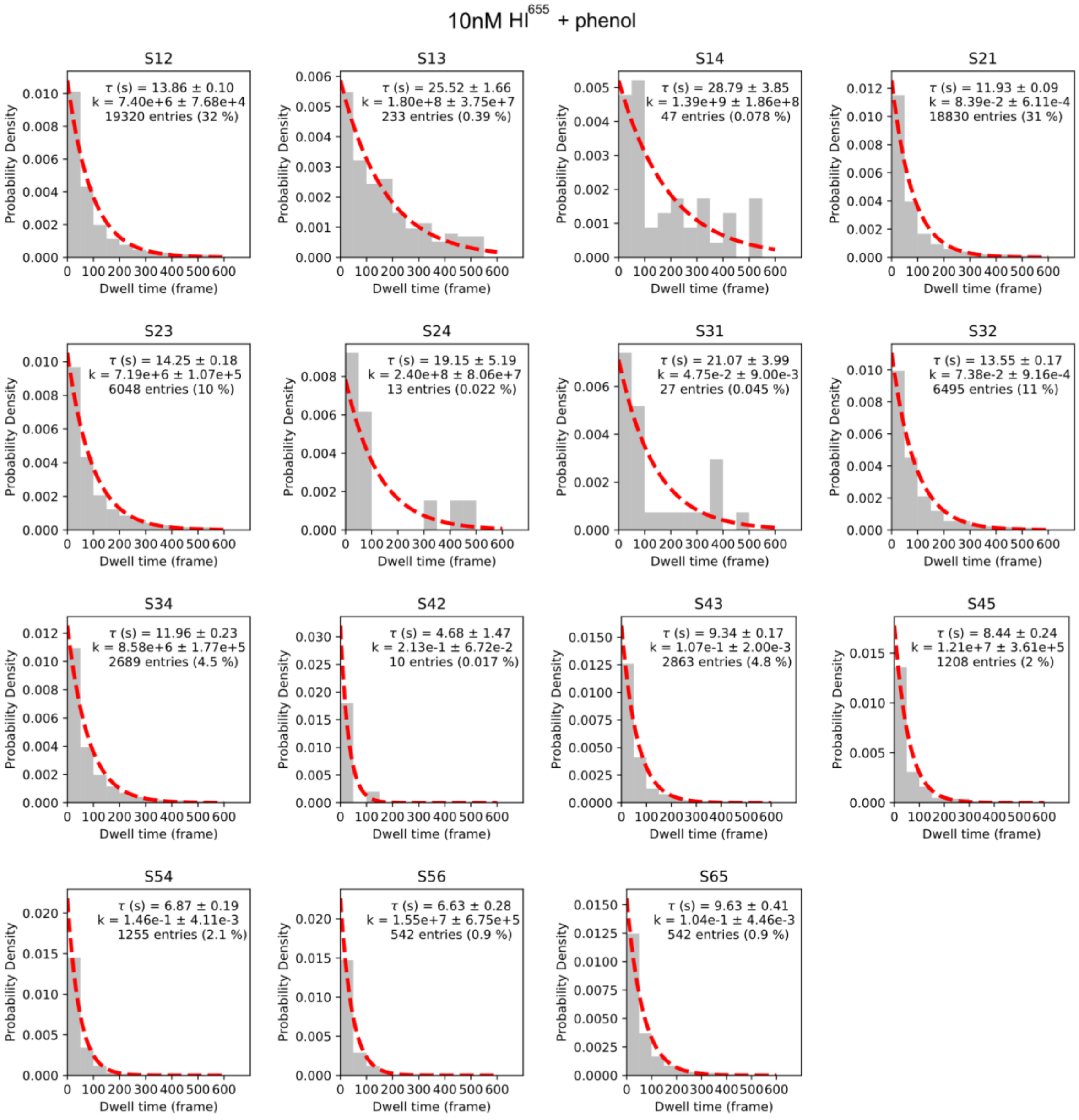


**Supplementary Figure 24:** Single exponential fit (red dotted line) to dwell times (gray) for 17 separable clusters of unique transitions for 10 nM $HI^{655}$ + 25 μM phenol. Within each fit is the extracted overall dwell time, rate constant and density (how many times the transition is observed) shown.

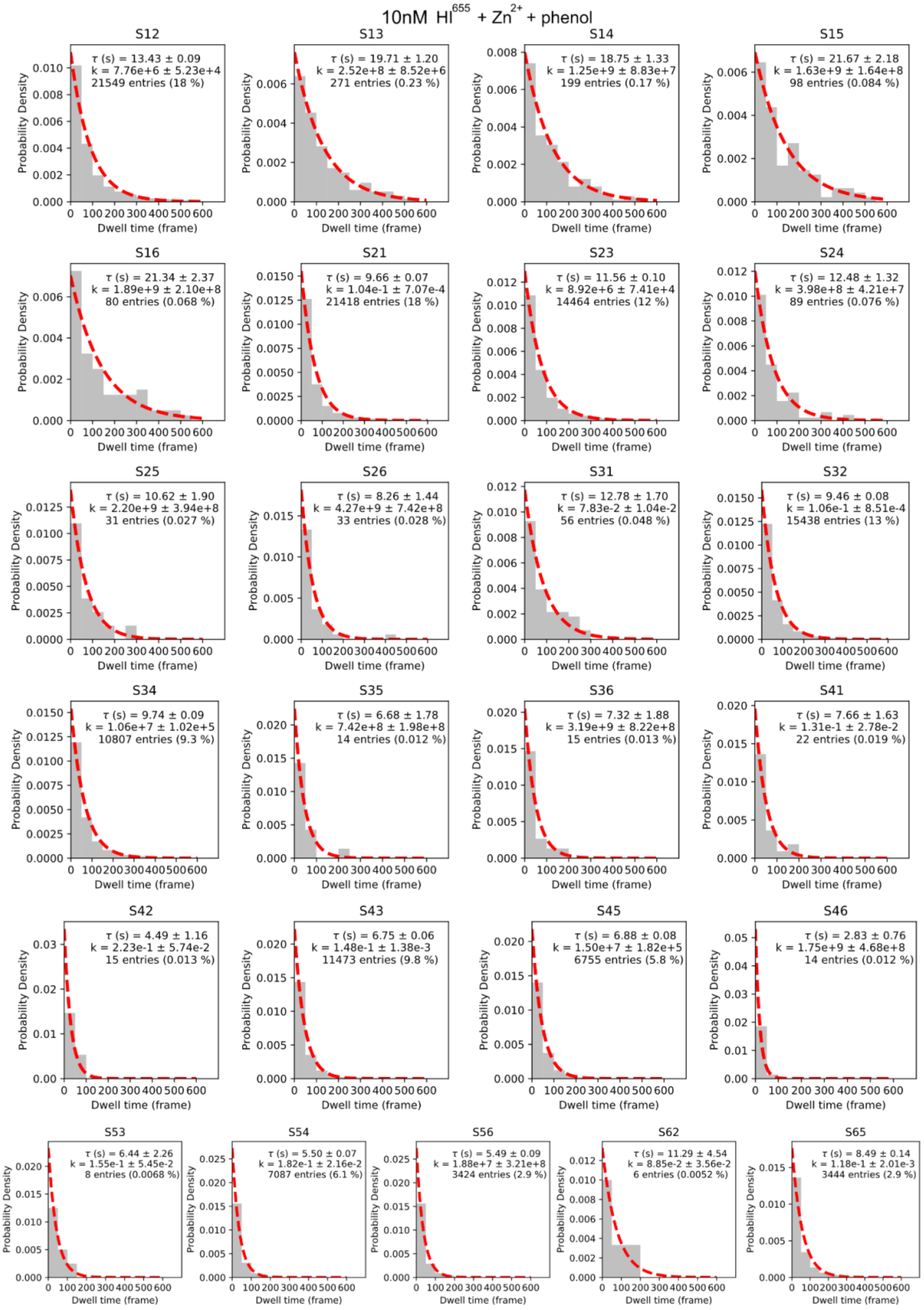


**Supplementary Figure 25:** Single exponential fit (red dotted line) to dwell times (gray) for 17 separable clusters of unique transitions for 10 nM $HI^{655}$ + 100 μM $Zn^{2+}$ + 25 μM phenol. Within each fit is the extracted overall dwell time, rate constant and density (how many times the transition is observed) shown.

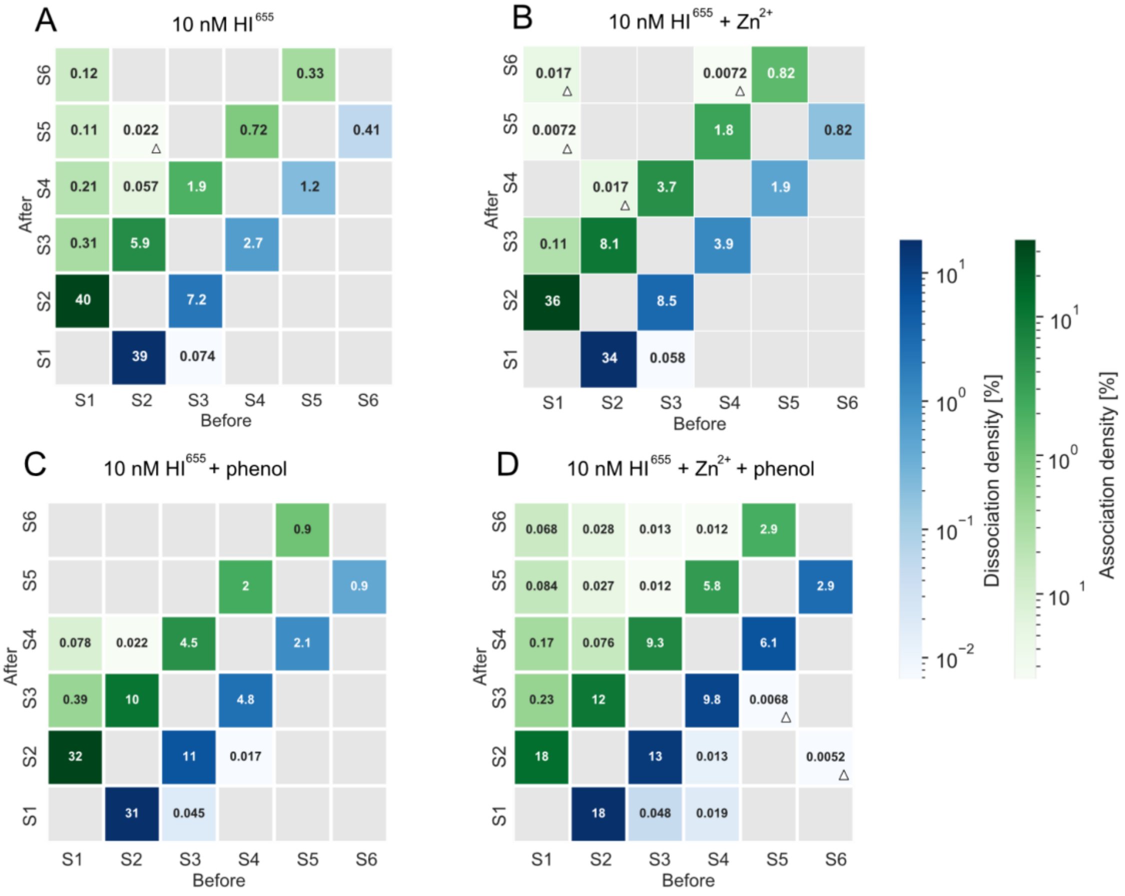


**Supplementary Figure 26:** CHESS (Complete HEatmap of State transitionS) plot shows thermodynamic parameters for each separate transition from a before state (x-axis) to an after state (y-axis). The numbers within the squares are the value for transition density. Gray squares are transitions with no data points. Transitions involving S0 (background) have been removed. Triangle denotes if less than 10 transitions were observed. Extracted association and dissociation transition densities for each transition for **(A)** 10 nM $HI^{655}$, **(B)** 10 nM $HI^{655}$ + 100 μM $Zn^{2+}$, **(C)**10 nM $HI^{655}$ + 25 μM phenol and **(D)** 10 nM $HI^{655}$ + 100 μM $Zn^{2+}$ + 25 μM phenol.

| 10 nM $HI^{655}$ | | | |
|---|---|---|---|
| Transition | $k_{association}$ [$s^{-1}$ $M^{-1}$] | $k_{dissociation}$ [$s^{-1}$] | $E_{act}$ [kJ/mol] |
| S12 | 7.25E+06 ± 6.02E+04 | | 3.38E+01 ± 2.06E-02 |
| S13 | 1.90E+08 ± 1.77E+07 | | 2.58E+01 ± 2.31E-01 |
| S14 | 6.10E+08 ± 6.77E+07 | | 2.29E+01 ± 2.75E-01 |
| S15 | 1.15E+09 ± 1.75E+08 | | 2.13E+01 ± 3.78E-01 |
| S16 | 1.22E+09 ± 1.20E+08 | | 2.12E+01 ± 2.44E-01 |
| S21 | | 7.52E-02 ± 6.28E-04 | 7.94E+01 ± 2.07E-02 |
| S23 | 5.72E+06 ± 1.23E+05 | | 3.44E+01 ± 5.34E-02 |
| S24 | 2.99E+08 ± 6.45E+07 | | 2.46E+01 ± 5.35E-01 |
| S25 | 1.01E+09 ± 3.49E+08 | | 2.16E+01 ± 8.55E-01 |
| S31 | | 3.72E-02 ± 6.77E-03 | 8.11E+01 ± 4.51E-01 |
| S32 | | 6.81E-02 ± 1.33E-03 | 7.96E+01 ± 4.82E-02 |
| S34 | 8.11E+06 ± 3.08E+05 | | 3.36E+01 ± 9.42E-02 |
| S43 | | 9.27E-02 ± 2.92E-03 | 7.89E+01 ± 7.81E-02 |
| S45 | 1.14E+07 ± 7.03E+05 | | 3.27E+01 ± 1.52E-01 |
| S54 | | 1.36E-01 ± 6.52E-03 | 7.79E+01 ± 1.19E-01 |
| S56 | 1.27E+07 ± 1.15E+06 | | 3.24E+01 ± 2.24E-01 |
| S65 | | 1.33E-01 ± 1.09E-02 | 7.80E+01 ± 2.03E-01 |

**Supplementary Table 12:** All extracted rate constants and corresponding activation energies for 10 nM $HI^{655}$. $k_{association}$ [$s^{-1}$ $M^{-1}$] is the association rate constant, while $k_{dissociation}$ [$s^{-1}$] is the dissociation rate constant. The activation energy $E_{act}$ [kJ/mol] is calculated as shown in eq. 3.

| 10 nM $HI^{655}$ + 100 µM $Zn^{2+}$ | | | |
|---|---|---|---|
| Transition | $k_{association}$ [$s^{-1}$ $M^{-1}$] | $k_{dissociation}$ [$s^{-1}$] | $E_{act}$ [kJ/mol] |
| S12 | 7.12E+06 ± 5.82E+04 | | 3.39E+01 ± 2.03E-02 |
| S13 | 4.16E+08 ± 6.14E+07 | | 2.38E+01 ± 3.65E-01 |
| S15 | 2.15E+10 ± 1.17E+10 | | 1.40E+01 ± 1.35E+00 |
| S16 | 9.89E+09 ± 3.40E+09 | | 1.60E+01 ± 8.53E-01 |
| S21 | | 7.90E-02 ± 6.64E-04 | 7.93E+01 ± 2.08E-02 |
| S23 | 6.73E+06 ± 1.16E+05 | | 3.40E+01 ± 4.28E-02 |
| S24 | 1.04E+09 ± 3.89E+08 | | 2.15E+01 ± 9.27E-01 |
| S31 | | 3.84E-02 ± 7.46E-03 | 8.11E+01 ± 4.82E-01 |
| S32 | | 8.89E-02 ± 1.50E-03 | 7.90E+01 ± 4.18E-02 |
| S34 | 9.73E+06 ± 2.48E+05 | | 3.31E+01 ± 6.32E-02 |
| S43 | | 1.34E-01 ± 3.36E-03 | 7.80E+01 ± 6.20E-02 |
| S45 | 1.62E+07 ± 5.87E+05 | | 3.19E+01 ± 8.98E-02 |
| S46 | 2.10E+09 ± 1.20E+09 | | 1.98E+01 ± 1.42E+00 |
| S54 | | 1.80E-01 ± 6.44E-03 | 7.72E+01 ± 8.85E-02 |
| S56 | 2.13E+07 ± 1.16E+06 | | 3.12E+01 ± 1.35E-01 |
| S65 | | 1.09E-01 ± 5.92E-03 | 7.85E+01 ± 1.34E-01 |

**Supplementary Table 13:** All extracted rate constants and corresponding activation energies for 10 nM $HI^{655}$ + 100 µM $Zn^{2+}$. $k_{association}$ [$s^{-1}$ $M^{-1}$] is the association rate constant, while $k_{dissociation}$ [$s^{-1}$] is the dissociation rate constant. The activation energy $E_{act}$ [kJ/mol] is calculated as shown in eq. 3.

| 10 nM $HI^{655}$ + 25 µM phenol | | | |
|---|---|---|---|
| Transition | $k_{association}$ [$s^{-1}$ $M^{-1}$] | $k_{dissociation}$ [$s^{-1}$] | $E_{act}$ [kJ/mol] |
| S12 | 7.40E+06 ± 7.68E+04 | | 3.38E+01 ± 2.57E-02 |
| S13 | 1.80E+08 ± 3.75E+07 | | 2.59E+01 ± 5.16E-01 |
| S14 | 1.39E+09 ± 1.86E+08 | | 2.08E+01 ± 3.31E-01 |
| S21 | | 8.39E-02 ± 6.11E-04 | 7.91E+01 ± 1.81E-02 |
| S23 | 7.19E+06 ± 1.07E+05 | | 3.39E+01 ± 3.69E-02 |
| S24 | 2.40E+08 ± 8.06E+07 | | 2.52E+01 ± 8.31E-01 |
| S31 | | 4.75E-02 ± 9.00E-03 | 8.05E+01 ± 4.70E-01 |
| S32 | | 7.38E-02 ± 9.16E-04 | 7.94E+01 ± 3.07E-02 |
| S34 | 8.58E+06 ± 1.77E+05 | | 3.34E+01 ± 5.13E-02 |
| S42 | | 2.13E-01 ± 6.72E-02 | 7.68E+01 ± 7.80E-01 |
| S43 | | 1.07E-01 ± 2.00E-03 | 7.85E+01 ± 4.63E-02 |
| S45 | 1.21E+07 ± 3.61E+05 | | 3.26E+01 ± 7.37E-02 |
| S54 | | 1.46E-01 ± 4.11E-03 | 7.78E+01 ± 6.99E-02 |
| S56 | 1.55E+07 ± 6.75E+05 | | 3.20E+01 ± 1.08E-01 |
| S65 | | 1.04E-01 ± 4.46E-03 | 7.86E+01 ± 1.06E-01 |

**Supplementary Table 14:** All extracted rate constants and corresponding activation energies for 10 nM $HI^{655}$ + 25 µM phenol. $k_{association}$ [$s^{-1}$ $M^{-1}$] is the association rate constant, while $k_{dissociation}$ [$s^{-1}$] is the dissociation rate constant. The activation energy $E_{act}$ [kJ/mol] is calculated as shown in eq. 3

| 10 nM $HI^{655}$ + 100 µM $Zn^{2+}$ + 25 µM phenol | | | |
|---|---|---|---|
| Transition | $k_{association}$ [$s^{-1}$ $M^{-1}$] | $k_{dissociation}$ [$s^{-1}$] | $E_{act}$ [kJ/mol] |
| S12 | 7.67E+06 ± 5.23E+04 | | 3.37E+01 ± 1.68E-02 |
| S13 | 2.52E+08 ± 8.52E+07 | | 2.51E+01 ± 8.38E-02 |
| S14 | 1.25E+09 ± 8.83E+07 | | 2.11E+01 ± 1.75E-01 |
| S15 | 1.63E+09 ± 1.64E+08 | | 2.04E+01 ± 2.49E-01 |
| S16 | 1.89E+09 ± 2.10E+08 | | 2.01E+01 ± 2.75E-01 |
| S21 | | 1.04E-01 ± 7.07E-04 | 7.86E+01 ± 1.69E-02 |
| S23 | 8.92E+06 ± 7.41E+04 | | 3.33E+01 ± 2.06E-02 |
| S24 | 3.98E+08 ± 4.21E+07 | | 2.39E+01 ± 2.62E-01 |
| S25 | 2.29E+09 ± 3.94E+08 | | 1.97E+01 ± 4.44E-01 |
| S26 | 4.27E+09 ± 7.42E+08 | | 1.80E+01 ± 4.31E-01 |
| S31 | | 7.83E-02 ± 1.04E-02 | 7.93E+01 ± 3.30E-01 |
| S32 | | 1.06E-02 ± 8.51E-04 | 7.86E+01 ± 1.99E-02 |
| S34 | 1.06E+06 ± 1.02E+05 | | 3.29E+01 ± 2.38E-02 |
| S35 | 7.42E+08 ± 1.98E+08 | | 2.24E+01 ± 6.59E-01 |
| S36 | 3.19E+09 ± 8.22E+08 | | 1.88E+01 ± 6.37E-01 |
| S41 | | 1.31E-01 ± 2.78E-02 | 7.80E+01 ± 5.27E-01 |
| S42 | | 2.23E-01 ± 5.74E-02 | 7.67E+01 ± 6.38E-01 |
| S43 | | 1.48E-01 ± 1.38E-03 | 7.77E+01 ± 2.31E-02 |
| S45 | 1.50E+07 ± 1.82E+05 | | 3.20E+01 ± 3.01E-02 |
| S46 | 1.75E+09 ± 1.68E+08 | | 2.02E+01 ± 6.61E-01 |
| S53 | | 1.55E-01 ± 5.45E-02 | 7.76E+01 ± 8.69E-01 |
| S54 | | 1.82E-01 ± 2.16E-03 | 7.72E+01 ± 2.94E-02 |

| | | | |
|---|---|---|---|
| S56 | 1.88E+07 ± 3.21E+05 | | 3.15E+01 ± 4.23E-02 |
| S62 | | 8.85E-02 ± 3.56E-02 | 7.89E+01 ± 9.96E-01 |
| S65 | | 1.18E-01 ± 2.01E-03 | 7.83E+01 ± 4.22E-02 |

**Supplementary Table 15:** All extracted rate constants and corresponding activation energies for 10 nM $HI^{655}$ + 100 µM $Zn^{2+}$ + 25 µM phenol. $k_{association}$ [$s^{-1}$ $M^{-1}$] is the association rate constant, while $k_{dissociation}$ [$s^{-1}$] is the dissociation rate constant. The activation energy $E_{act}$ [kJ/mol] is calculated as shown in eq. 3.

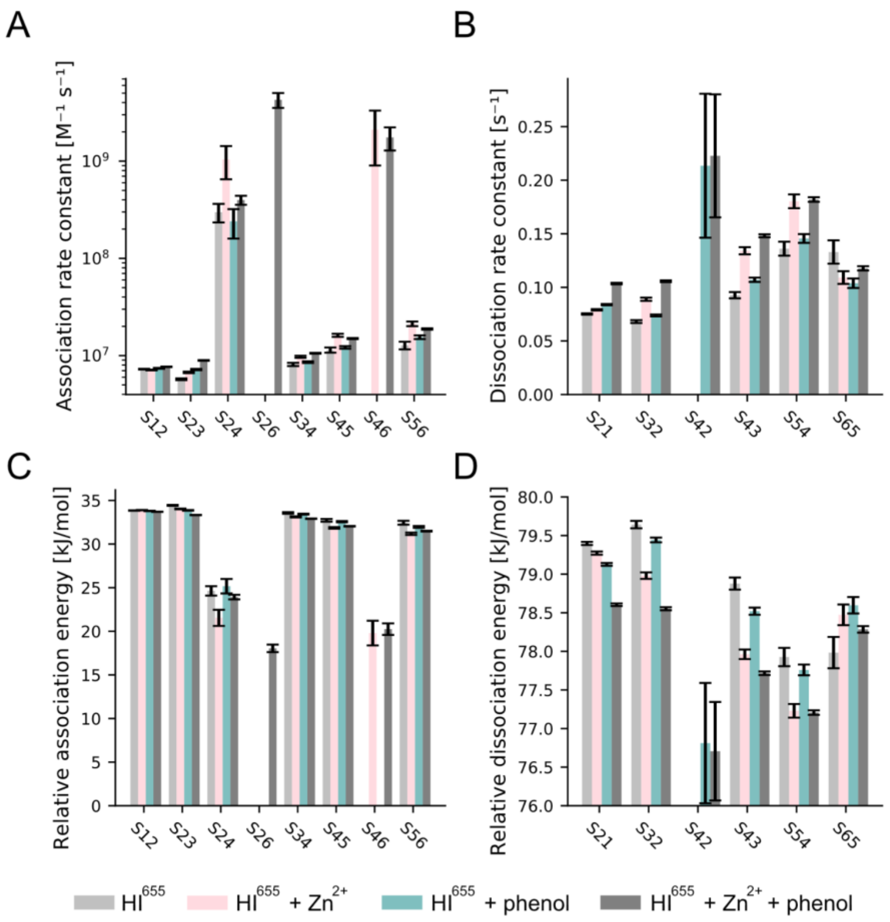


**Supplementary Figure 27:** Rate constants and relative activation energy for association and dissociation from HMM analysis. **(A)** Rate constants for monomer addition and dimer addition for 10 nM $HI^{655}$, 10 nM $HI^{655}$ + 25 µM phenol, 10 nM $HI^{655}$ + 100 µM $Zn^{2+}$ and 10 nM $HI^{655}$ + 100 µM $Zn^{2+}$ + 25 µM phenol. **(B)** Rate constants for monomer disassembly for 10 nM $HI^{655}$, 10 nM $HI^{655}$ + 25 µM phenol, 10 nM $HI^{655}$ + 100 µM $Zn^{2+}$ and 10 nM$HI^{655}$ + 100 µM $Zn^{2+}$ + 25 µM phenol. Error Bars are fit errors. **(C)** Activation energy for monomer addition and dimer addition for 10 nM $HI^{655}$, 10 nM $HI^{655}$ + 25 µM phenol, 10 nM $HI^{655}$ + 100 µM $Zn^{2+}$ and 10 nM $HI^{655}$ + 100 µM $Zn^{2+}$ + 25 µM phenol. **(D)**

Activation energy for monomer disassembly for 10 nM $HI^{655}$, 10 nM $HI^{655}$ + 25 μM phenol, 10 nM $HI^{655}$ + 100 μM $Zn^{2+}$ and 10 nM $HI^{655}$ + 100 μM $Zn^{2+}$ + 25 μM phenol. Error bars are fit errors.

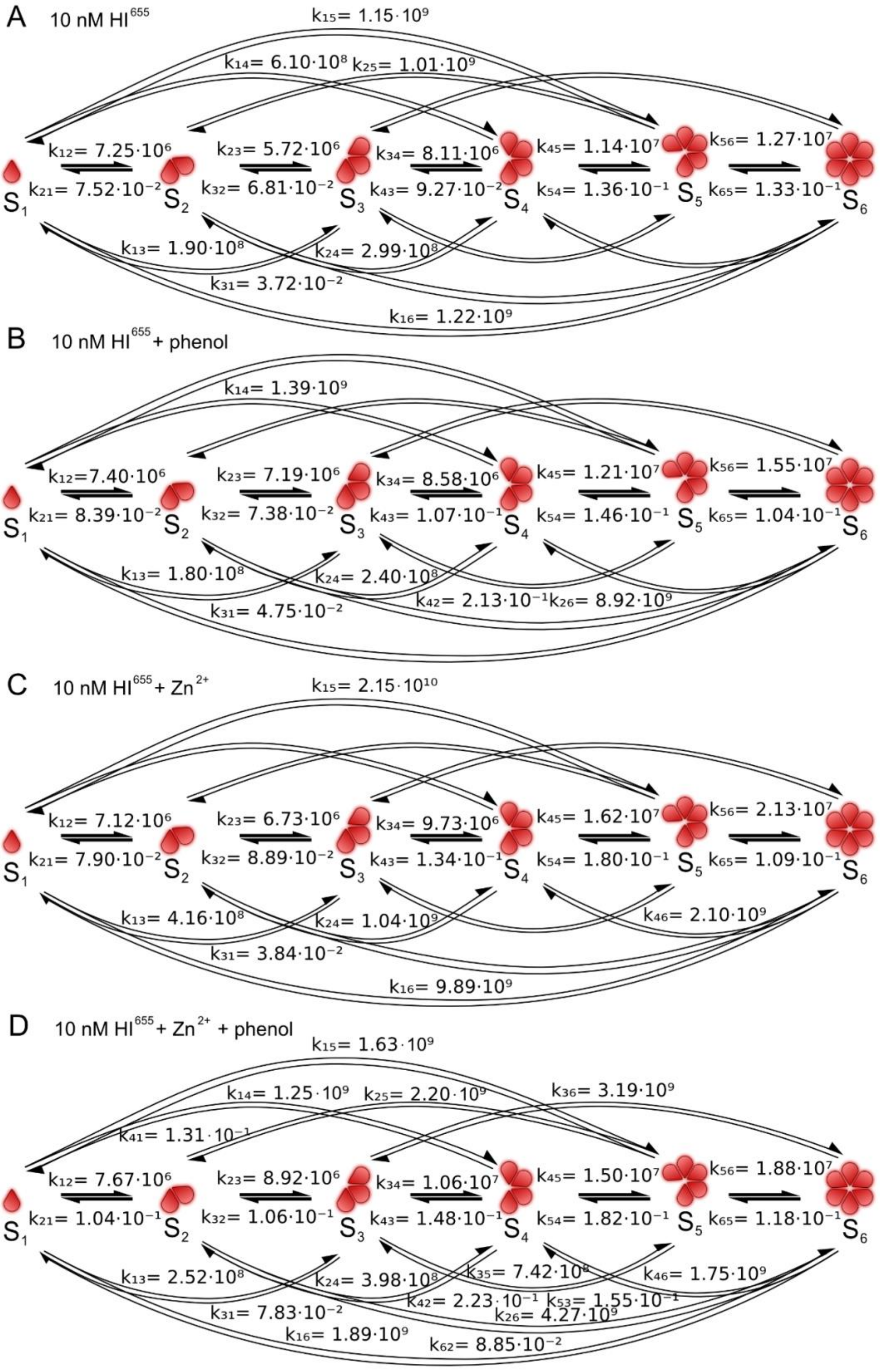


**Supplementary Figure 28:** Proposed mechanistic model for insulin oligomerization. Rate constants are obtained from Hidden Markov Model analysis for **(A)** 10 nM HI$^{655}$, **(B)** 10 nM HI$^{655}$ + 25 μM phenol, **(C)** 10 nM $HI^{655}$ + 100 µM $Zn^{2+}$ and (**D)** 10 nM $HI^{655}$ + 100 µM $Zn^{2+}$ + 25 µM phenol and plotted into a cartoon of the insulin oligomerization mechanism. The found rate constants suggest that oligomerization mostly happens via monomeric addition of insulin, while addition of $Zn^{2+}$ promotes dimeric insulin addition via monomer - dimer - tetramer - hexamer equilibrium and $Zn^{2+}$ and phenol reroutes the pathway to monomer - dimer - hexamer equilibrium.

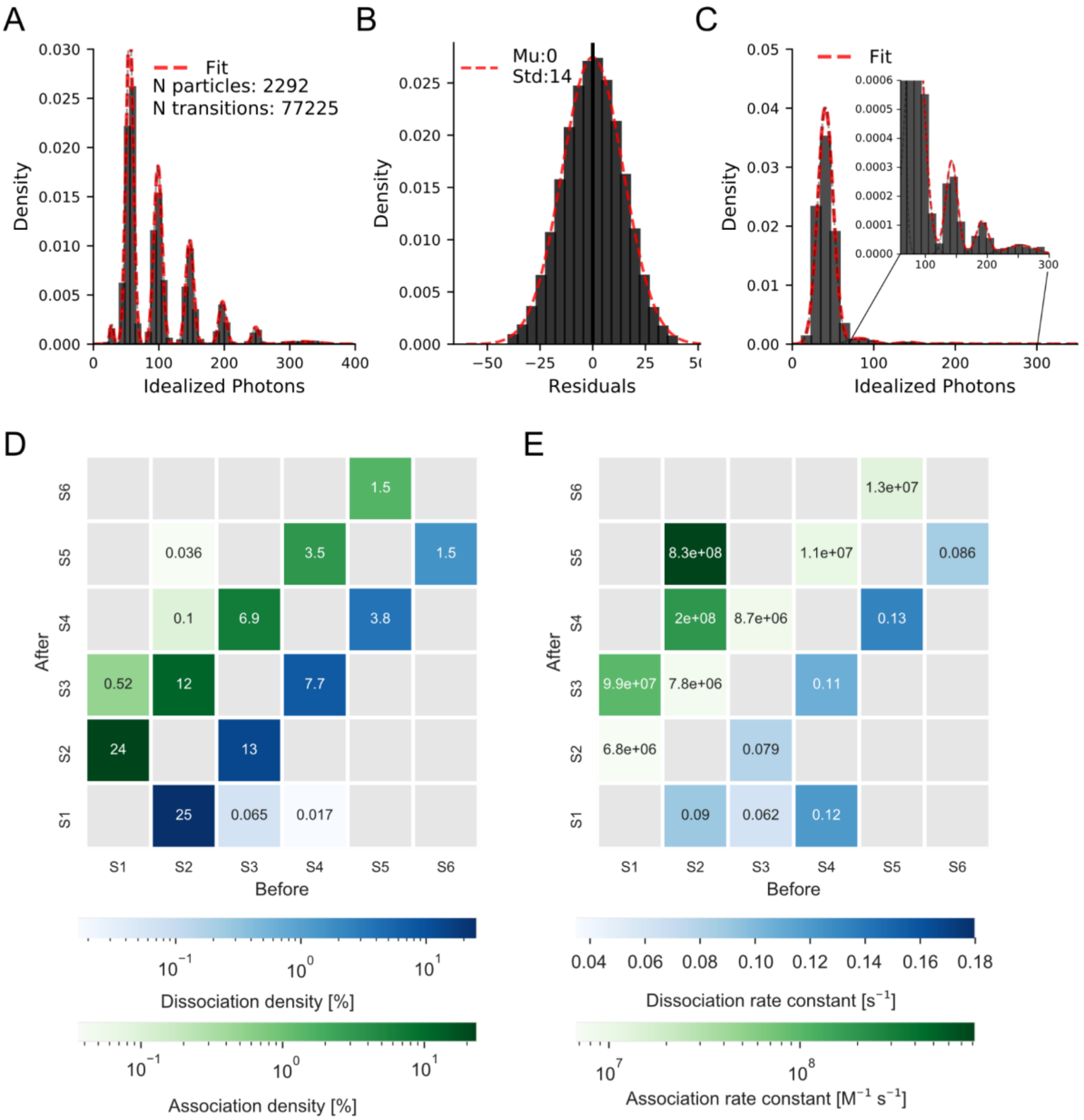


**Supplementary Figure 29:** Hidden Markov Modeling of 10 nM NovoRapid$^{655}$. **(A)** Idealized photon histogram with a Gaussian mixture model fit to the seven distinct distributions. **(B)** Residuals for experiments grouped in histogram and fitted with Gaussian. μ and σ are displayed and show no systematic error in HMM fit. **(C)** Estimation of oligomer concentration on solution. Histogram of stepheight (essentially difference between photon count after a transition and photon count before a transition). Distributions correspond to monomer addition, dimer addition, trimer addition, tetramer addition and pentamer addition. Histogram fitted with five Gaussian distributions to get weight for each distribution. **(D, E)** CHESS (Complete HEatmap of State transitionS) plot shows thermodynamic (D) and kinetic (E) parameters for each separate transition from a before state (x-axis) to an after state (y-axis). The numbers within the squares are the value for transition density. Gray squares are transitions with no data points. Transitions involving S0 (background) have been removed. Triangle denotes if less than 10 transitions were observed.

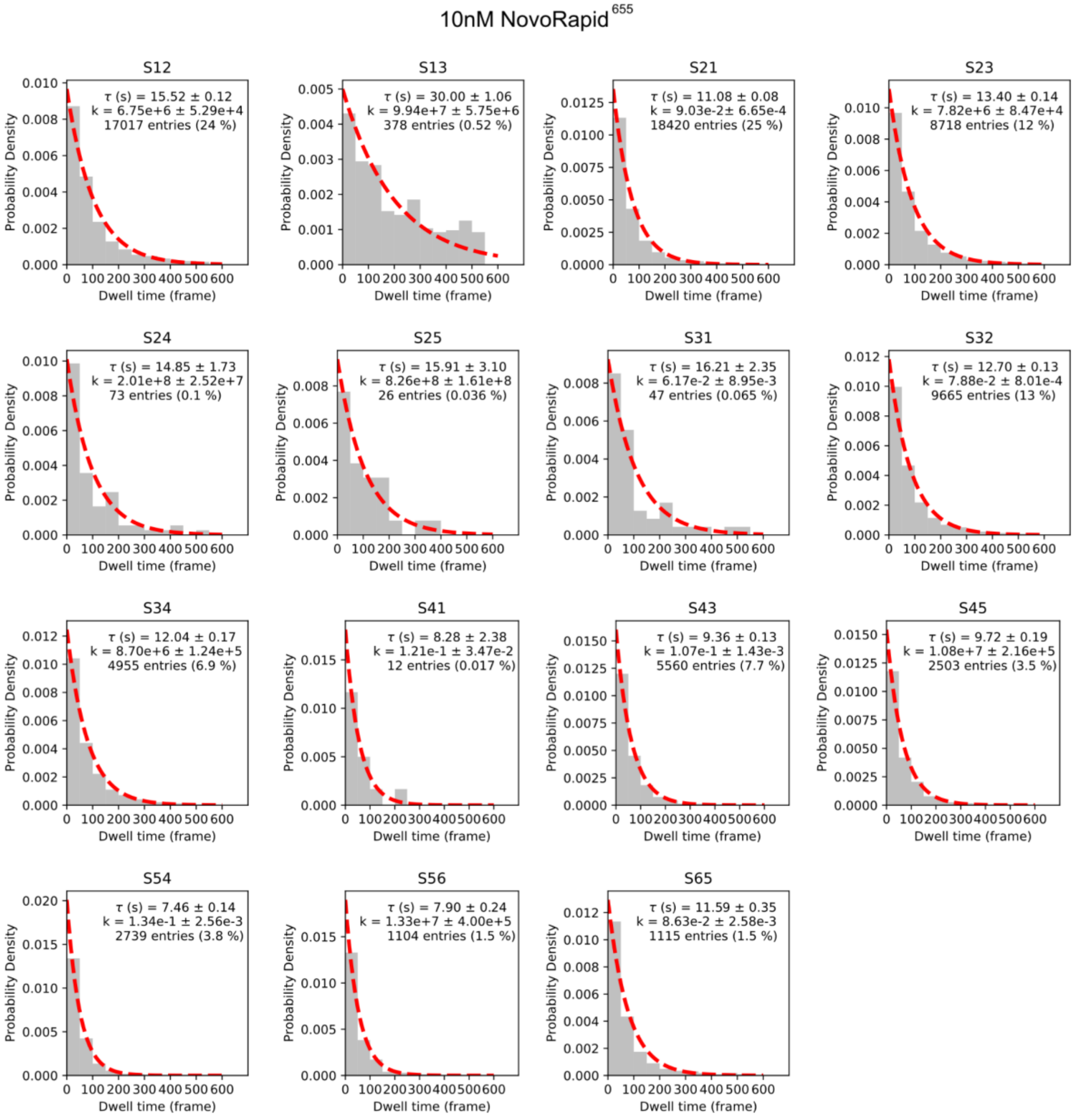


**Supplementary Figure 30:** Single exponential fit (red dotted line) to dwell times (gray) for 17 separable clusters of unique transitions for 10 nM NovoRapid$^{655}$. Within each fit is the extracted overall dwell time, rate constant and density (how many times the transition is observed) shown.

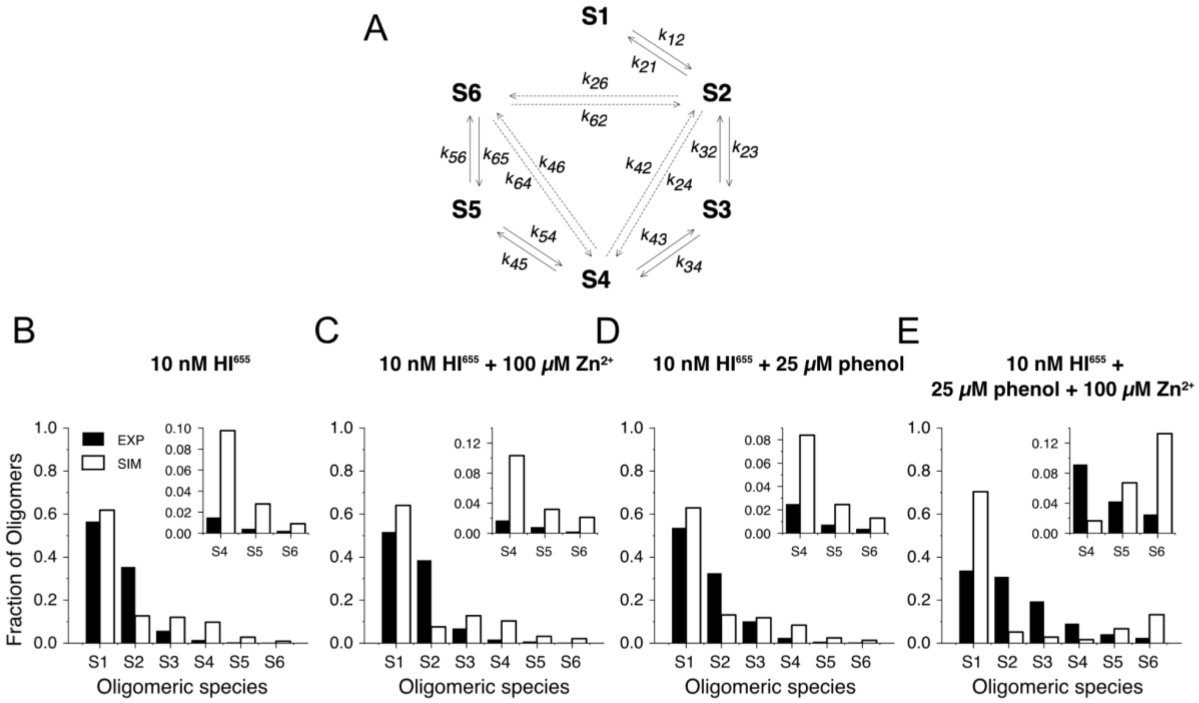


**Supplementary Figure 31:** Simulations **(A)** Model used for simulating time evolution of oligomeric species including monomer and dimer association and dissociation steps. Comparison of experimental (black bars) and experimentally (empty bars) found fraction of oligomers for **(B)** 10 nM $HI^{655}$, **(C)** 10 nM $HI^{655}$ + 100 μM $Zn^{2+}$, **(D)** 10 nM $HI^{655}$ + 25 μM phenol and **(E)** 10 nM $HI^{655}$ + phenol + $Zn^{2+}$. Simulated oligomeric fractions have no associated error. Error bars representing the standard error from the experimental oligomeric fractions are smaller than the lines.

| 10 nM $HI^{655}$ | | |
|---|---|---|
| Transition | $k_{association}$ [$s^{-1}$ $M^{-1}$] | $k_{dissociation}$ [$s^{-1}$] |
| S12 | 7.25E+06 ± 6.02E+04 | |
| S21 | | 7.52E-02 ± 6.28E-04 |
| S23 | 5.72E+06 ± 1.23E+05 | |
| S32 | | 6.81E-02 ± 1.33E-03 |
| S34 | 8.11E+06 ± 3.08E+05 | |
| S43 | | 9.27E-02 ± 2.92E-03 |
| S45 | 1.14E+07 ± 7.03E+05 | |
| S54 | | 1.36E-01 ± 6.52E-03 |
| S56 | 1.27E+07 ± 1.15E+06 | |
| S65 | | 1.33E-01 ± 1.09E-02 |
| S24 | 2.99E+08 ± 6.45E+07 | |
| S42 | | 2.13E-01 ± 6.72E-02 |
| S26 | 0.00E ± 0.00 | |
| S62 | | 0.00E ± 0.00 |
| S46 | 0.00E ± 0.00 | |
| S64 | | 0.01 |

**Supplementary Table 16:** Rate constants for dissociation and association used for the simulations with $HI^{655}$. Values in black are experimentally found as described earlier. Blue value is the experimentally found value for $HI^{655}$ + 25 μM phenol, as this dissociation rate constant was unobtainable from $HI^{655}$ experiments. S64 (red) was set as an arbitrary upper limit for the dissociation of hexamer into tetramer.

| 10 nM $HI^{655}$+ 100 µM $Zn^{2+}$ | | |
|---|---|---|
| Transition | $k_{association}$ [$s^{-1}$ $M^{-1}$] | $k_{dissociation}$ [$s^{-1}$] |
| S12 | 7.12E+06 ± 5.82E+04 | |
| S21 | | 7.90E-02 ± 6.64E-04 |
| S23 | 6.73E+06 ± 1.16E+05 | |
| S32 | | 8.89E-02 ± 1.50E-03 |
| S34 | 9.73E+06 ± 2.48E+05 | |
| S43 | | 1.34E-01 ± 3.36E-03 |
| S45 | 1.62E+07 ± 5.87E+05 | |
| S54 | | 1.80E-01 ± 6.44E-03 |
| S56 | 2.13E+07 ± 1.16E+06 | |
| S65 | | 1.09E-01 ± 5.92E-03 |
| S24 | 1.04E+09 ± 3.89E+08 | |
| S42 | | 2.13E-01 ± 6.72E-02 |
| S26 | 0.00E ± 0.00 | |
| S62 | | 0.00E ± 0.00 |
| S46 | 2.10E+09 ± 1.20E+09 | |
| S64 | | 0.01 |

**Supplementary Table 17:** Rate constants for dissociation and association used for the simulations with $HI^{655}$ + 100 µM $Zn^{2+}$. Values in black are experimentally found as described earlier. Blue value is the experimentally found value for $HI^{655}$ + 25 µM phenol, as this dissociation rate constant was unobtainable from $HI^{655}$ experiments. S64 (red) was set as an arbitrary upper limit for the dissociation of hexamer into tetramer.

| 10 nM $HI^{655}$ + 25 µM phenol | | |
|---|---|---|
| Transition | $k_{association}$ [$s^{-1}$ $M^{-1}$] | $k_{dissociation}$ [$s^{-1}$] |
| S12 | 7.40E+06 ± 7.68E+04 | |
| S21 | | 8.39E-02 ± 6.11E-04 |
| S23 | 7.19E+06 ± 1.07E+05 | |
| S32 | | 7.38E-02 ± 9.16E-04 |
| S34 | 8.58E+06 ± 1.77E+05 | |
| S43 | | 1.07E-01 ± 2.00E-03 |
| S45 | 1.21E+07 ± 3.61E+05 | |
| S54 | | 1.46E-01 ± 4.11E-03 |
| S56 | 1.55E+07 ± 6.75E+05 | |
| S65 | | 1.04E-01 ± 4.46E-03 |
| S24 | 2.40E+08 ± 8.06E+07 | |
| S42 | | 2.13E-01 ± 6.72E-02 |
| S26 | 0.00E ± 0.00 | |
| S62 | | 0.00E ± 0.00 |
| S46 | 0.00E ± 0.00 | |
| S64 | | 0.01 |

**Supplementary Table 18:** Rate constants for dissociation and association used for the simulations with $HI^{655}$ + 25 µM phenol. Values in black are experimentally found as described earlier. S64 (red) was set as an arbitrary upper limit for the dissociation of hexamer into tetramer.

| 10 nM $HI^{655}$ + 25 µM phenol + 100 µM $Zn^{2+}$ | | |
|---|---|---|
| Transition | $k_{association}$ [$s^{-1}$ $M^{-1}$] | $k_{dissociation}$ [$s^{-1}$] |
| S12 | 7.67E+06 ± 5.23E+04 | |
| S21 | | 1.04E-01 ± 7.07E-04 |
| S23 | 8.92E+06 ± 7.41E+04 | |
| S32 | | 1.06E-02 ± 8.51E-04 |
| S34 | 1.06E+07 ± 1.02E+05 | |
| S43 | | 1.48E-01 ± 1.38E-03 |
| S45 | 1.50E+07 ± 1.82E+05 | |
| S54 | | 1.82E-01 ± 2.16E-03 |
| S56 | 1.88E+07 ± 3.21E+05 | |
| S65 | | 1.18E-01 ± 2.01E-03 |
| S24 | 3.98E+08 ± 4.21E+07 | |
| S42 | | 2.23E-01 ± 5.74E-02 |
| S26 | 4.27E+09 ± 7.42E+08 | |
| S62 | | 8.85E-02 ± 3.56E-02 |
| S46 | 1.75E+09 ± 4.68E+08 | |
| S64 | | 0.01 |

**Supplementary Table 19:** Rate constants for dissociation and association used for the simulations with $HI^{655}$ + 25 µM phenol + 100 µM $Zn^{2+}$ . Values in black are experimentally found as described earlier. S64 (red) was set as an arbitrary upper limit for the dissociation of hexamer into tetramer.

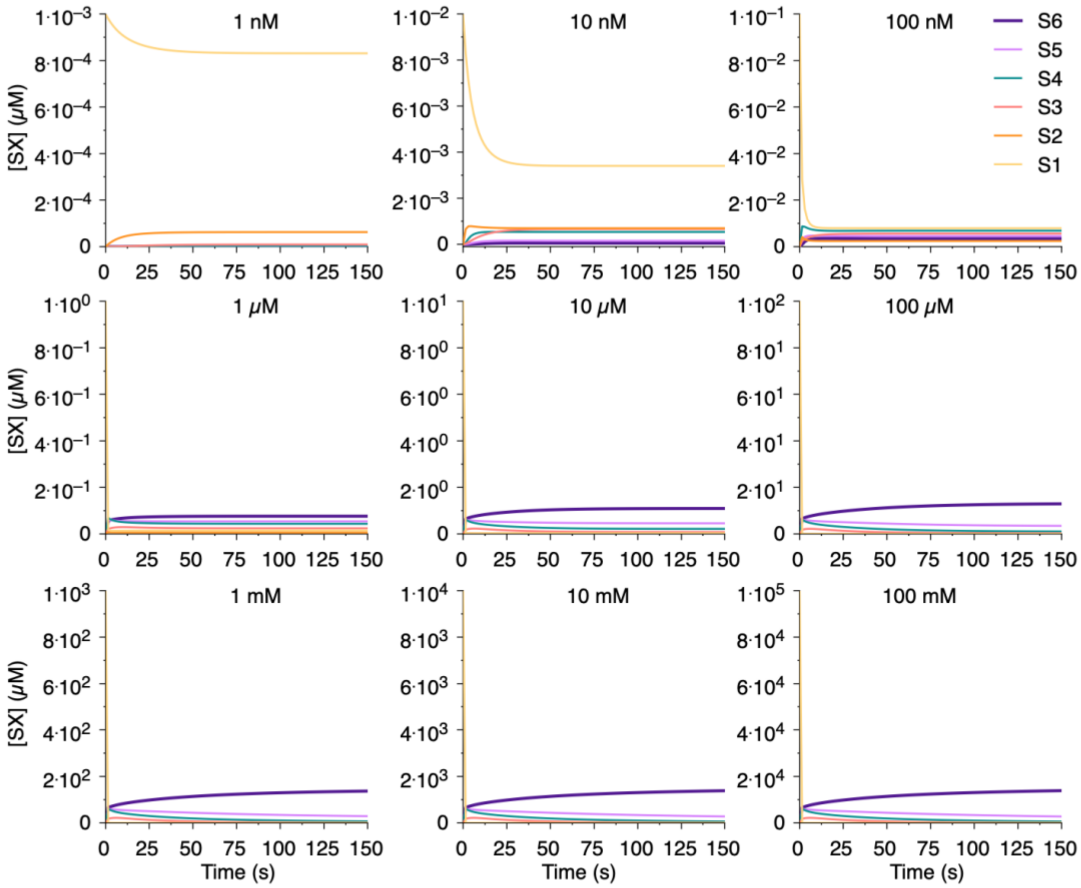


**Supplementary Figure 32:** Simulations for HI[655] with insulin concentration ranging from 1 nM to 100 mM.

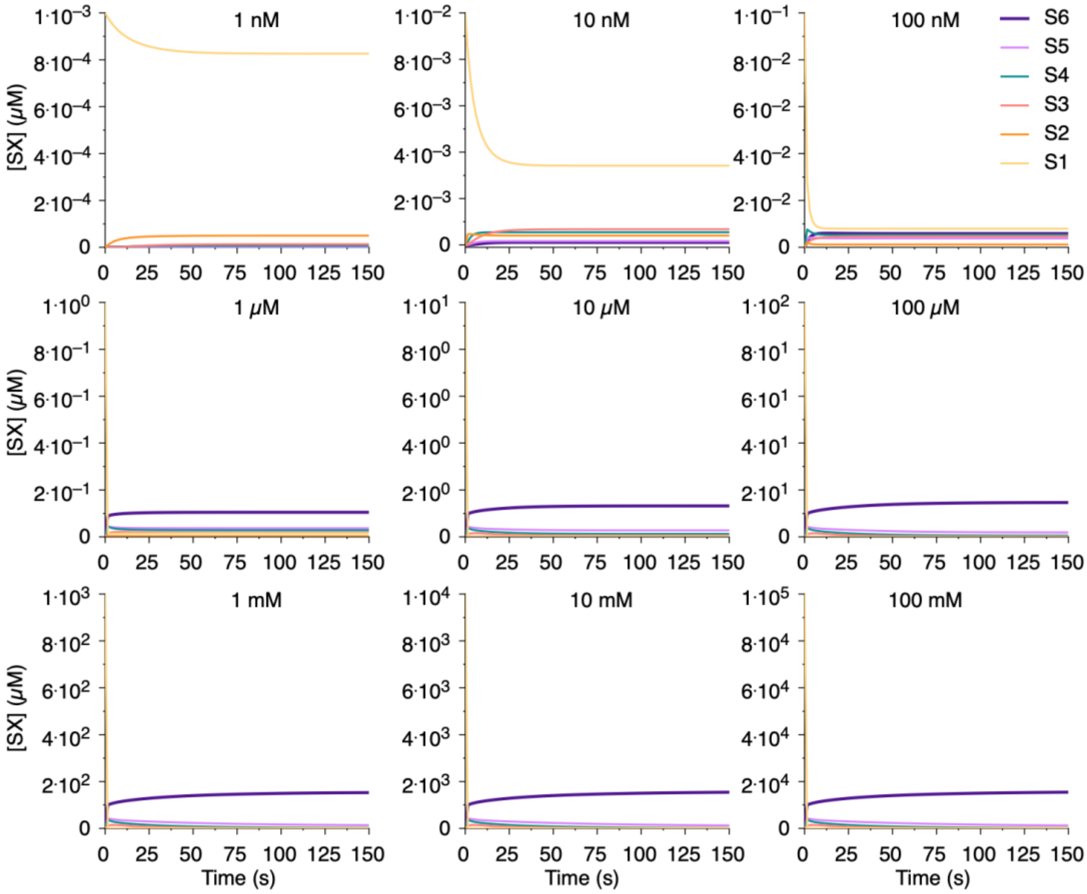


**Supplementary Figure 33:** Simulations for $HI^{655}$ + 100 μM $Zn^{2+}$ with insulin concentration ranging from 1 nM to 100 mM.

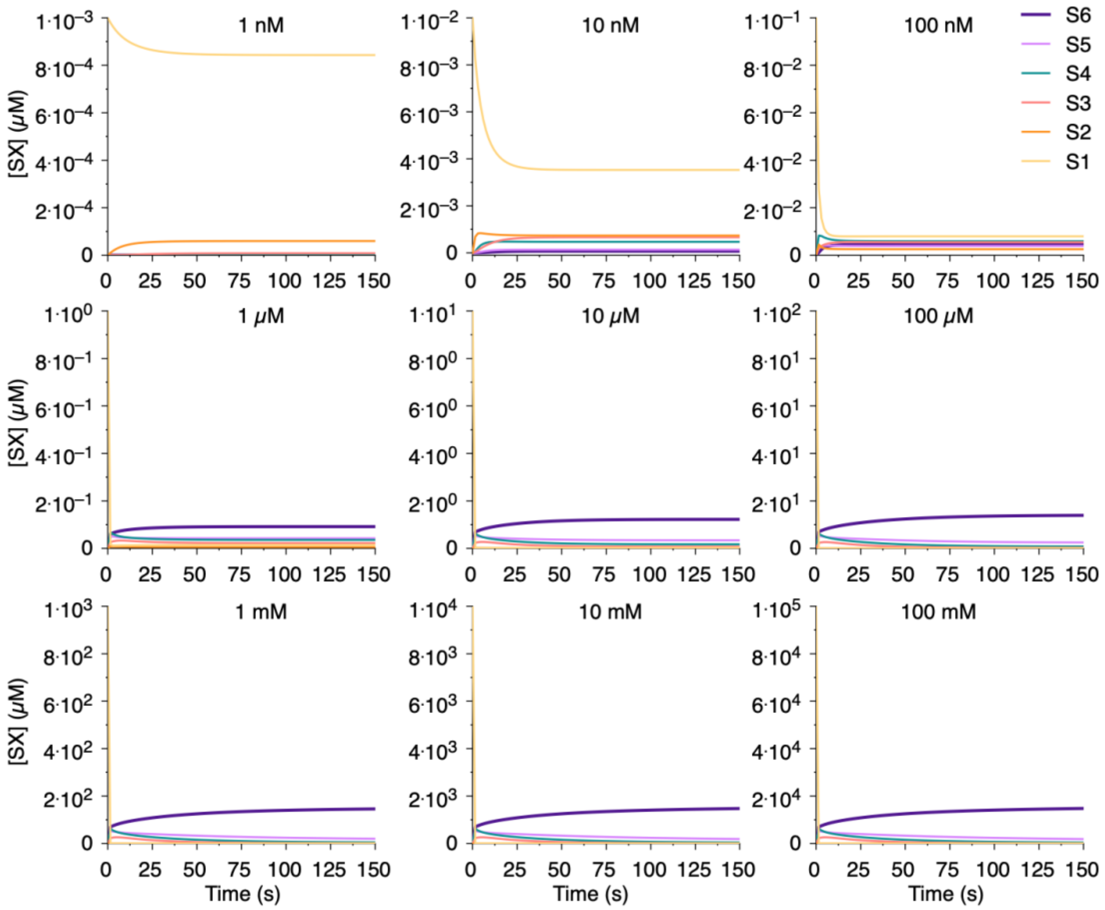


**Supplementary Figure 34:** Simulations for HI$^{655}$ + 25 μM phenol with insulin concentration ranging from 1 nM to 100 mM.

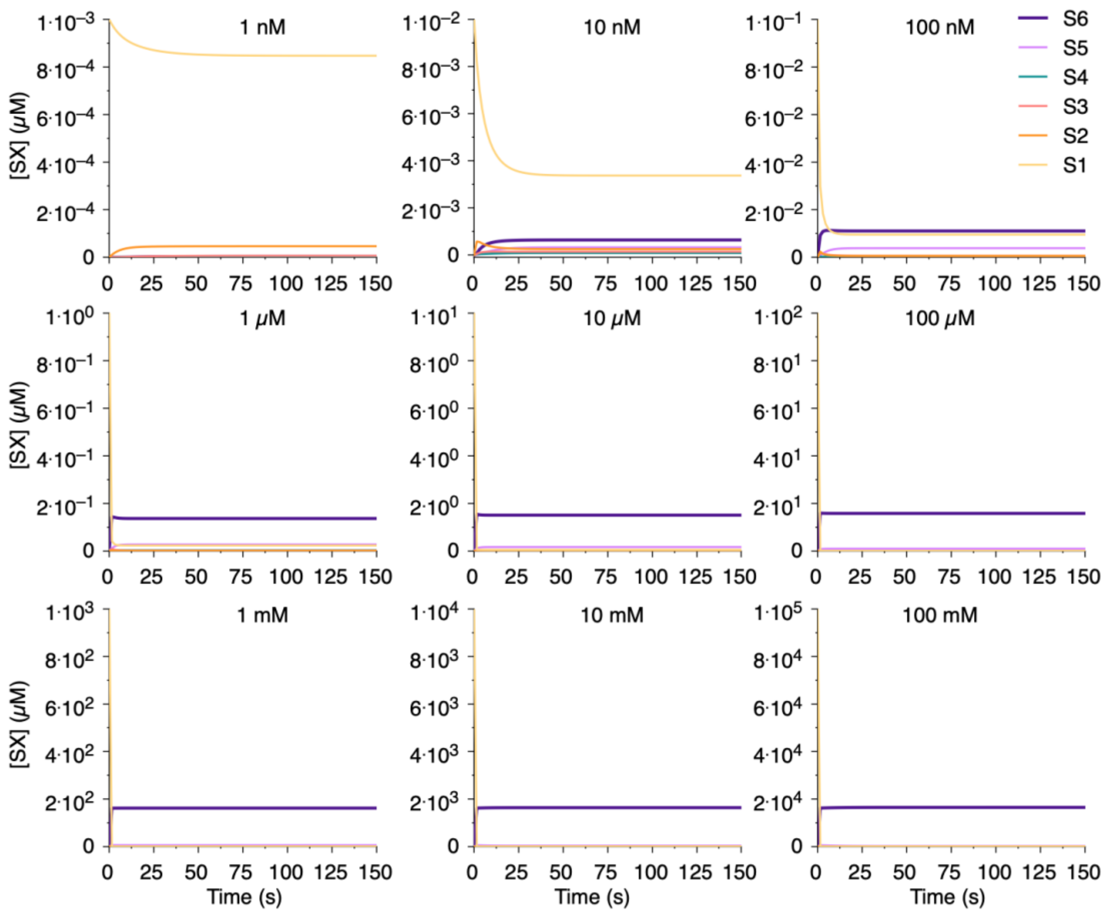


**Supplementary Figure 35:** Simulations for $HI^{655}$ + 25 μM phenol + 100 μM $Zn^{2+}$ with insulin concentration ranging from 1 nM to 100 mM.

| | $HI^{655}$ | $HI^{655}$ + $Zn^{2+}$ | $HI^{655}$ + phenol | $HI^{655}$ + $Zn^{2+}$ + phenol |
|---|---|---|---|---|
| $n_h$ | 0.54 ± 0.04 | 0.59 ± 0.05 | 0.57 ± 0.04 | 0.59 ± 0.04 |
| K | 2.61 ± 0.40 | 0.87 ± 0.14 | 1.50 ± 0.25 | 0.18 ± 0.02 |
| $B_{max}$ | 0.86 ± 0.01 | 0.92 ± 0.02 | 0.90 ± 0.02 | 0.97 ± 0.01 |

**Supplementary Table 20:** Fitting parameters from fitting fraction of hexameric oligomers with a Hill equation.
